# Evolutionary integration of the geography and pacing of the annual cycle in migratory birds

**DOI:** 10.1101/2024.03.08.584102

**Authors:** Benjamin M. Winger, Frank A. La Sorte, Matthew D. Hack, Teresa M. Pegan

**Author notes:** Department of Ecology and Evolutionary Biology, Yale University, New Haven, Connecticut, United States of America (Frank A. La Sorte). Center for Biodiversity and Global Change, Yale University, New Haven, Connecticut, United States of America (Frank A. La Sorte).

## Abstract

In migratory species, the temporal phases of the annual cycle are intrinsically linked to seasonally shifting geographic ranges. Despite intense interest in the annual cycle ecology of migration, a synthetic understanding of the relationship between the biogeography and phenology of seasonal migration remains elusive. Here, we interrogate the spatiotemporal structure of the annual cycle in a novel phylogenetic comparative framework. We use eBird, a massive avian occurrence dataset, to demarcate and measure in a consistent manner among species the portions of the annual cycle when a geographic distribution is stationary versus dynamic due to migration. Through comparative analyses of the durations of annual cycle stages for 150 species of migratory birds breeding in North America, we show that the duration of the migratory periods is remarkably consistent among species and is unrelated to the distance between breeding and nonbreeding locations. In other words, the seasonal distributions of long-distance migrants shift between their geographically distant stationary phases in the same amount of time as short-distance migrants, suggesting that individuals of long-distance migratory species have more synchronous periods of migration and likely a faster individual migratory pace than short-distance migrants. Our results further show that the amount of time a species spends on the breeding grounds is strongly inversely related to time spent on the nonbreeding grounds, revealing the length of the breeding versus nonbreeding stationary period to be the primary source of species-level variation in the pacing of the annual cycle, as opposed to the time needed for the migratory period. Further, our study reveals that the amount of time spent annually on the breeding versus nonbreeding grounds predicts the distance between breeding and nonbreeding locations, demonstrating key linkages between the biogeography of the migratory cycle, its phenology, and the evolution of life history tradeoffs.

## Introduction

The processes that sustain life and allow individuals to pass on their genes rarely occur evenly throughout the year. Rather, organisms tend to have distinct periods of the annual cycle dedicated to reproduction, with the remaining stages devoted to survival [1–5]. The balance of energy devoted to reproduction versus survival—both throughout the annual cycle and across the lifetime of individuals—characterizes much of a species’ life history strategy [6,7]. In migratory species, not only do individuals have distinct times of year when they reproduce, but they also reproduce in a location that is geographically and often ecologically distinct from where they spend the rest of the year [8,9]. Thus, their temporal annual cycles are linked to their biogeographic annual cycles. The annual cycle dynamics of migratory populations have been the focus of much research, for example by tracking individual movements throughout the year [10–14], modelling population regulation across seasons [15,16] or studying annual hormonal cycles [17–19]. Diagrams of the annual cycle illustrating the phenology of migratory and reproductive phases are commonplace in scientific literature [1,17,20]. Yet, when considering the great diversity of seasonally varying geographic distributions of migratory species and populations, we have a limited understanding of how the timing and duration of annual cycle stages relates to the complex seasonal biogeography of migration. Why do some species spend longer on their breeding or nonbreeding grounds than their sympatric close relatives? Why do some species migrate short distances, whereas others travel much further between distant breeding and nonbreeding locations?

Here, we interrogate the relationship between the biogeographic distributions of migratory bird species and the pacing and phenology of their migrations by developing a novel comparative approach that uses bird occurrence data to characterize the spatiotemporal structure of the annual cycle. We use this approach to test ecological and evolutionary factors that govern where and how long migratory species occupy the different parts of their seasonally varying geographic distributions. Specifically, we used occurrence data from the eBird citizen-science program [21] to estimate the geographic centroids of the daily distributions of 150 species of passerine birds (Table S1, *Materials and Methods*) across the annual cycle (Fig. 1, Fig. S3., Table S1). These species undergo seasonal migration along a latitudinal axis between breeding grounds in North America occupied during the northern hemisphere summer and nonbreeding grounds at lower latitudes occupied during the northern hemisphere winter (Fig. 1), and thus the geographic centroids of their distributions shift latitudinally throughout the year. By fitting curves to the latitudes of daily centroids and calculating their inflection points, we demarcated and quantified the durations of the four major geographic stages of the annual cycle for each species: the breeding stationary period and nonbreeding stationary period and the spring (pre-breeding) and fall (post-breeding) migratory periods (Fig. 1). The breeding stationary period encompasses reproduction and development, as all species in the study attain adult body size on or near the summer breeding grounds before their first fall migration. Our delineation of the breeding stationary period also typically includes the energy-intensive annual post-breeding feather molt (see Supplementary Text; [22–25]).

**Fig 1.**
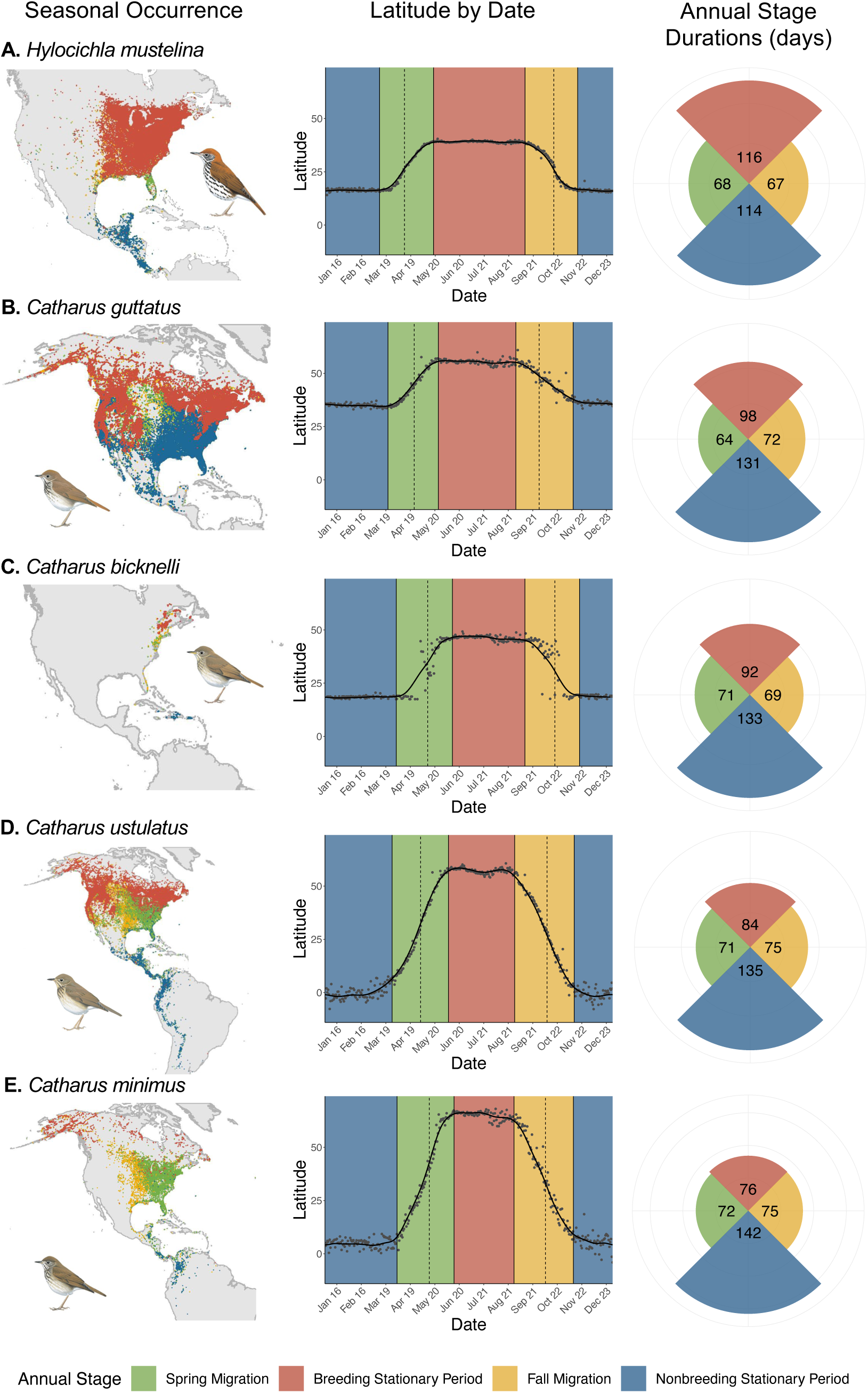
Geography and phenology of the annual cycle. Annual cycle demarcation for example species, (A) the Wood Thrush (*Hylocichla mustelina)* and (B-E) four close relatives in the genus *Catharus*. For these and the other species in the study (Table S1, Fig. S3.), reproduction occurs at higher latitudes during the northern hemisphere summer whereas the nonbreeding stationary period occurs at lower latitudes during the northern winter. Spring and fall migration occur in distinct periods of dynamic geographic movement. The species are ordered top to bottom according to their migratory distance (the geographic separation between breeding and nonbreeding ranges). A fifth species of migratory *Catharus (C. fuscescens*) was excluded from the study (see *Materials and Methods*). *Left column*: Distributional records used in the study from the eBird citizen science database [21]. *Center column*: Latitudes from daily geographic centroids (gray points, one point per day) derived from 20 years of eBird data. The smoothed black line depicts the average of 37 generalized additive model (GAM) fits of latitude by day of year (see *Materials and Methods*). Dashed vertical lines indicate the maximum (spring, green) and minimum (fall, gold) first derivatives of GAM fits, signifying the fastest change in latitude over time and corresponding to peak northbound (spring) and southbound (fall) migration. Solid vertical lines indicate the maximum and minimum second derivatives of GAM fits and demarcate the beginning and end of the spring and fall migratory periods. The first and second derivative (dashed and solid vertical lines) were drawn from the distribution of derivatives from 37 GAM fits (*k* = 4-40, see *Materials and Methods*). *Right column*: Pie chart showing the durations in days of each of the four stages depicted in the center column, with wedges sized proportionally to their duration. All pie charts are oriented with the middle of the breeding stationary period at the top center, and thus do not depict phenological differences in calendar date among species. Species with longer migrations spend less time on the breeding grounds and more time on the nonbreeding grounds, while the time spent migrating varies little across species. Illustration of thrushes reproduced with permission by Lynx Edicions. Color scheme follows [26].

We then used comparative analyses to test how interspecific variation in the geography and pacing of one annual cycle stage influences the remaining annual stages, and how the staging of the annual cycle is related to the geographic distance separating species’ breeding and nonbreeding ranges. Our approach affords a macroecological perspective on the seasonal flux of migratory animals, which is different than that provided by studies of individual movement [27,28], and specifically facilitates comparing broad-scale patterns of the pacing and geography of annual cycle stages in a consistent manner across species. Just as species’ geographic ranges are emergent properties of the home ranges occupied by individuals [29], the geographic separation of seasonal ranges emerges as a species-level property from the seasonal migration distances of individuals between breeding and nonbreeding locations. This distance, which we refer to simply as “migration distance,” reflects the seasonal movement of entire populations and highlights the extent to which breeding and nonbreeding periods are separated in geographic and environmental space [9,30,31]. Likewise, the four annual stages demarcated by our analyses (Fig. 1) indicate periods of relative seasonal geographical movement or stationarity and represent emergent species-level patterns as opposed to movements of individuals.

To illustrate the conceptual framework for our study, we highlight the annual cycle of the Wood Thrush (*Hylocichla mustelina*), whose annual cycle stages we found to be highly symmetrical: the durations of the breeding and nonbreeding stationary periods are nearly equivalent to each other, as are the durations of spring and fall migration (Fig. 1A). The geographic separation of the centroids of the breeding (summer) and nonbreeding (winter) ranges is 2569 km, which is close to the mean distance of 2600 km for all 150 species included in the analysis (Fig. S1.). Our study asks, if the durations of any annual cycle phases diverge from the seasonal symmetry exhibited by the Wood Thrush, as seen in other closely related species (Fig. 1B-E), what are the cascading effects on the remaining phases? If breeding and nonbreeding areas are closer or farther apart, what impact do these differences in migratory distance have on the pacing of migratory and stationary phases? Our comparative framework helps illuminate the relationship between the temporal and geographic dynamics of the annual cycle in migratory species, with implications for the life history tradeoffs that underly the evolution of diverse migratory strategies.

## Results

To understand how the annual cycle is structured across our study species, we first characterized the variance and phylogenetic signal of each annual cycle stage’s duration (number of days). We found that 79.3% of species had breeding stationary periods that were shorter than their nonbreeding stationary periods (Fig. 2; mean breeding stationary period duration = 90.9 days, *SD =* 26.1; mean nonbreeding stationary period duration = 131.9 days, *SD* = 27.0, *t* = - 9.70, df = 149, *P* <<0.001 paired *t*-test). Both stationary durations exhibited modest but significant phylogenetic signal (Table S2). The duration of spring migration was 3.4 days shorter on average and showed less variance than fall migration (Fig. 2; mean fall duration = 72.76 days, *SD =* 9.29; mean spring duration *=* 69.38 days, *SD* = 5.0, *t* = -4.52, df = 149, *P* <<0.001 paired *t*-test), consistent with research indicating that many bird species migrate in a narrower window in the spring than fall [32–35]. However, in contrast to the stationary periods, the durations of both migratory periods were less variable among species (Fig. 2) and consequently exhibited no phylogenetic signal (Table S2).

**Fig. 2.**
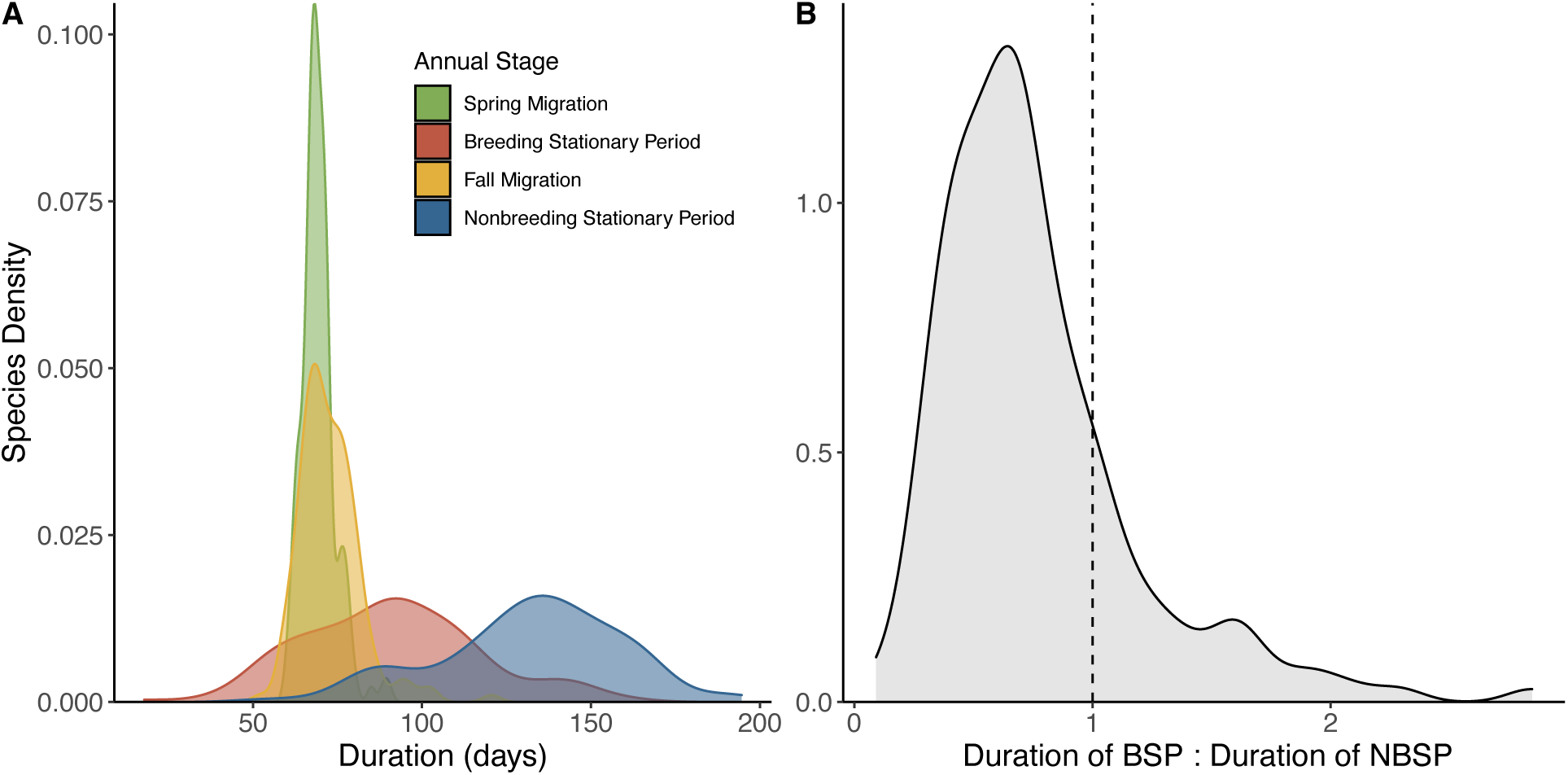
Variation in the duration of annual cycle stages. (A) Density distributions of annual cycle durations estimated for the 150 passerine bird species included in the study during four annual cycle stages. (B) Density distribution of the ratio between the duration of the breeding and nonbreeding stationary periods (BSP and NBSP, respectively) indicates that 79% of the 150 passerine bird species have shorter breeding than nonbreeding stationary periods.

To further interrogate the constraints on migratory durations, we tested the relationships between migration duration, the calendar timing of spring migration, and migration distance. Previous studies have shown that long-distance migratory species arrive on the summer breeding grounds later in spring than short-distance migratory species [30,36,37]. It is intuitive to predict that this pattern is driven by a phenological constraint imposed by migration distance, such that longer-distance migrations should require more time to migrate than shorter-distance migrations owing to the greater geographic distance covered. Therefore, we predicted that species that migrate longer distances—that is, whose stationary distributions are further apart—should have longer migratory durations. Further, we predicted that these long-distance migrations should force the well-documented phenological patterns of later spring arrival on the breeding grounds, which in our study is reflected as the date at which a species’ geographic distribution becomes stationary following northward spring movement. For each species, we quantified migration duration as the number of days that the geographic range centroid exhibits latitudinal movement during spring or fall (Fig. 1; *Materials and Methods*), spring arrival as the date demarcating the end of spring migration and the beginning of the breeding stationary period (Fig. 1), and migration distance as the geodesic distance between the mean latitude and longitude of the breeding and nonbreeding stationary periods (*Materials and Methods*). We evaluated the influence of body mass (which is thought to affect migration speed [38–40]) and breeding latitude (which is known to be positively correlated with migration distance [41]) as covariates.

Consistent with predictions, phylogenetic generalized least squares (PGLS) regression showed that the ordinal date reflecting the end of spring migration and the beginning of the breeding stationary period is positively correlated with migration distance such that long-distance migratory species transition to the stationary breeding period later in the spring (Fig. 3A, *β* = 8.044 ± 1.260, *P* << 0.001), even when including breeding latitude and body mass as covariates (migration distance *β* = 3.844 ± 1.440, *P* = 0.008; breeding latitude *β* = 6.759 ± 1.428, *P* << 0.001; log mass *β* = -3.491 ± 1.416, *P* = 0.015). However, contrary to our predictions, PGLS results show that the duration of the migratory periods does not underly the relationship between migration distance and spring phenology. Instead, we found that the migration duration of a species was uncorrelated with migration distance, whether evaluating the duration of spring and fall migration separately (Fig. 3C-D; spring *β* = 0.136±0.080, *P* = 0.108; fall *β* = 0.126 ± 0.077, *P* = 0.110) or the combined duration of both seasonal migrations (*β* = 0.105 ± 0.056, *P* = 0.065). The lack of relationship between migration duration and migration distance was robust to the inclusion of breeding latitude and body mass as covariates (migration distance *β* = 0.110 ± 0.060, *P* = 0.065), neither of which predicted migration duration (log mass *β* = 0.100 ± 0.089, *P* = 0.220, latitude *β* = 0.050 ± 0.060, *P* = 0.450).

**Figure 3.**
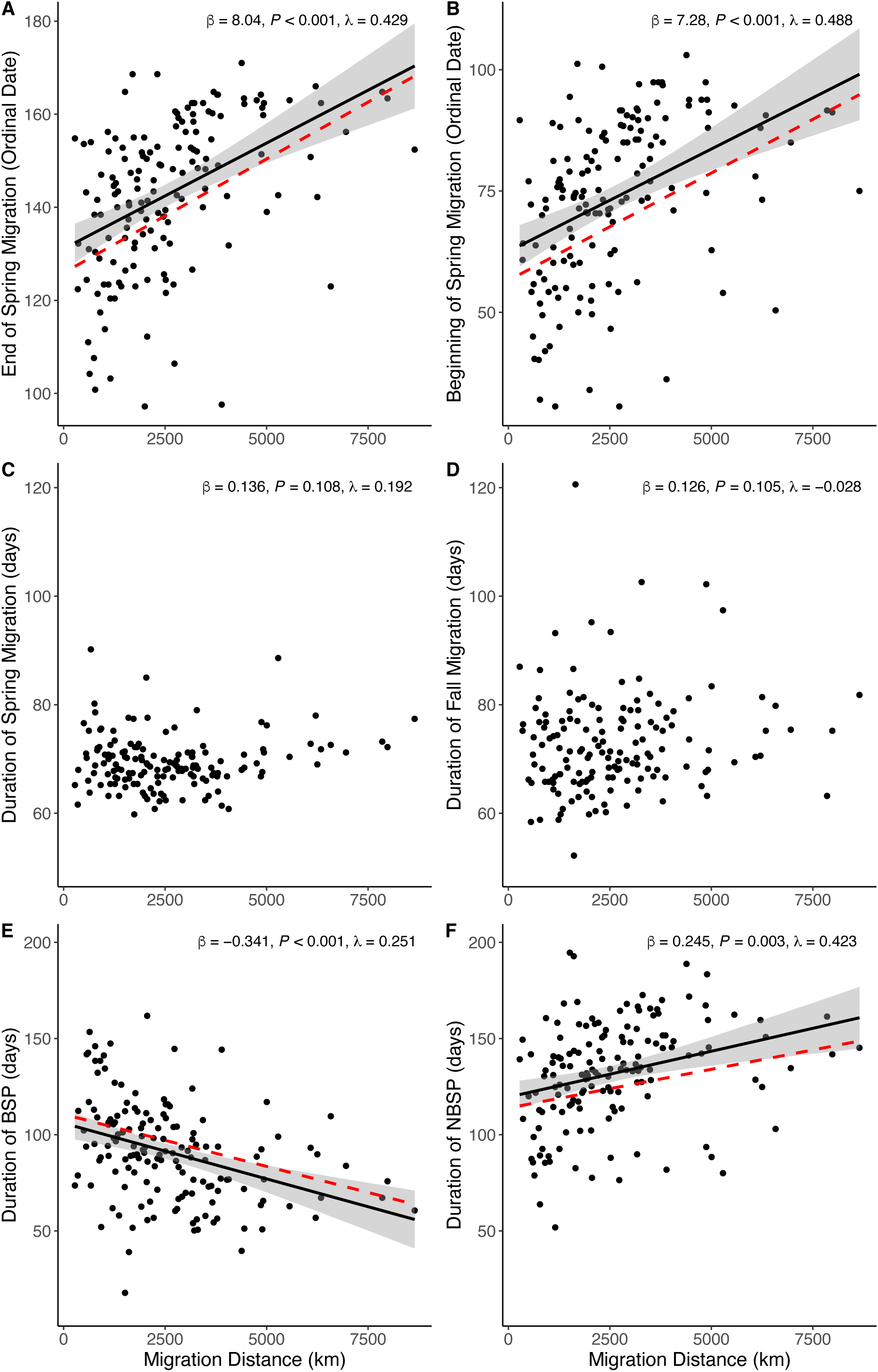
Relationships between annual cycle phenology and migration distance. There is a positive relationship between migration distance and (A) the date corresponding to the end of spring migration and the beginning of the breeding stationary period, and (B) the date corresponding to the beginning of spring migration and the end of the nonbreeding stationary period (Fig. 1, center column). Because migration distance is positively correlated with both the beginning and end of spring migration, migration distance is not correlated with the duration of (C) spring migration. Migration distance is also not correlated with (D) the duration of the fall migratory period. Migration distance is negatively correlated with the duration of (E) the breeding stationary period (BSP) and positively correlated with the duration of (F) the nonbreeding stationary period (NBSP). Slope estimates, *P*-values, and Pagel’s λ are from phylogenetic generalized least squares (PGLS) regressions of centered and standardized variables with simultaneous estimation of λ. For the four significant relationships (A, B, E and F), regression lines from PGLS are shown as the red dashed lines and ordinary least squares regression with confidence intervals as the black line and gray ribbon.

Our finding that migration duration and distance are decoupled reveals key dimensions of the phenology of seasonal range shifts in migratory species. Whereas the calendar timing of spring migration varies among species and scales with migratory distance (longer distance migrants arrive later to their breeding grounds than short-distance migrants; Fig. 3A), the duration of time in which species’ geographic distributions undergo a seasonal shift is surprisingly consistent across species (Fig. 2). Indeed, not only do long-distance migrants arrive on their breeding grounds later in spring, they also depart their non-breeding grounds (initiate spring migration) later in the year (Fig. 3B; migration distance *β* = 7.281 ± 1.287, *P* << 0.001), even after controlling for nonbreeding latitude (migration distance *β* = 10.696 ± 1.657, *P* << 0.001; nonbreeding latitude *β* = 5.903 ± 1.900, *P* = 0.001). These results indicate that the durations of the dynamic period of seasonal range movement (migration) are relatively constant across species regardless of the distances between seasonal ranges.

That migration duration is constrained and decoupled from both migration distance and migration timing carries further important implications for the evolution of the annual cycle. First, because so little variation in annual cycle pacing among species is explained by the duration of migration, most of the interspecific variation in the annual cycle is necessarily structured by the tradeoff in duration of the two stationary periods (Fig. 4). Consequently, the durations of the breeding and nonbreeding stationary periods are strikingly negatively related to one another such that species with longer breeding stationary periods necessarily have shorter nonbreeding stationary periods and vice versa (Fig. 4; *r* = -0.84, *P* <0.001, Pearson’s correlation of phylogenetic independent contrasts). Secondly, migratory distance is, intriguingly, more strongly correlated with the duration of the stationary periods (Fig. 3E-F; PGLS *β* breeding stationary period*=* -0.341±.080, *P*<<0.001; PGLS *β* nonbreeding stationary periods *=* 0.250±.080, *P*=0.003) than it is with the duration of either migratory period (see above; Fig. 3C-D), suggesting that the factors that underlie the duration of the stationary periods are paramount to explaining the evolution of migration distance and vice versa.

**Fig. 4.**
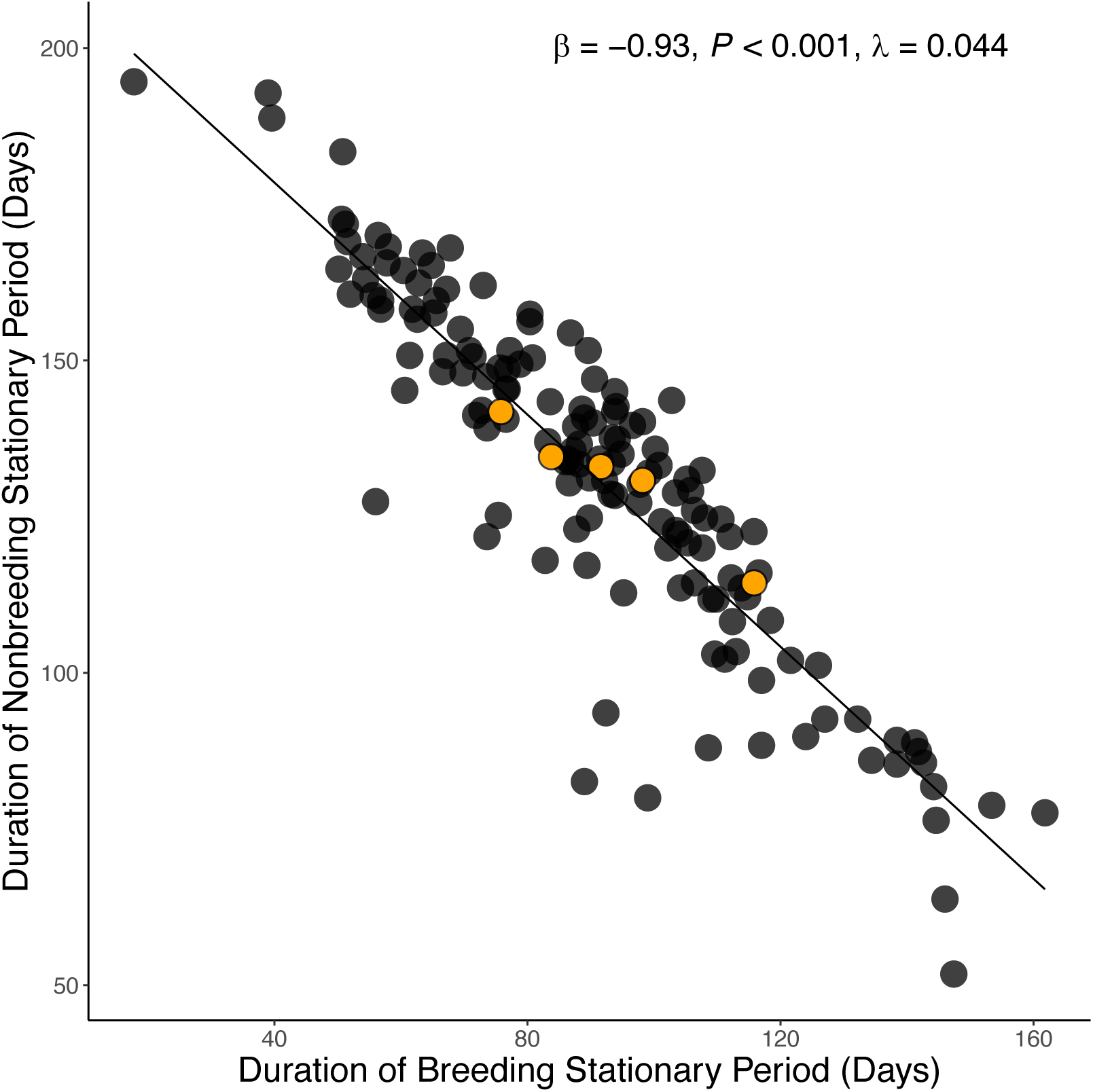
The duration of the breeding and nonbreeding stationary periods are strongly negatively correlated. The slope estimate, *P*-value, Pagel’s λ and regression line are from a phylogenetic generalized least squares (PGLS) regression of all 150 species in the study. Orange points correspond to the thrush species shown in Fig. 1.

To further understand the determinants of stationary durations and how they are related to migration distance, we used model selection to evaluate the importance of intrinsic (phylogeny, body size and pre-basic molt location) and environmental (latitude and habitat) factors on the duration of the breeding stationary period. We focused on predicting the length of the breeding stationary period as opposed to the nonbreeding stationary period because the breeding stationary period encompasses reproduction, development, and growth in our study species. We predicted breeding latitude to be negatively correlated with the breeding stationary period owing to the shorter window of food resources at high latitudes and longer daylight hours during the short growing season [52]. We predicted that body mass would be positively correlated with breeding stationary period duration due to longer developmental durations in larger species [53] and predicted moderate effects of habitat (breeding biome) owing to previous research suggesting that, for example, grassland birds spend longer on their stationary breeding grounds than species breeding in other habitats [54]. We predicted a shorter breeding stationary period in the 8.0% of included species that undergo pre-basic flight-feather molt on or near their nonbreeding grounds as opposed to their breeding grounds [22] (see Supplementary Text for discussion of molt patterns). We included migration distance and the interaction of migration distance and breeding latitude as predictors owing to evidence that migration distance and breeding durations are negatively correlated [30,55].

We found that the breeding stationary period was best explained by models that included body mass, breeding latitude, and migration distance as predictors (Table 1). Pre-basic molt location was also included in the top models but did not affect the relationship between the duration of the breeding stationary period and mass, latitude or migration distance (Table 1, Table S3, Supplementary Text). Breeding biome was not included in highly ranked candidate models (Table 1), and we found that all candidate models contained no phylogenetic signal (Pagel’s λ close to zero [56,57]). These results support a fundamental influence of body mass, breeding latitude, and migration distance on the broad scale structuring of the stationary periods regardless of phylogeny and biome. The best fitting models predicting breeding stationary period duration (Table 1) further indicated a negative relationship between breeding latitude and breeding stationary period and a positive relationship between mass and breeding stationary period (see also Fig. S2.). These results are consistent with predictions from classic life history theory wherein developmental durations scale allometrically and latitudinally [53,58], and provide confidence that our method of demarcating annual cycle durations at macroecological scales produces reliable results.

**Table 1.** Model comparison of candidate models explaining of the duration of the breeding stationary period (BSP). The second-order Akaike Information Criterion (AIC_c_) for each model and the change in AIC_c_ from the best model (ΔAIC_c_). The four top-ranked models (shown in bold; ΔAICc < 2.0) included body mass, migration distance and breeding (BSP) latitude, and the interaction of migration distance and breeding latitude as predictors. The location of pre-basic molt (see *Materials and Methods*) improved model fits slightly but did not change the relationship with other variables. Body mass was log-transformed and all variables were centered and standardized before analysis. We used ordinary least square regression models for model selection because we found that there was no phylogenetic signal in the residuals of the candidate models.

| Intercept | Mass | Latitude<br>BSP | Migration<br>Distance | Latitude BSP:<br>Migr. Dist. | Biome | Molt | df | logLik | AICc | ΔAICc | weight |
| --- | --- | --- | --- | --- | --- | --- | --- | --- | --- | --- | --- |
| **0.02** | **0.27** | **-0.20** | **-0.22** |  |  | + | **7** | **-192.47** | **399.73** | **0.00** | **0.30** |
| **0.07** | **0.27** | **-0.22** | **-0.17** | **-0.10** |  | + | **8** | **-191.56** | **400.13** | **0.40** | **0.25** |
| **0.00** | **0.24** | **-0.17** | **-0.25** |  |  |  | **5** | **-195.15** | **400.72** | **0.99** | **0.18** |
| **0.04** | **0.25** | **-0.18** | **-0.20** | **-0.09** |  |  | **6** | **-194.45** | **401.49** | **1.76** | **0.12** |
| 0.00 | 0.23 |  | -0.33 |  |  |  | 4 | -197.12 | 402.53 | 2.80 | 0.07 |
| -0.02 | 0.25 |  | -0.32 |  |  | + | 6 | -195.14 | 402.86 | 3.13 | 0.06 |
| 0.00 | 0.29 | -0.28 |  |  |  |  | 4 | -199.27 | 406.81 | 7.08 | 0.01 |
| 0.63 | 0.23 | -0.28 | -0.20 |  | + |  | 13 | -191.38 | 411.43 | 11.70 | 0.00 |
| 0.00 | 0.28 |  |  |  |  |  | 3 | -206.06 | 418.29 | 18.56 | 0.00 |
| 0.01 | 0.30 |  |  |  |  | + | 5 | -203.99 | 418.39 | 18.66 | 0.00 |
| 0.03 | 0.25 |  |  |  | + |  | 11 | -199.48 | 422.87 | 23.14 | 0.00 |
| 0.00 |  |  |  |  |  |  | 2 | -212.34 | 428.76 | 29.03 | 0.00 |
| 0.02 |  |  |  |  |  | + | 4 | -211.17 | 430.62 | 30.89 | 0.00 |

## Discussion

Our finding that migratory distance and duration are decoupled (Fig. 3B-C) is notable given that migration distances in our study species vary from *ca.* 300 to 9,000 km (Fig. S1.). This result implies that species with longer migratory journeys experience seasonal range shifts in roughly the same amount of time as species with short migrations, suggesting that long-distance migrants migrate faster. Our data do not allow us to directly link the duration in which a population is shifting latitudes to the migration tempo of individual birds, because it is difficult to disentangle a greater synchronicity of migration across individuals within a species from greater migration speed of individuals. However, our results are consistent with a positive correlation between migration speed and distance recovered in other studies [37,38,42–44]. Our results therefore provide interspecific comparative context for understanding the evolution of fast-paced long-distance migrations that have been well-documented by tracking individuals [49,50] or populations [51]. By revealing emergent species-level patterns in the pacing of migratory movements, our study lends support to the idea that migrants that have further to travel are under stronger selection to shorten the window of time during which they migrate [40,42,48], and highlights the macroecological consequences of selection on migration speed for the seasonal redistribution of populations.

Our results that migratory durations are relatively consistent across species also illuminate critical connections between the durations of the stationary periods and the timing and distance of migration. Specifically, the negative relationship we found between the length of the breeding stationary period and migration distance (Table 1, Fig. 3E) supports an emerging understanding of the significance of migration distance as a life history trait connected to the slow-fast continuum [30,55,59–61]. In a previous study of a subset of these species (boreal species breeding in the same latitude and biome [30]), species that migrate further distances away from boreal breeding grounds were shown to spend less time on the breeding grounds than shorter-distance migratory species breeding at similar latitudes. The constrained time on the breeding grounds in longer-distance migrants was correlated with truncated egg dates and lower annual fecundity [30]. Further, as predicted by life history tradeoffs in reproduction versus survival [6], long-distance migratory species also exhibited higher annual adult survival [30], highlighting migratory distance as an axis of life history variation.

Our results support the connection between migration distance and life history tradeoffs at broader geographic and taxonomic scales by showing how the stationary periods scale with migration distance (Table 1, Fig. 3E-F). Longer migrations yield shorter breeding stationary periods, even when accounting for breeding latitude, which explained less variation in breeding stationary period duration than did migration distance (Table 1). That is, as the geographic separation of the breeding and nonbreeding locations increases, the duration of time spent on the breeding grounds decreases, through effects beyond the latitudinal position of the breeding grounds (Table 1). Further, our results show that this phenomenon is not driven by the greater time required to achieve long-distance migration because migratory durations are not lengthened by increasing migration distance (Fig. 3C-D). Therefore, our results suggest that a shortened breeding stationary period and lengthened nonbreeding stationary period are epiphenomenal outcomes of greater geographic separation between breeding and nonbreeding grounds.

Our example of the Wood Thrush and its close relatives (Fig. 1) illustrates the novel dynamics we have identified here. The Wood Thrush has breeding and nonbreeding stationary periods both of ∼115 days, and spring and fall migration both of ∼67 days (Fig. 1). Hypothetically, if a different thrush species’ breeding and nonbreeding ranges were further apart geographically (Fig. 1), that species could have a breeding and nonbreeding stationary period of the same duration as the Wood Thrush, so long as its longer-distance migration did not require more days. Alternatively, if longer-distance migration requires more time, the lengthened migratory periods could shorten both the breeding and nonbreeding stationary periods equally and maintain balanced durations. However, neither possibility is supported by our results. Instead, we found 1) that migration duration does not vary across species depending on distance (Fig. 1, Fig. 3C-D), and 2) that the breeding and nonbreeding stationary periods scale in opposite directions with migration distance (Fig. 3E-F) and inversely with one another (Fig. 1, Fig. 4), such that geography and timing are interrelated.

Because migration is a geographic phenomenon connecting different seasonal distributions, our results imply that the biogeographic distributions of migratory species are both drivers and outcomes of the temporal pacing of the annual cycle. Specifically, our results suggest that any biological factor that increases the duration of the breeding stationary period—e.g., by lengthening the time devoted to the various stages of reproduction and development [53] or post-breeding molt [62,63]—not only inherently decreases the duration of the nonbreeding stationary period (Fig. 4) but is predictive of reduced geographic separation of the breeding and nonbreeding ranges and therefore shortened migration distance (Fig. 3). Conversely, a species with a longer migration distance inherently has a shorter breeding period, constraining the amount of time available for reproduction and development but extending the amount of time spent on the nonbreeding grounds. Owing to the strong negative relationship between breeding and nonbreeding durations (Fig. 4), it follows that selection on the duration of the nonbreeding period should have inverse effects.

The mechanisms underlying these patterns demand further study, but we suggest that the phenological patterns shown here are driven by how selection on the balance between investment in annual reproduction and annual survival (life history strategy) relates to the timing and geography of the annual cycle. That is, species that have truncated breeding periods likely produce fewer offspring each year, but balance this reduced fecundity with greater time spent in resource-rich nonbreeding environments that increases annual survival—a strategy that demands longer-distance migration between high latitude breeding and low-latitude nonbreeding locations [30]. In other words, endless combinations of breeding and nonbreeding distributions, and migratory journeys between them, are possible for seasonal migrants, as evidenced by the diversity of migratory patterns observed in the world’s birds. But to be viable, each combination of breeding and nonbreeding ranges demands a specific annual schedule that balances annual reproduction versus survival, leading to integrated evolution of migratory geography and phenology. Selection tunes the balance for each migratory population in subtle and variable ways, but at a broad scale, has led to the emergent interspecific patterns we reveal here of shorter summer and longer winter stationary periods for long-distance migrants, and longer summers and shorter winters for short-distance migrants.

A potential limitation of our study is that by focusing on the species as the unit of comparison, we cannot assess variation in annual cycle timing among populations within a species or individuals in a population to test mechanisms directly. That is, geographically widespread species exhibit spatial variation in the timing of their annual cycles that is reduced here to species-level averages. However, by focusing on emergent species-level patterns, our study reveals how the structure of species’ annual cycles differ broadly in a comparative framework, thereby illuminating patterns whose mechanisms can be interrogated at the individual and population level in future research using different approaches [65]. We predict that many of the relationships we have demonstrated between migration distance, breeding latitude and the durations of the annual cycle phases may be recapitulated among populations within a species and therefore relate to intraspecific life history variation.

The journeys of long-distance migratory birds are highly celebrated and studied, but it has been difficult to understand why species differ so greatly in their migratory strategy. In any given temperate biome exist species that migrate great distances to far away nonbreeding grounds during the winter and closely related species that migrate much shorter distances to closer nonbreeding grounds [8,66,67]. By elucidating the connections between the geography of migration and the temporal structuring of the annual cycle, our study helps untangle this puzzling complexity. We have shown that migration distance is a fundamental predictor of the duration of time during which breeding and development occurs, alongside the well-known effects of latitude and mass. Breeding durations are in turn predictive of both the location and the amount of time spent in stationary nonbreeding locations. Therefore, migration distance predicts not only where species are found throughout the year but also how much time they spend in each location and thus how much time can be devoted to reproductive processes versus survival to the next breeding opportunity. The evolution of biogeographic distribution, migration phenology, and life history strategy in migratory species are tightly linked.

## Materials and Methods

### Study system

We focused our study on migratory passerine bird species breeding in North America, owing to the wealth of comparable observational data on their annual cycle distributions. We started with 228 species of passerines previously identified [31] to have breeding distributions that extend north of 23° latitude. This latitude coincides with a biogeographic transition zone between an avifauna breeding in the temperate zone that primarily undergoes regular, long-distance migrations along a north-south (latitudinal) axis and therefore appropriate for the analytical approaches employed here, and an avifauna breeding in the subtropics and tropics that is mostly sedentary or that undergoes short-distance migrations primarily along an elevational axis [31,68]. We removed species with migration distances of less than 300 km (*n* =22 species [31]) because we found the latitudinal difference between breeding and nonbreeding ranges was not great enough to quantify seasonal variation in latitudinal position using our methods, leaving 206 species. After further analyses to determine the robustness of data and the reliability of our methods for demarcating the annual cycle for each species, we filtered this initial set of species to include 150 species based on criteria discussed below (see *Species Filtering*, below).

### Calculating daily geographic centroids

To calculate the seasonal change in latitude for each species, we first calculated daily geographic centers of occurrence for each species using the following approach. We compiled eBird [21] checklists from across the Western Hemisphere (170° to 30° W longitude and 60° S to 90° N latitude) during the period 1 January 2002 to 25 September 2021. Observations entered into eBird are organized in checklist format containing species observed and the time and location (longitude and latitude) of the sampling event. We first compiled checklists by day across years within equal-area hexagonal cells (49,811 km^2^). The hexagon cells were contained in an icosahedral discrete global grid system defined by a Fuller icosahedral projection using an aperture four hexagon partition method [69,70]. For each hexagon cell and day, we counted the total number of checklists and the number of checklists where each species was observed. We calculated each species’ daily geographic center of occurrence using the *n*-vector framework for geographical position calculations [71]. This framework uses the normal vector to the Earth ellipsoid (the *n*-vector) as a nonsingular positive representation. We first extracted the latitude and longitude of the center of the hexagon cells. We converted these two-dimensional geodetic coordinates into three-dimensional *n*-vectors and calculated the weighted average *n*-vector for each day using the species’ frequency of occurrence as a weighting factor. We then converted the *n*-vector back to latitude and longitude. We implemented the *n*-vector calculations using the nvctr R package [72].

### Demarcating the annual cycle

We characterized how the latitude of each species’ daily geographic centers of occurrence changed across the year using the following approach. Building on previous methods [73], for each species we fit 37 Generalized Additive Models (GAM; [74]) to latitude as a function of day using 37 different smooth term basis dimensions (*k* = 4-40); we found this range of *k* to reliably capture the meaningful range of variation in GAM fits for the data. We used a cyclic penalized cubic regression spline in each GAM model to smoothly join the first day (1 January) and last day (31 December) of the year. To estimate the dates representing the beginning and end of spring and fall migration for each species, we calculated the first and second derivatives of the GAM fits for each of the 37 basis dimensions. The second derivatives represent changepoints in the acceleration of latitudinal change over time, thereby capturing the dates when geographic range movement begins and ends in both the spring and fall. This approach allowed us to define periods of range movement (spring and fall migration) as well as the stationary periods occurring between migrations (Figs. 1, S3).

To identify second derivatives that reliably correspond to the boundaries of spring and fall migration (as opposed to more subtle and idiosyncratic variation occurring during stationary periods), we first identified the most positive and negative first derivatives of the GAM functions. These first derivatives represent the greatest change in latitude over time for northbound (spring) and southbound (fall) migration and can be interpreted as the date of “peak” spring or fall migration. We then identified the second derivatives, representing the beginning and end of the period of migratory latitudinal change, within a 40-day window before and after the first derivatives in spring and fall. To minimize the influence of outliers, we repeated the calculation of first and second derivatives for all 37 GAM functions and used the 0.1 and 0.9 quantiles of the distribution of days to estimate the beginning and end, respectively, of both spring and fall migration. In exploratory visualizations, we found this method reliably bracketed changepoints in the latitudinal curves. We used the mgcv R package [74] to implement the GAM analysis and the features R package [75] to estimate first and second derivatives from the GAM fits.

For the species included in our analysis (see next section on *Species Filtering*), our method reliably demarcates a higher latitude period of relative stationarity occurring between the cessation of spring migration and the initiation of fall migration in the Northern Hemisphere summer months (the breeding stationary period), a lower latitude stationary period occurring between the cessation of spring and initiation of fall migration in the Northern Hemisphere winter months (the nonbreeding stationary period), and two periods of dynamic population movement between the two stationary periods (Figs 1, S3). The latter two periods correspond to northward (spring or pre-breeding) and southward (fall or post-breeding) migration. This method captures the average position throughout the annual cycle of a species’ entire population (Fig. 1).

### Species Filtering

For the 206 species initially included, we plotted the daily latitudinal centers of occurrence, GAM fits, and the changepoints (annual phase boundaries) identified by the second derivative analysis to visually assess the success and reliability of this method for demarcating the annual cycle in each species. We found that for some species, changes in daily latitudinal positions were too subtle, erratic, or irregular to reliably demarcate the annual cycle phases using our methods. In these species, changepoints were identified in times of year that clearly did not match known timing of migration or breeding, or GAM fits showed uninterpretable irregularity.

To identify a set of reliably usable species, we filtered species using the following quantitative criteria. 1) We removed species with the largest geographical ranges (greater than the 0.95 quantile in breeding range area within North America [31] across all initially included species, *n* = 11 species). The large amount of geographical variation in migration phenology in these large-ranged species (*e.g., Hirundo rustica* and *Agelaius phoeniceus*) led to plots that were difficult to interpret. 2) We removed species with the smallest change in latitude between breeding and nonbreeding stationary periods (less than the 0.05 quantile of latitudinal change across all initially included species, *n* = 11 species), as we found that latitudinal change over time was not reliably captured in those species with the greatest geographic overlap in breeding and nonbreeding ranges. 3) We removed 35 species with the lowest (less than 0.10 quantile across species) expected degrees of freedom in GAM functions or lowest GAM signal to noise ratio (less than 0.10 quantile, calculated in the features R package [75]), as these species showed latitudinal change over time that was too subtle or the latitudinal data was too messy to reliably fit a GAM of latitude over time.

The union of these filtering criteria resulted in 47 species excluded from analysis, as some species failed to pass multiple filtering criteria. Lastly, we removed an additional 9 species that passed these criteria but for which visual assessment of latitude-date plots revealed obvious inaccuracies with estimation of latitude over time. Collectively, this filtering removed 56 species, leaving a total of 150 species for analysis (Table S1). The 56 species excluded from the analysis tended to be species that undergo irregular irruptive movements rather than regular latitudinal seasonal migrations, those whose primary axis of migration is longitudinal, those whose geographic distributions are very broad and include populations exhibiting a wide diversity of migratory strategies, or those for which data was less reliably available throughout the full annual cycle. Additionally, excluded species tended to be species whose breeding and nonbreeding geographic ranges are less separated such that seasonal variation in latitude was more subtle. By contrast, the species for which our method worked well to demarcate the annual cycle phases are those that undergo regular, annual migrations primarily along a latitudinal axis. Thus, by necessity, our filtering inherently focuses the analysis on somewhat longer-distance migrants with regular latitudinal movement. However, we do not expect this would bias our results, and filtering to species for which the method was most useful improved the quality and reliability of the annual cycle demarcations.

### Calculating predictor variables

Our models included breeding and nonbreeding latitude, migration distance, mass, breeding biome, and pre-basic molt location as predictors or outcome variables. We calculated breeding and nonbreeding latitudes as the average latitude of centroids during the breeding and nonbreeding stationary periods, respectively, and migration distance as the geodesic distance between the average latitude and longitude during the breeding and nonbreeding stationary periods. To avoid issues with averaging geographic positions across latitudes, we first converted latitude and longitude to an equal-area (Mollweide) projection before averaging the geographic positions of centroids. We used species’ body mass estimates from reference [31], which were compiled from reference [76]. To classify species’ breeding habitat, we used the breeding biome designations from reference [77]. We followed reference [22] in categorizing species’ principal pre-basic molt locations (the location of energy intensive replacement of flight feathers following breeding) as occurring on the breeding grounds, nonbreeding grounds, or during fall migration (molt-migration). We categorized molt patterns of three species not included in reference [22] based on consulting other literature [20,24]. We provide a fuller discussion of the impact of molt-migration on the breeding stationary period in the Supplementary Text.

### Modeling

We assessed the relationships between the durations estimated for the two migratory periods and two stationary periods and their predictors primarily using phylogenetically corrected correlations and phylogenetically corrected linear models. We centered and standardized all predictor variables and log-transformed body mass to improve its distributional properties. We checked models to ensure they met the assumptions of linear modeling. In some models, a semi-log relationship between migration distance and the predictor variables was evident, and log transformation of migration distance improved the normality of model residuals. However, this transformation did not influence our results, and so we elected not to transform migration distance in the models presented here.

To test for phylogenetic signal in the annual cycle stages (as individual trait vectors), we used the picante R package [78]. To assess relationships between variables, we performed phylogenetic generalized least squares (PGLS) regression using simultaneous estimation of Pagels’ λ [79] using the corPagel function in the ape R package [80] and the gls function in the nlme R package [81]. We found that λ values were typically very low in our models, and therefore ordinary least squares linear models produced similar results to those presented here. For phylogenetically corrected models, we used a phylogeny generated from references [31,82] in which phylogenetic information was downloaded from birdtree.org [83] (‘Hackett all species’ dataset) as a sample of 2000 phylogenetic trees and a consensus tree calculated from these data using the SumTrees program in the ‘DendroPy’ python package [84].

For comparing candidate models explaining the duration of the breeding stationary period (Table 1), we used a model selection approach. For this test, we present ordinary least square regression because we could not reliably fit all candidate PGLS models using simultaneous estimation of Pagel’s λ, and the λ value of the residuals of each candidate model (evaluated with the phylosig function in the phytools R package [85]) were close to zero, indicating no phylogenetic effects in the model. To assess relative model performance in predicting the breeding stationary period, we calculated the second-order Akaike Information Criterion (AICc) in the MuMIn R package [86].

## Acknowledgements

We thank Andrea Benavides Castaño, Jacob Berv, Eric Gulson-Castillo, Natalie Hofmeister, Abby Kimmitt and Kristen Wacker for helpful feedback on earlier versions of the manuscript. We thank the eBird participants, expert reviewers and the many other individuals whose work and engagement make the eBird project successful.

## Supporting Information

### Supplementary Text. The influence of molt migration and migratory double breeding on the breeding stationary period

For all the passerine bird species considered in the study, the breeding stationary period (BSP) is when reproduction (from nest building through growth of fledglings) occurs. The majority (71%) of the 150 species included in our study also undergo an energy intensive full feather molt (the pre-basic molt) on or near the breeding grounds prior to the initiation of fall migration and thus during the period demarcated as the breeding stationary period [1,2], whereas only 12 species (8.0%) are thought to molt during the nonbreeding stationary period [1–3] (Table S1). However, some passerine species are known to interrupt migration to undergo a flight feather molt (often referred to as “molt migration” [4], or the more specific “stopover pre-basic molt-migration”[1]). The interruption of migration to molt at a location distinct from either the breeding or nonbreeding stationary period could complicate demarcation of the annual cycle phases using our method. Therefore, we categorized species as molt-migrants based on reference [2] and assessed how the possibility of molt migration intersects with our methods of demarcating the annual cycle. Additionally, we considered that a small number of species with western distributions are known to breed for a second time after a southward movement into the Mexican monsoon region [5].

We found that a similar proportion of species included versus excluded from the study based on our filtering criteria (see Methods for details) had documented molt migration (22.6% of included versus 16% of excluded species were molt-migrants), and thus the possibility of molt migration did not bias the reliability of our methods for demarcating the major stationary and migratory periods. Among the included species, we found that some species of molt migrants exhibited slower-paced and more subtle southward shifts in latitude towards the end of the breeding stationary period, but prior to the dynamic period of fall migration (e.g., *Passerina ciris*, Fig. S3 panel 150, a species that is also a double breeder [5]). However, such patterns were also exhibited in some species without documented molt migration or double breeding (e.g., *Setophaga citrina*, Fig. S4 panel 117 or *Mniotilta varia*, Fig. S3 panel 105), and other known double-breeders did not exhibit this latitudinal pattern of multiple stationary stages (e.g., *Icterus spurius,* Fig S3 panel 91). Thus, it is unclear whether these more complex latitudinal patterns represent actual population-level movements across a species’ annual cycle stages such as molt or double-breeding. Regardless, these more subtle movements were reliably encompassed by our methods as falling within the breeding stationary period, and therefore this stage should be interpreted as a period with *relatively* greater stationarity than the obvious period of spring or fall migration.

Finally, we found using a phylogenetic ANOVA that the duration of the breeding stationary period did not differ significantly between molt migrants and species that undergo prebasic molt on their breeding grounds, indicating that molt migrations involve a relatively subtle latitudinal movement that did not influence our categorization of the breeding stationary period (Table S3). The breeding stationary period of the 8% of species that undergo pre-basic flight feather molt on their nonbreeding grounds also did not differ significantly from the breeding stationary period of other species, though the trend was towards shorter breeding stationary periods, as expected given the time required for molt (Table S3).

---

**Table S1.** (separate .xlsx file). Species included in the study and their covariates used in modeling. Tab 1: The 150 species that passed filtering criteria (*Materials and Methods*) and the attributes that we calculated and included as covariates or predictors in our models as described in *Materials and Methods*. Tab 2: The 55 species that were initially included in calculation of annual cycle durations but did not pass our filtering criteria and were excluded from modeling and hypothesis testing as described in *Materials and Methods: Species Filtering*.

**Table S2:** Phylogenetic signal for each of the four annual stages.

|  | **K** | **PIC OBS** | **PIC RANDOM** | **P** | **Z** |
| --- | --- | --- | --- | --- | --- |
| **BSP** | 0.138 | 110.908 | 167.664 | 0.007 | -1.753 |
| **NBSP** | 0.128 | 131.186 | 178.740 | 0.047 | -1.472 |
| **SM** | 0.098 | 5.085 | 5.898 | 0.327 | -0.549 |
| **FM** | 0.072 | 24.084 | 21.309 | 0.760 | 0.469 |

BSP = breeding stationary period, NBSP = nonbreeding stationary period, SM = spring migration, FM = fall migration. K is Blomberg's K statistic, PIC OBS is the variance of phylogenetic independence contrasts (PIC), PIC RANDOM is the mean PIC variance of 999 tests with data randomly shuffled with respect to phylogeny, and *P* and *Z* refer to the observed value versus the randomization test. The breeding and nonbreeding stationary periods had significant and marginally significant K values respectively, but phylogenetic signal was low for all annual stages.

**Table S3:** Comparison of breeding stationary period duration by pre-basic molt location.

| Molt Location | Breeding grounds | Molt-migrant | Nonbreeding grounds |
| --- | --- | --- | --- |
| Breeding grounds | 0.000 | -0.325 | 1.413 |
| Molt-migrant | 0.325 | 0.000 | 1.474 |
| Nonbreeding grounds | -1.413 | -1.474 | 0.000 |

Pairwise *t*-values from a phylogenetic ANOVA performed in phytools<sup>59</sup> with pairwise posthoc tests between all categories. Species that molt on their stationary nonbreeding grounds had shorter breeding stationary periods than those that molt on their breeding grounds or during fall migration. However, none of the pairwise tests were significant ( $P=1.0$ for all pairwise comparisons).

**Fig. S1.**
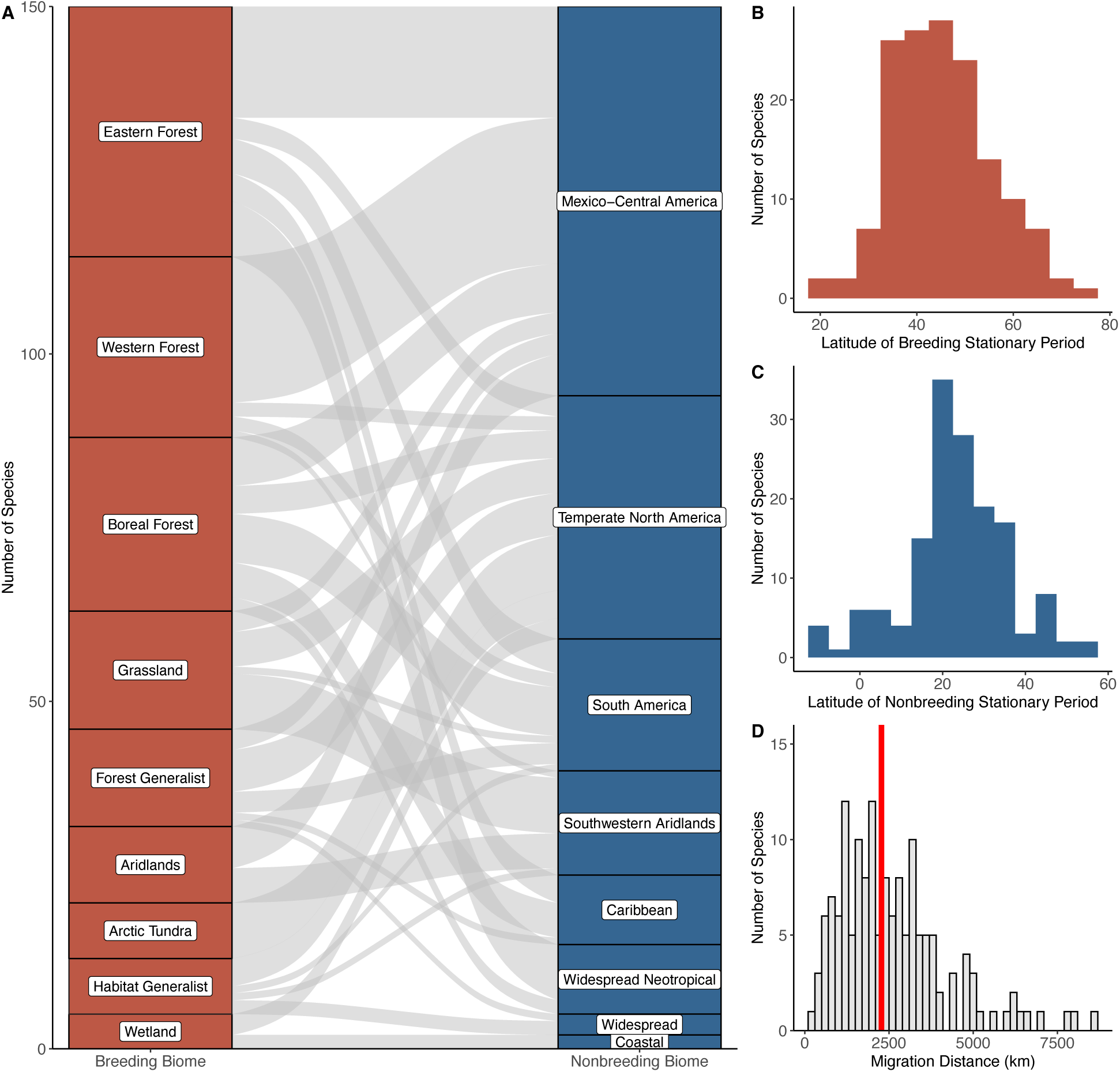
Migratory connections between stationary breeding and nonbreeding biomes and latitudes in the study system. (A) Breeding and nonbreeding biomes and the migratory connections between them for the 150 passerine bird species included in the study. The width of the curves is proportional to the relative number of species that breed and winter in the biomes connected by the curves. Number of species by (B) breeding latitude of range centroid, (C) nonbreeding latitude of range centroid and (D) migratory distance (representing geographic distance between breeding and nonbreeding range centroids). The red bar indicates the median migration distance among the 150 species in the study.

**Fig. S2.**
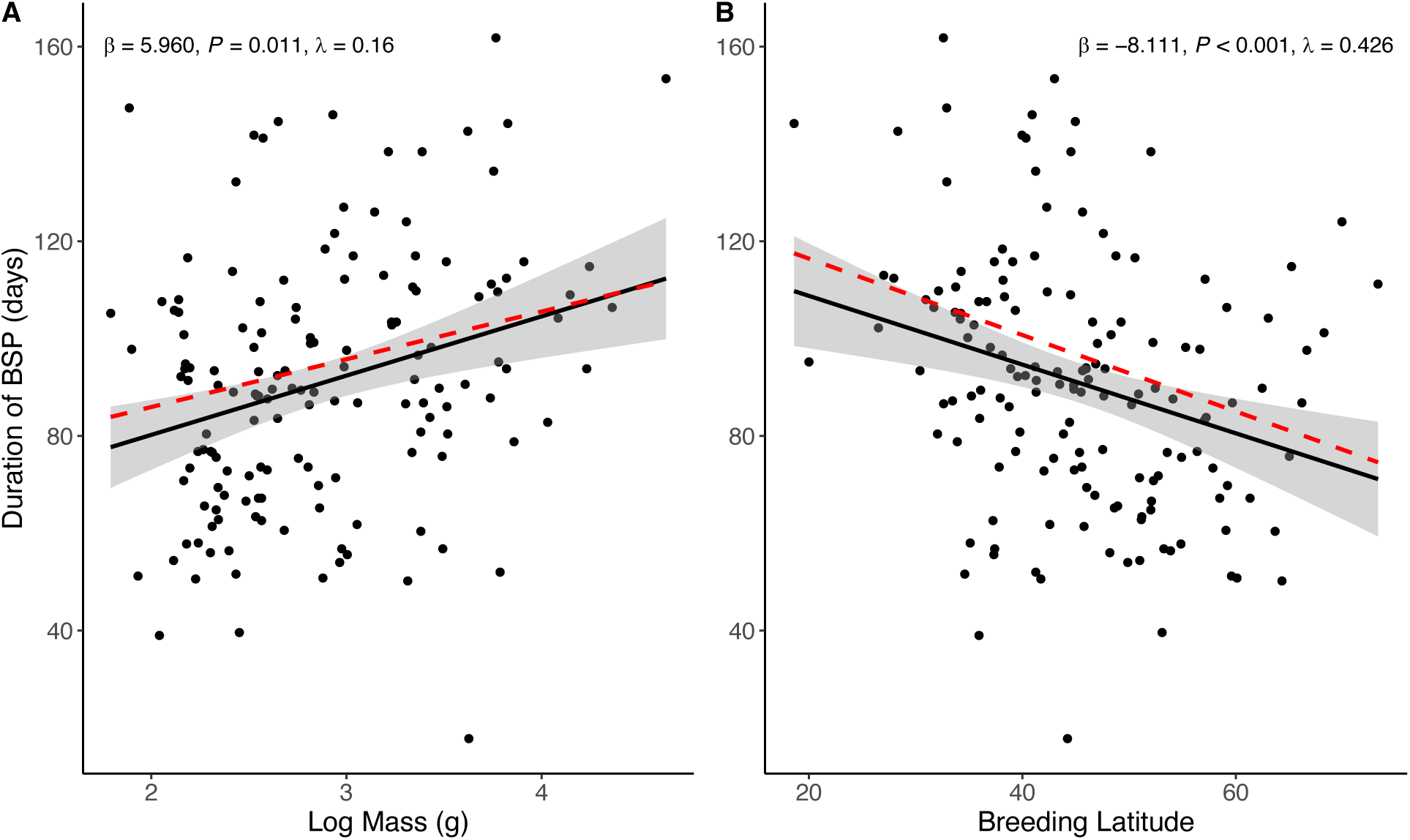
The duration of the breeding stationary period is predicted by body mass and latitude. The relationship between (A) body mass and (B) breeding latitude (latitude of the breeding stationary period) and the duration of the breeding stationary period (BSP). The slope estimates, *P*-values, Pagel’s λ and red-dashed lines are from phylogenetic generalized least squares (PGLS) regressions of centered and standardized variables with simultaneous estimation of λ. The black lines with confidence intervals depict ordinary least squares regression.

**Fig. S3.** (separate PDF file). Annual cycle demarcations for each of 150 species included in the study. Plots are ordered following the checklist of the American Ornithological Society North American Classification Committee and numbered according to Table S1. Latitudes from daily geographic centroids are shown as black dots (one point per day), derived from 20 years of eBird data. Dashed vertical lines indicate the maximum (spring, green) and minimum (fall, gold) first derivatives of GAM fits, signifying the fastest change in latitude over time and corresponding to peak northbound (spring) and southbound (fall) migration. Solid vertical lines indicate the maximum and minimum second derivatives of GAM fits and demarcate the beginning and end of the spring and fall migratory periods. These first and second derivative (dashed and solid vertical lines) were drawn from the distribution of derivatives from 37 individual GAM fits (see *Methods*). The nonbreeding (blue) and breeding (red) stationary periods signify periods of relative latitudinal stasis for the species.

## Notes

### Competing Interest Statement

The authors have declared no competing interest.

