## Supplementary material for "Evolutionary integration of the geography and pacing of the annual cycle in migratory birds": Fig. S3

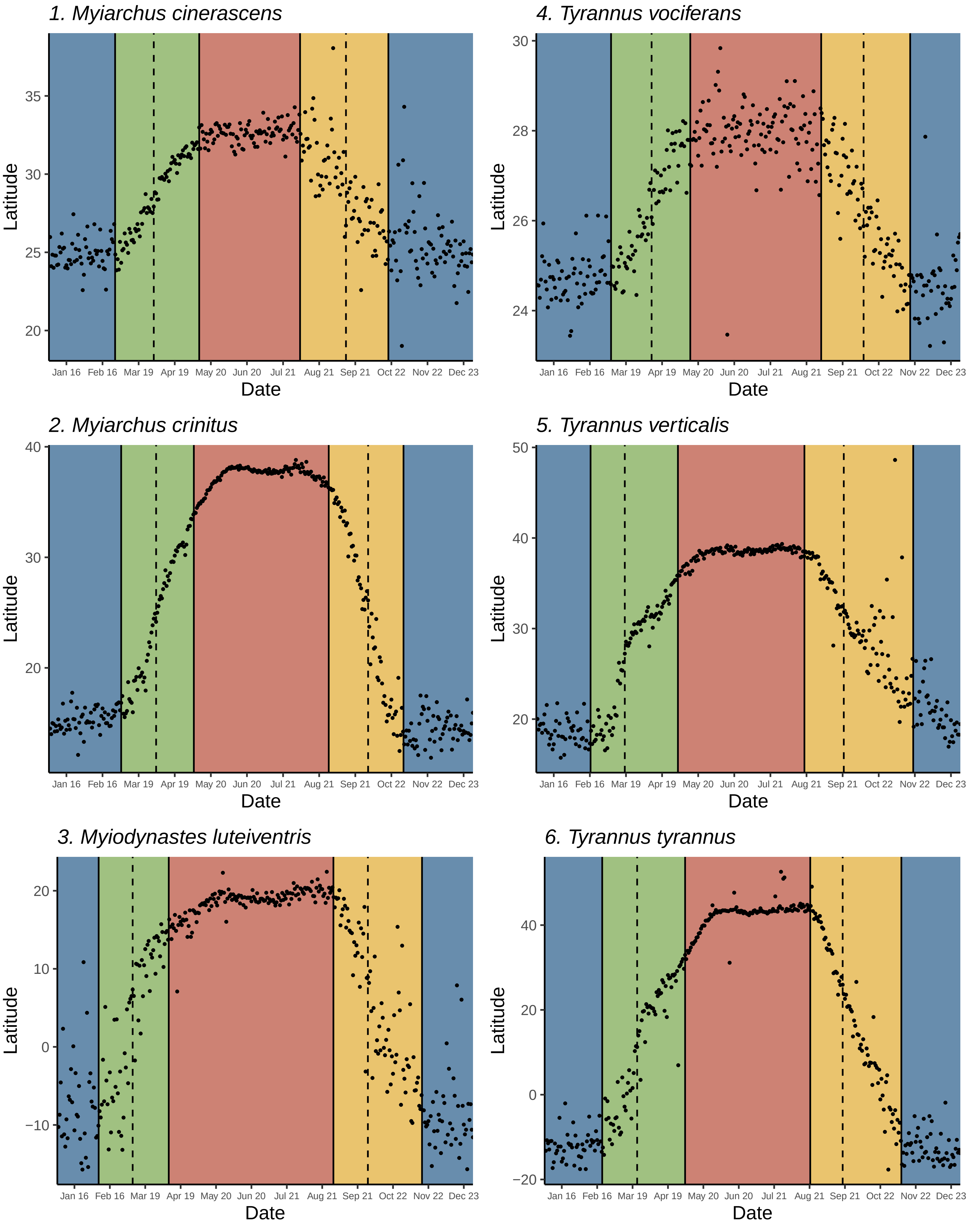
1. Myiarchus cinerascens
4. Tyrannus vociferans
2. Myiarchus crinitus
5. Tyrannus verticalis
3. Myiodynastes luteiventris
6. Tyrannus tyrannus
Latitude
Date
Jan 16 Feb 16 Mar 19 Apr 19 May 20 Jun 20 Jul 21 Aug 21 Sep 21 Oct 22 Nov 22 Dec 23

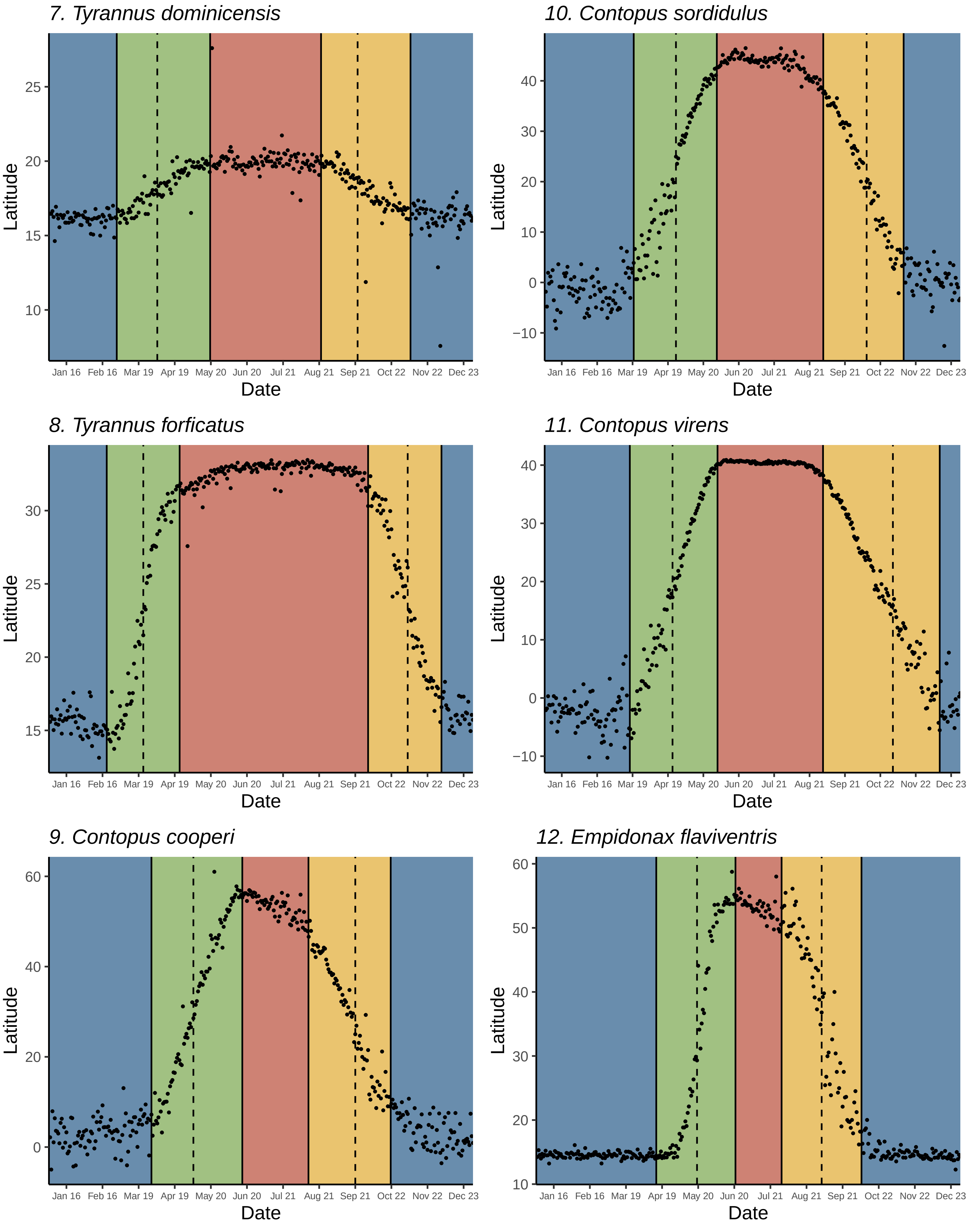
7. Tyrannus dominicensis
10. Contopus sordidulus
8. Tyrannus forficatus
11. Contopus virens
9. Contopus cooperi
12. Empidonax flaviventris
Latitude
Date
Jan 16 Feb 16 Mar 19 Apr 19 May 20 Jun 20 Jul 21 Aug 21 Sep 21 Oct 22 Nov 22 Dec 23

*13. Empidonax virescens*

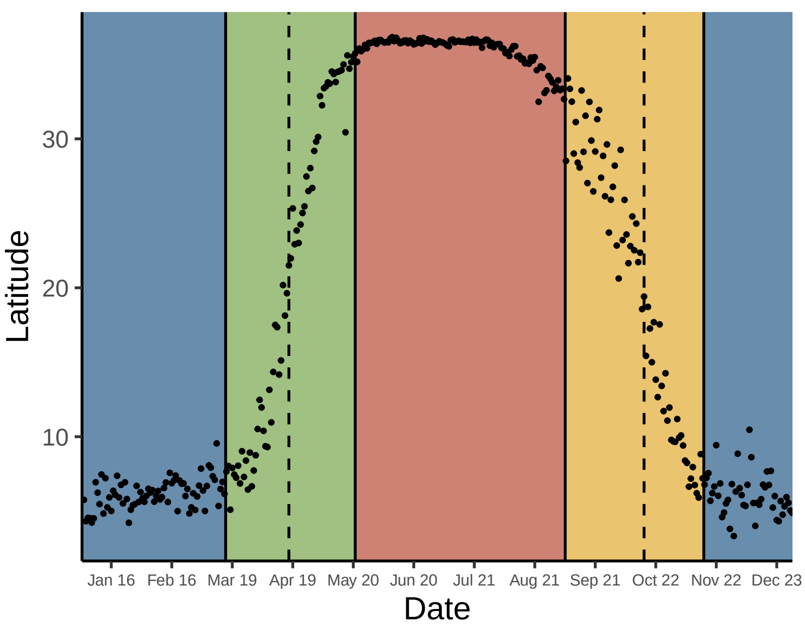


*16. Empidonax minimus*

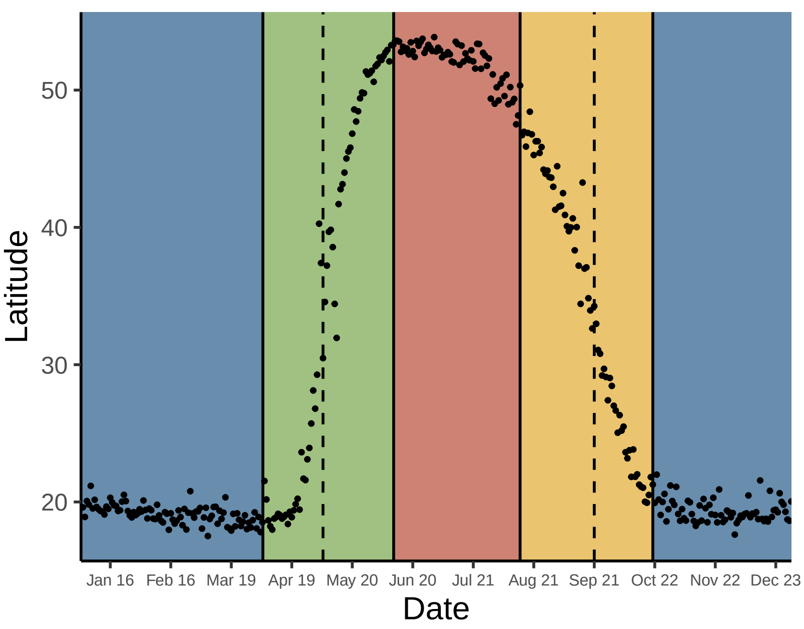


*14. Empidonax alnorum*

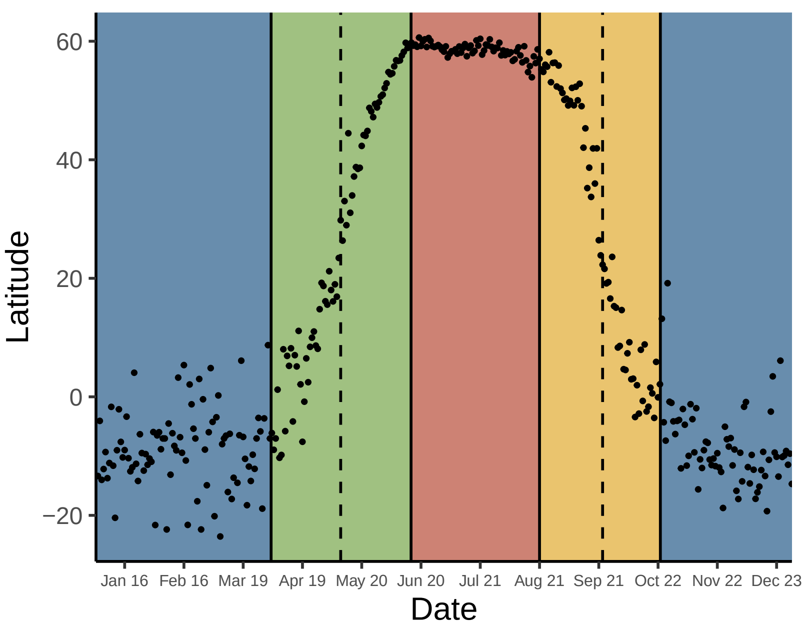


*17. Empidonax hammondii*

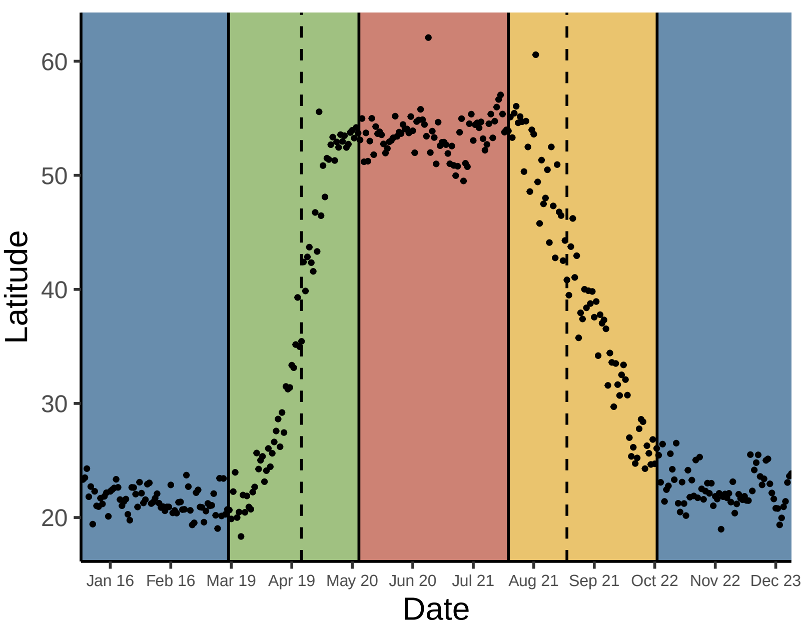


*15. Empidonax traillii*

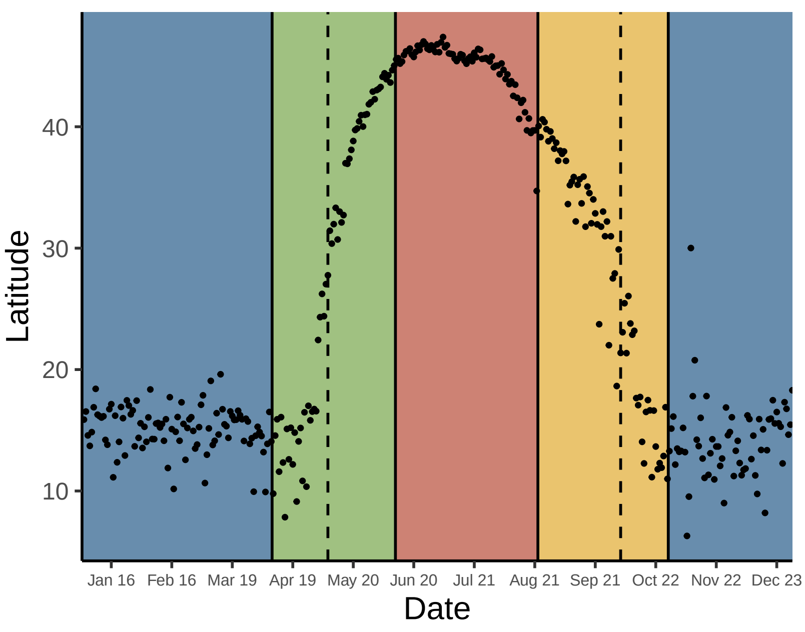


*18. Empidonax wrightii*

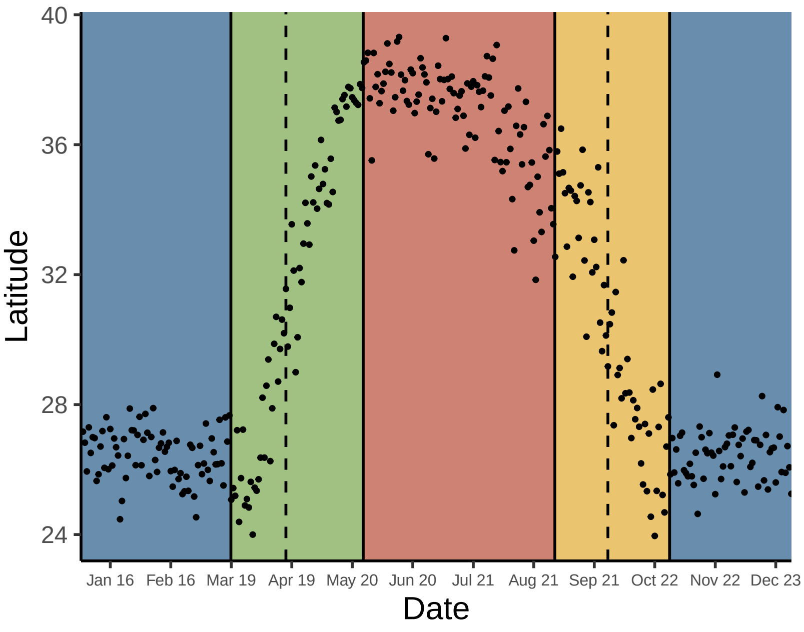

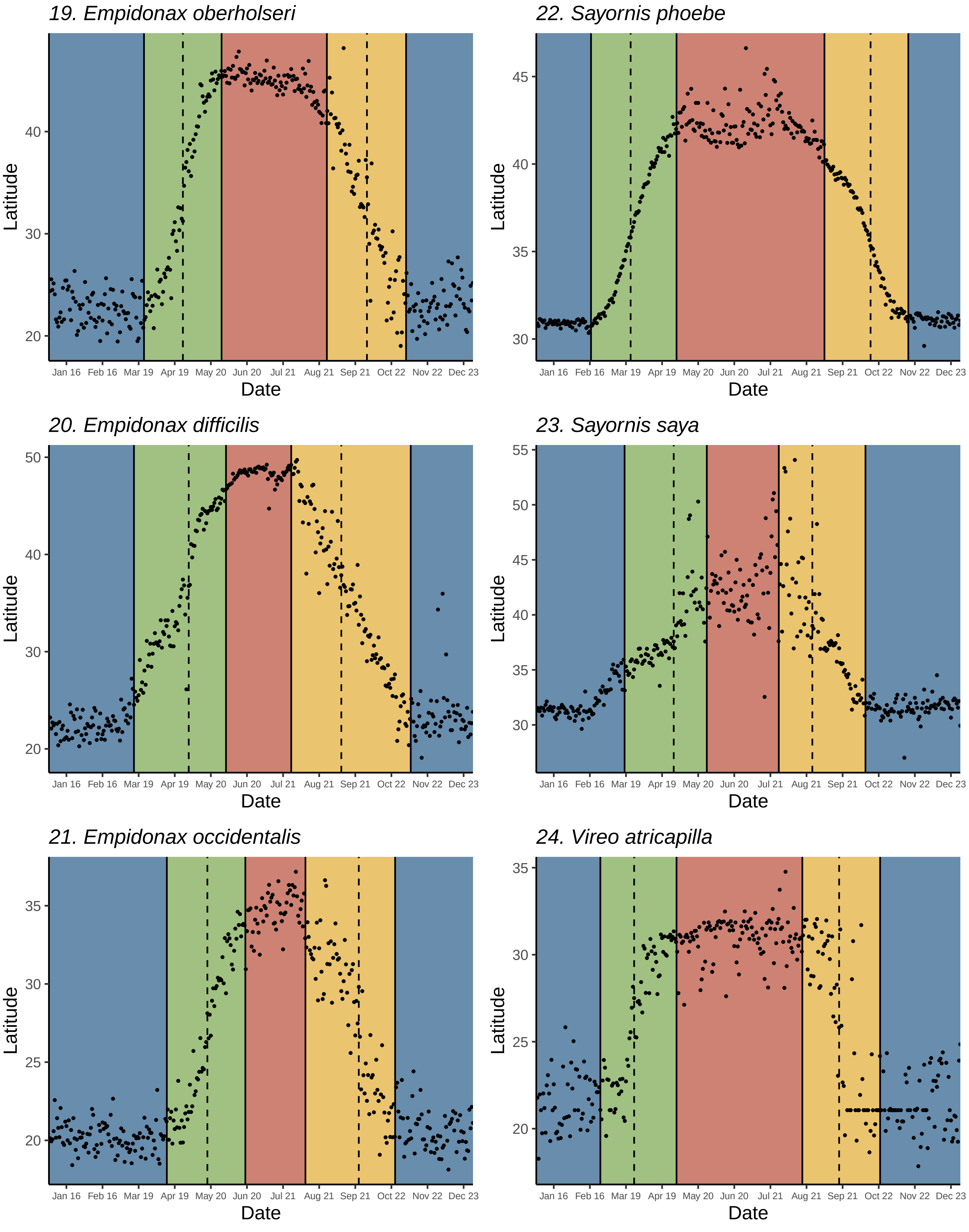
19. Empidonax oberholseri
22. Sayornis phoebe
20. Empidonax difficilis
23. Sayornis saya
21. Empidonax occidentalis
24. Vireo atricapilla
Latitude
Date
Jan 16
Feb 16
Mar 19
Apr 19
May 20
Jun 20
Jul 21
Aug 21
Sep 21
Oct 22
Nov 22
Dec 23

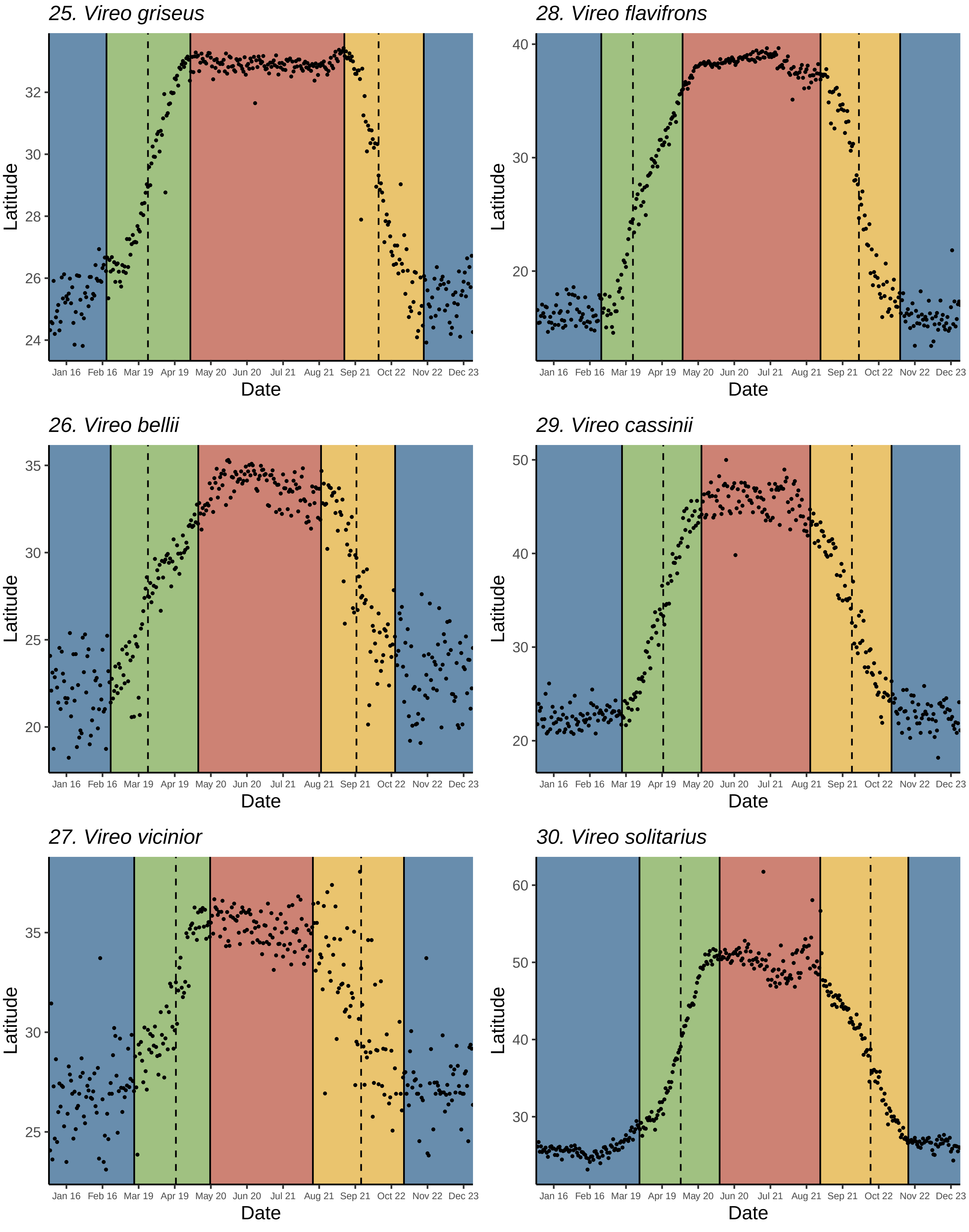
25. Vireo griseus
Latitude
24
26
28
30
32
Jan 16 Feb 16 Mar 19 Apr 19 May 20 Jun 20 Jul 21 Aug 21 Sep 21 Oct 22 Nov 22 Dec 23
Date
28. Vireo flavifrons
Latitude
20
30
40
Jan 16 Feb 16 Mar 19 Apr 19 May 20 Jun 20 Jul 21 Aug 21 Sep 21 Oct 22 Nov 22 Dec 23
Date
26. Vireo bellii
Latitude
20
25
30
35
Jan 16 Feb 16 Mar 19 Apr 19 May 20 Jun 20 Jul 21 Aug 21 Sep 21 Oct 22 Nov 22 Dec 23
Date
29. Vireo cassinii
Latitude
20
30
40
50
Jan 16 Feb 16 Mar 19 Apr 19 May 20 Jun 20 Jul 21 Aug 21 Sep 21 Oct 22 Nov 22 Dec 23
Date
27. Vireo vicinior
Latitude
25
30
35
Jan 16 Feb 16 Mar 19 Apr 19 May 20 Jun 20 Jul 21 Aug 21 Sep 21 Oct 22 Nov 22 Dec 23
Date
30. Vireo solitarius
Latitude
30
40
50
60
Jan 16 Feb 16 Mar 19 Apr 19 May 20 Jun 20 Jul 21 Aug 21 Sep 21 Oct 22 Nov 22 Dec 23
Date

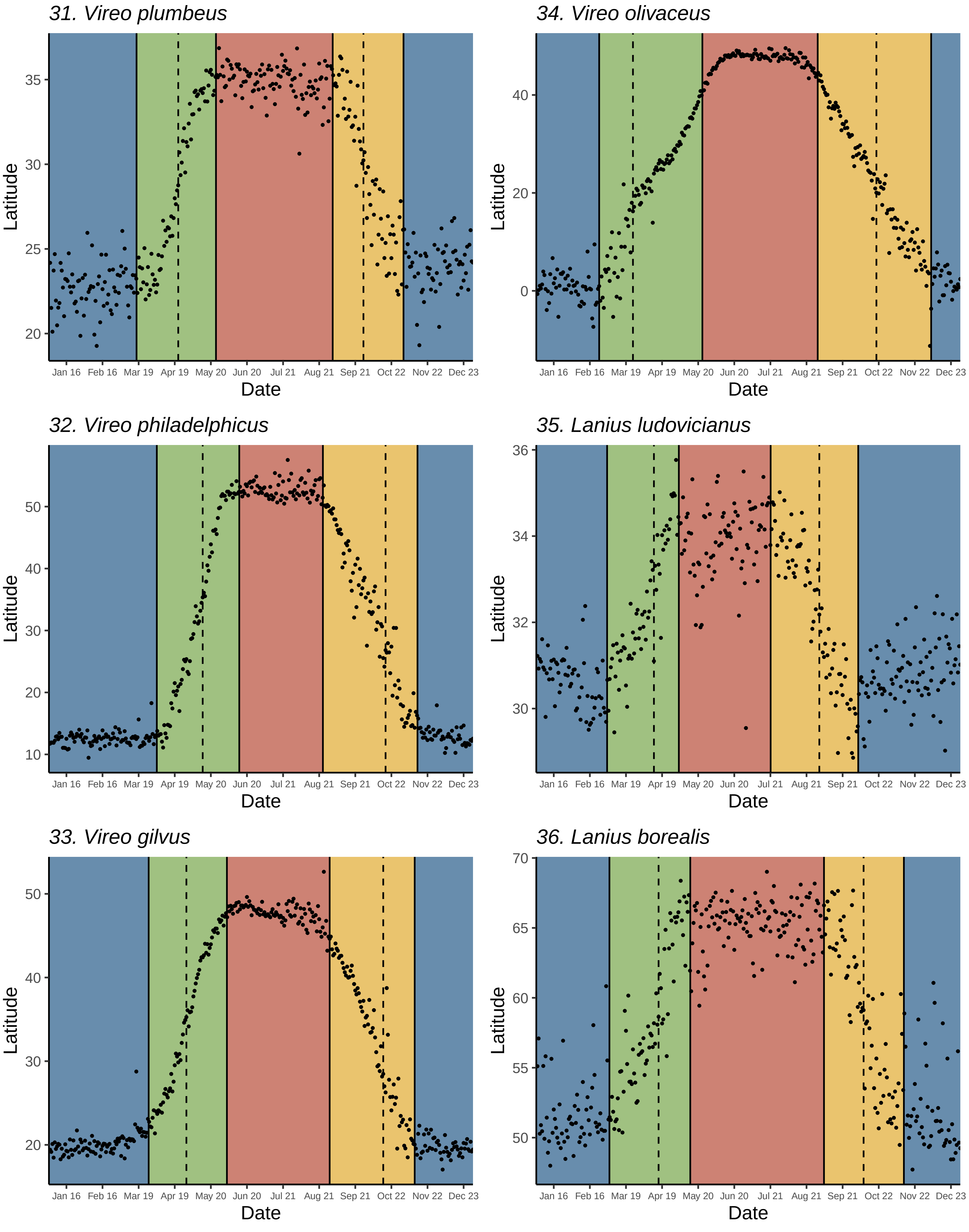
31. Vireo plumbeus
34. Vireo olivaceus
32. Vireo philadelphicus
35. Lanius ludovicianus
33. Vireo gilvus
36. Lanius borealis
Latitude
Date
Jan 16
Feb 16
Mar 19
Apr 19
May 20
Jun 20
Jul 21
Aug 21
Sep 21
Oct 22
Nov 22
Dec 23

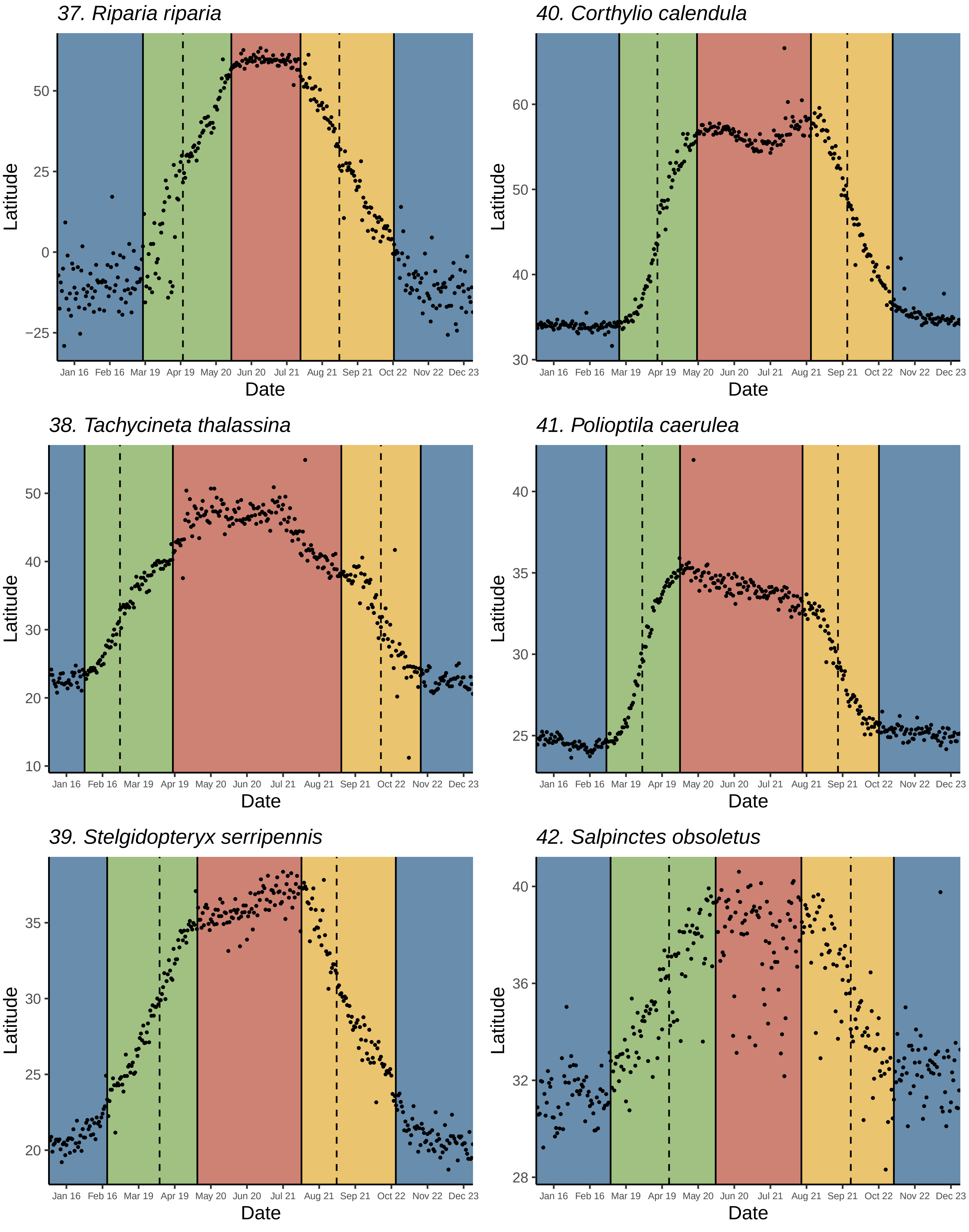
37. Riparia riparia
40. Corthylio calendula
38. Tachycineta thalassina
41. Polioptila caerulea
39. Stelgidopteryx serripennis
42. Salpinctes obsoletus
Latitude
Date
Jan 16
Feb 16
Mar 19
Apr 19
May 20
Jun 20
Jul 21
Aug 21
Sep 21
Oct 22
Nov 22
Dec 23

*43. Troglodytes hiemalis*

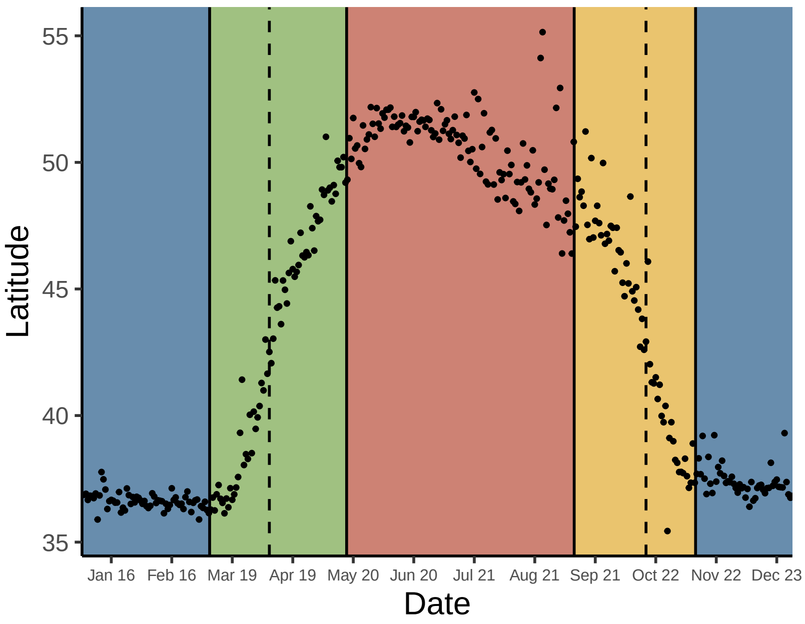


*46. Dumetella carolinensis*

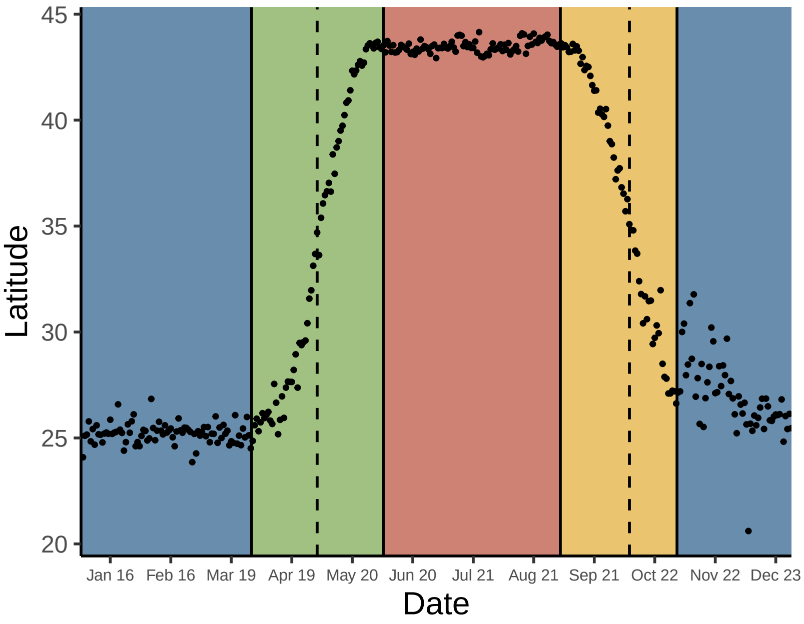


*44. Cistothorus stellaris*

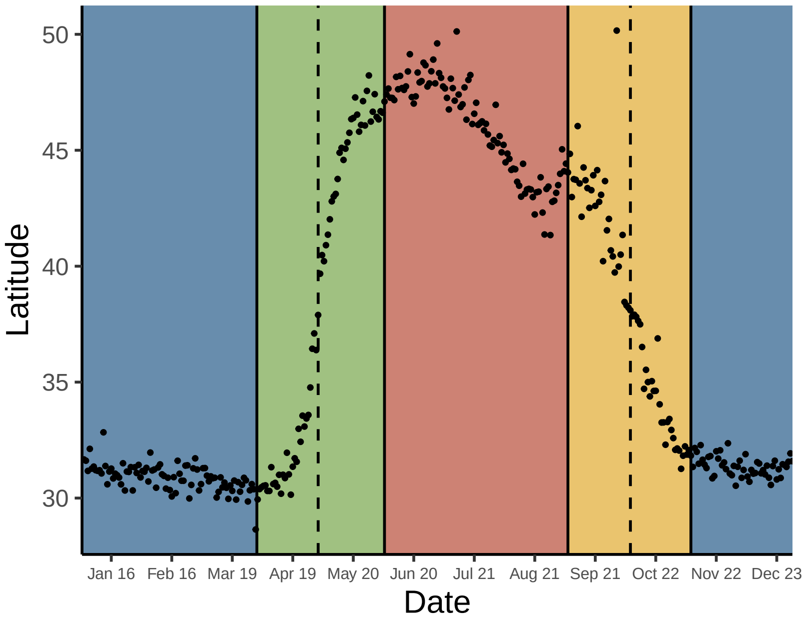


*47. Toxostoma rufum*

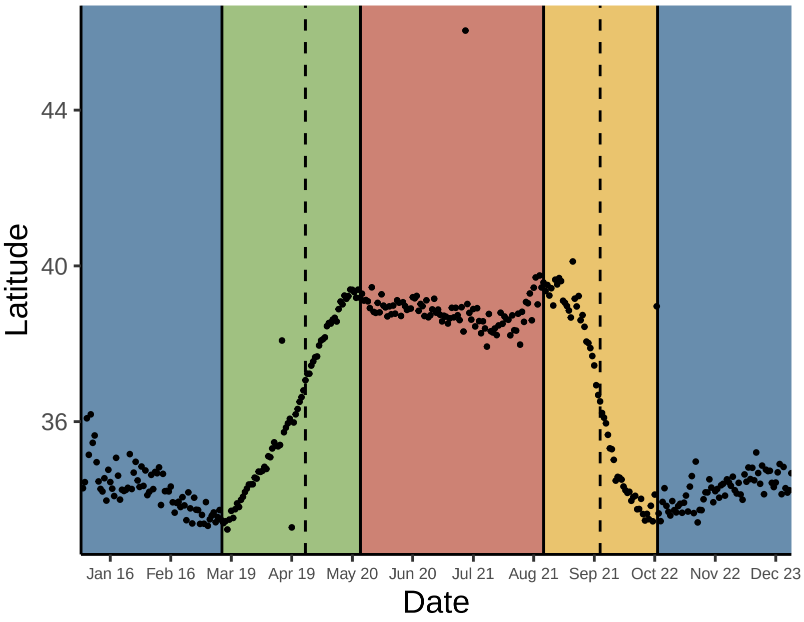


*45. Cistothorus palustris*

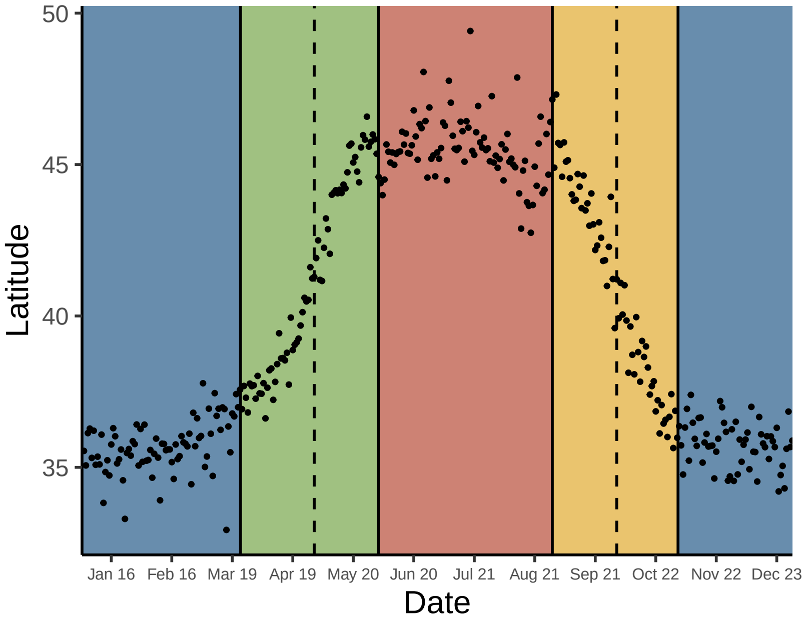


*48. Oreoscoptes montanus*

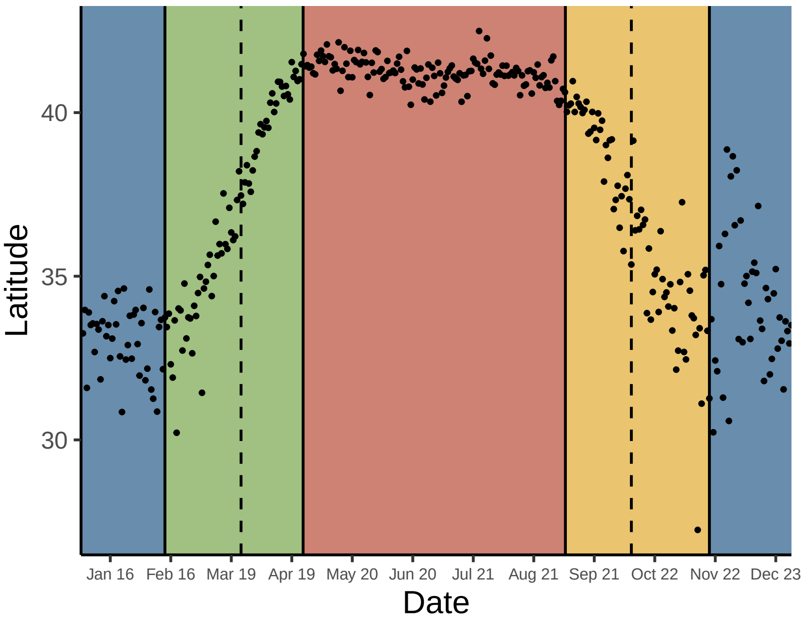

49. *Sialia currucoides*

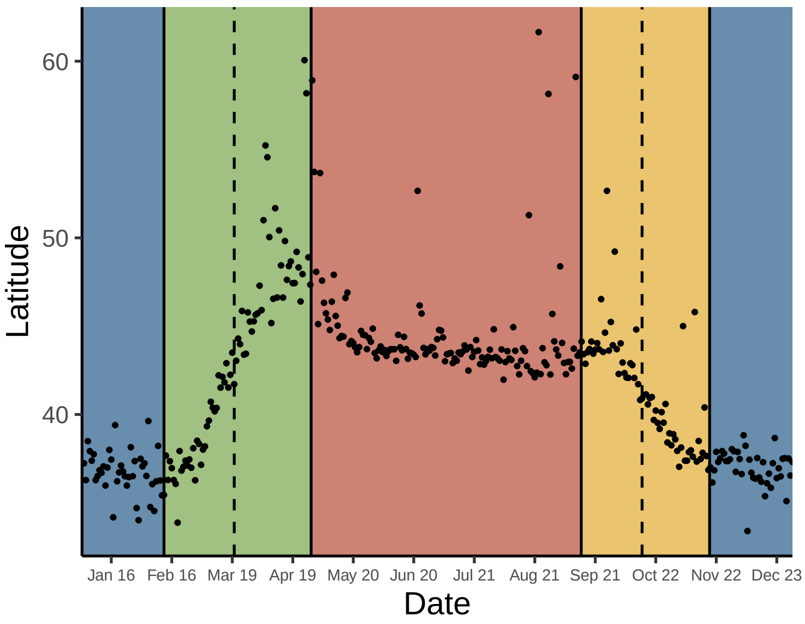


52. *Catharus ustulatus*

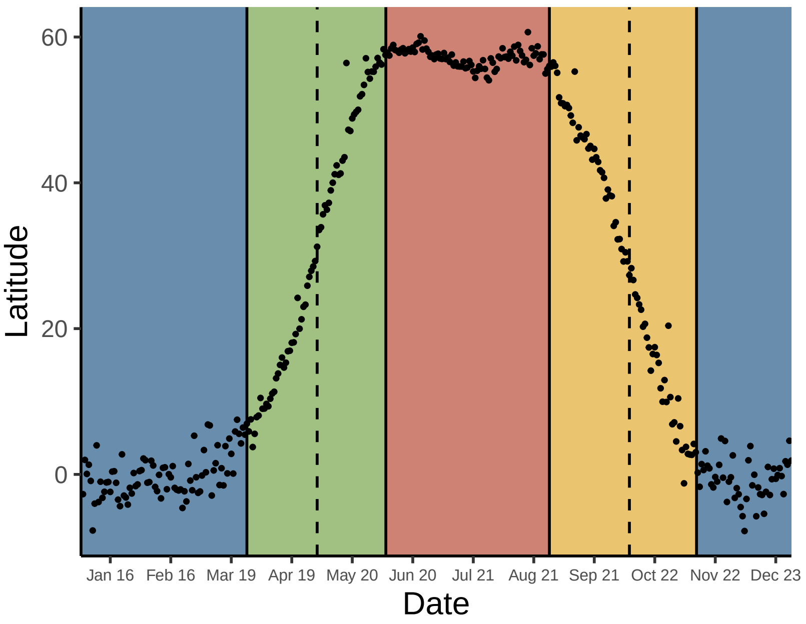


50. *Catharus minimus*

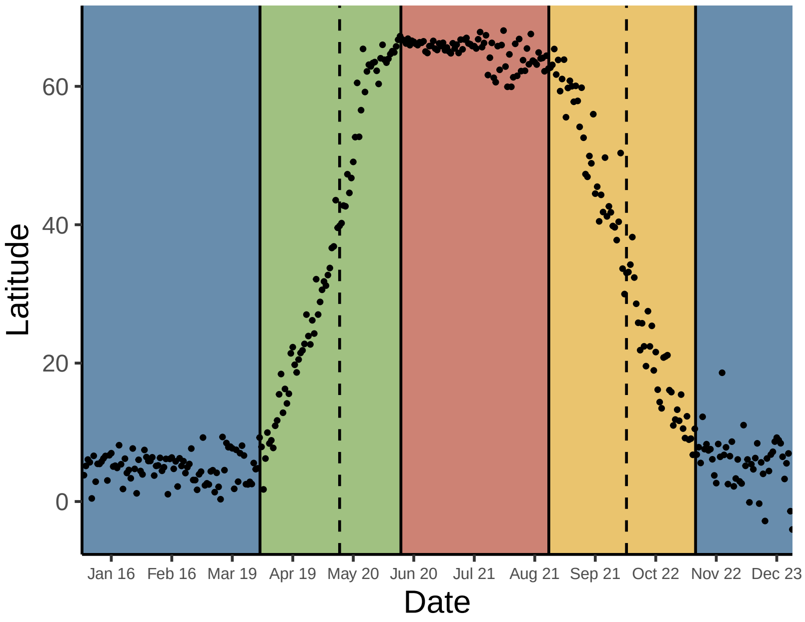


53. *Catharus guttatus*

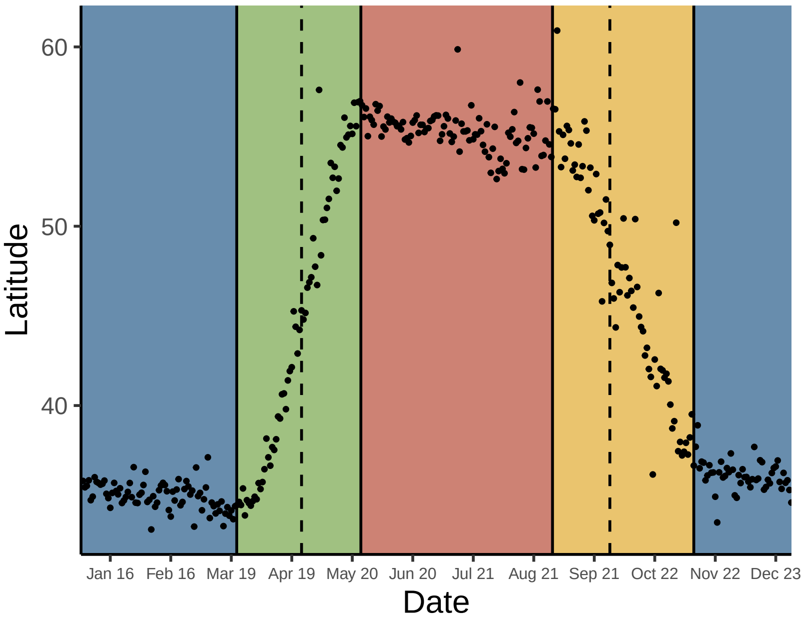


51. *Catharus bicknelli*

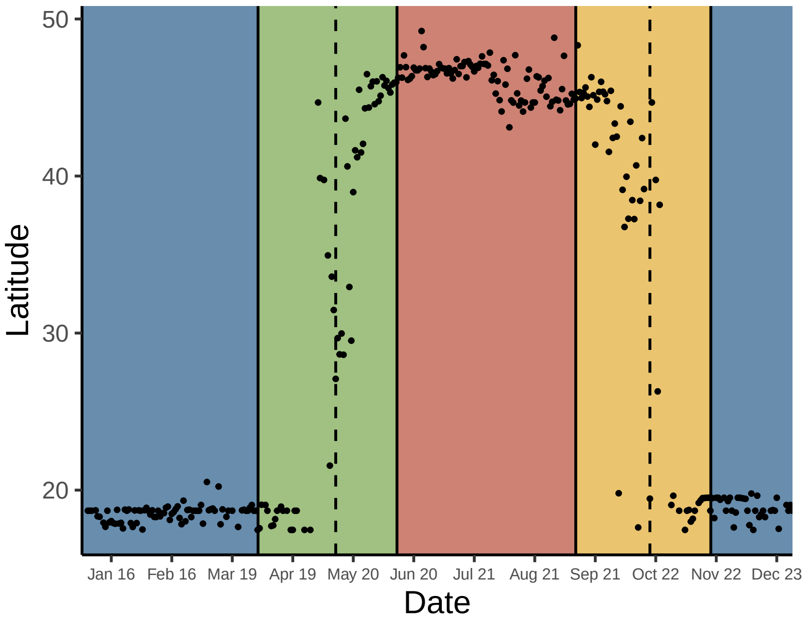


54. *Hylocichla mustelina*

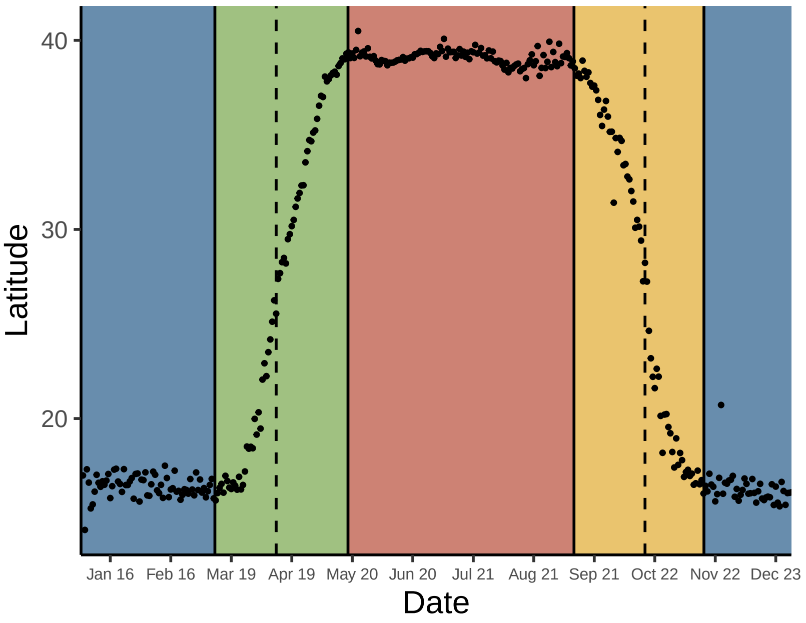

Figure S3 page 10 of 25

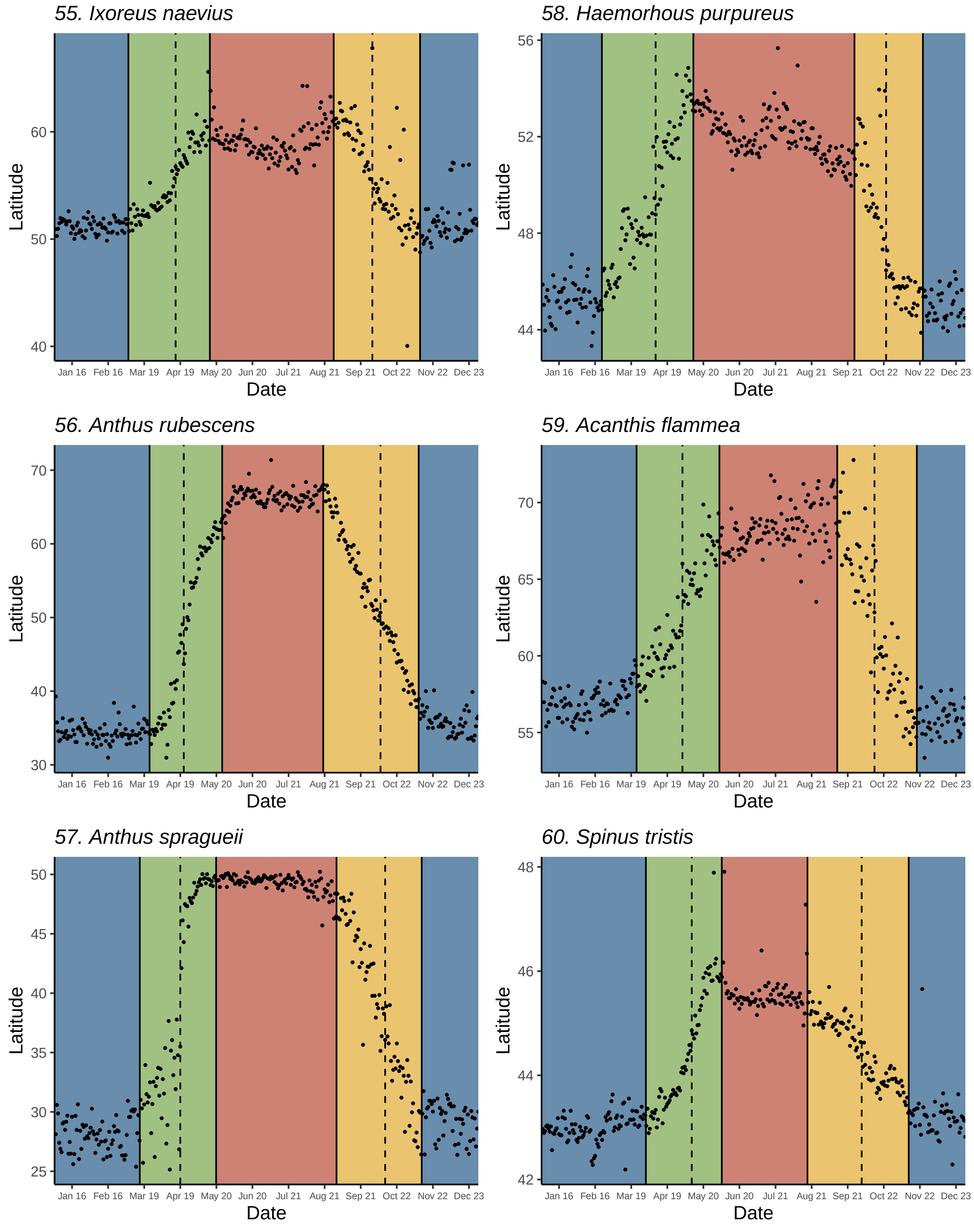

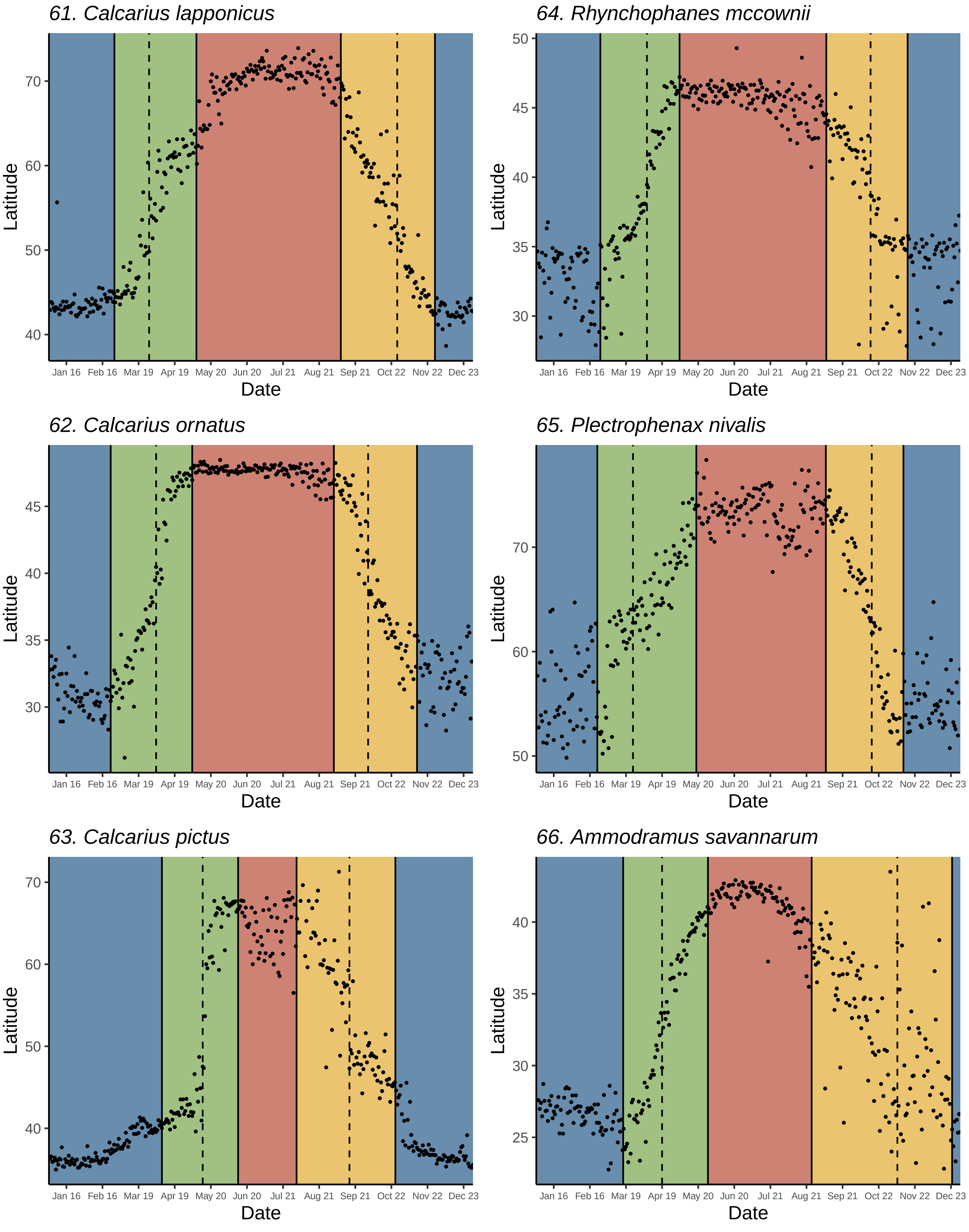
61. Calcarius lapponicus
64. Rhynchophanes mccownii
62. Calcarius ornatus
65. Plectrophenax nivalis
63. Calcarius pictus
66. Ammodramus savannarum
Latitude
Date
Jan 16
Feb 16
Mar 19
Apr 19
May 20
Jun 20
Jul 21
Aug 21
Sep 21
Oct 22
Nov 22
Dec 23

*67. Chondestes grammacus*

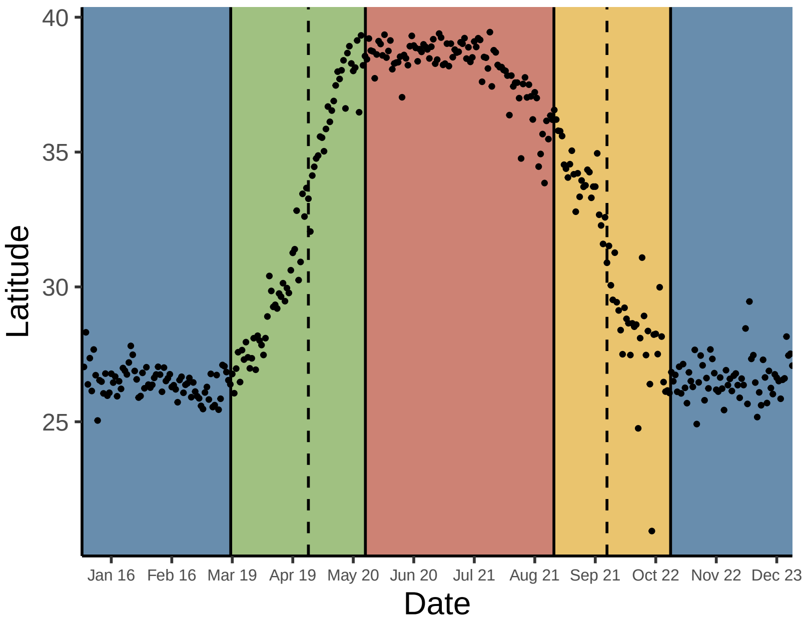


*70. Spizella pusilla*

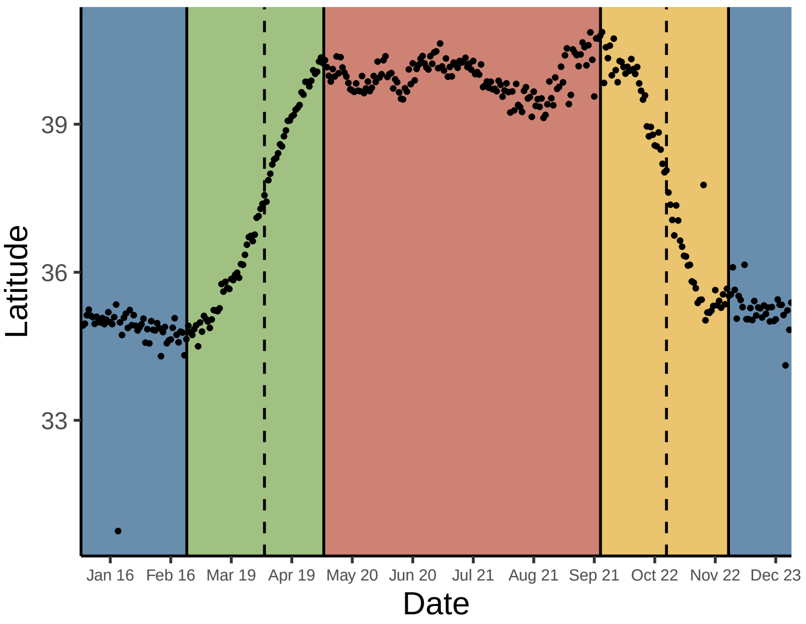


*68. Calamospiza melanocorys*

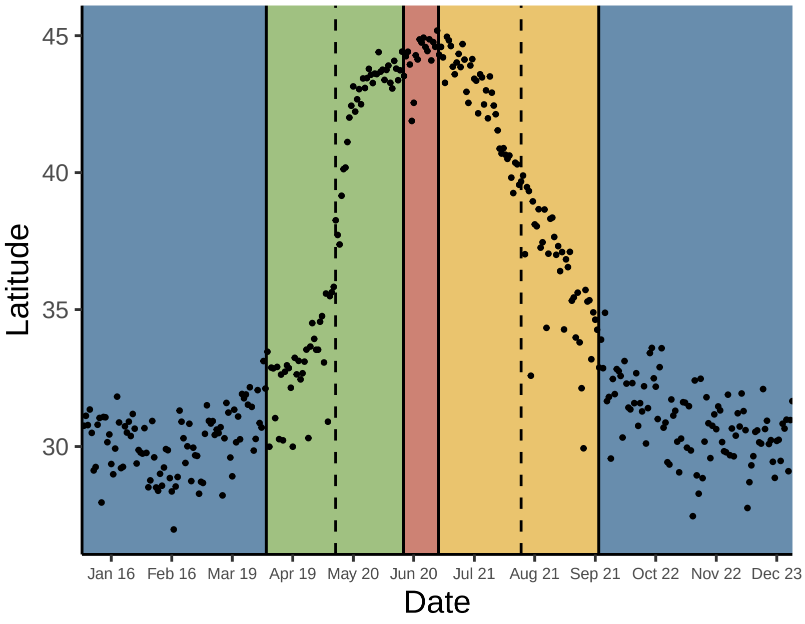


*71. Spizella breweri*

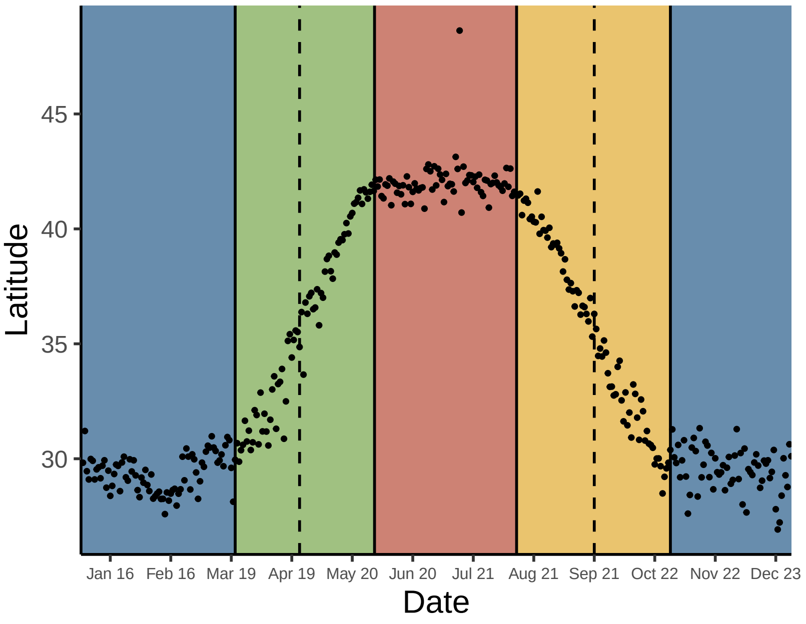


*69. Spizella pallida*

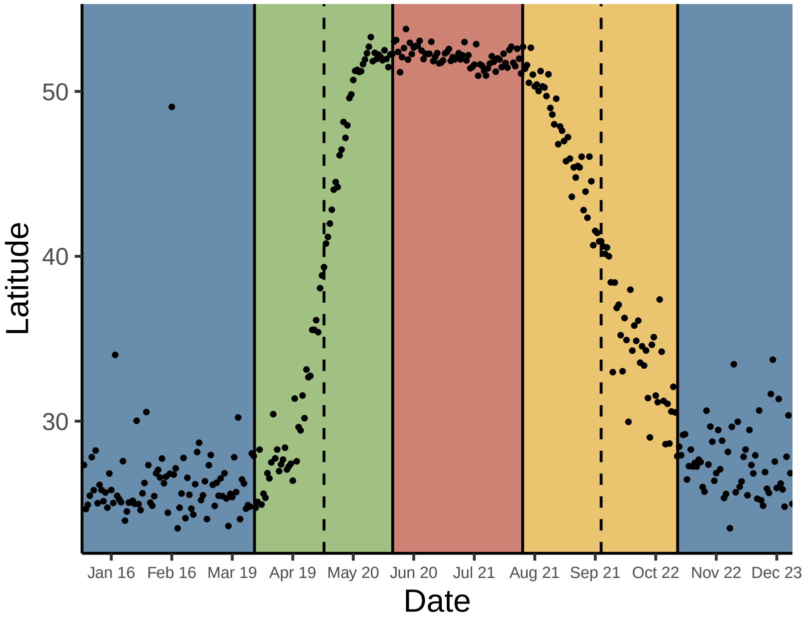


*72. Passerella iliaca*

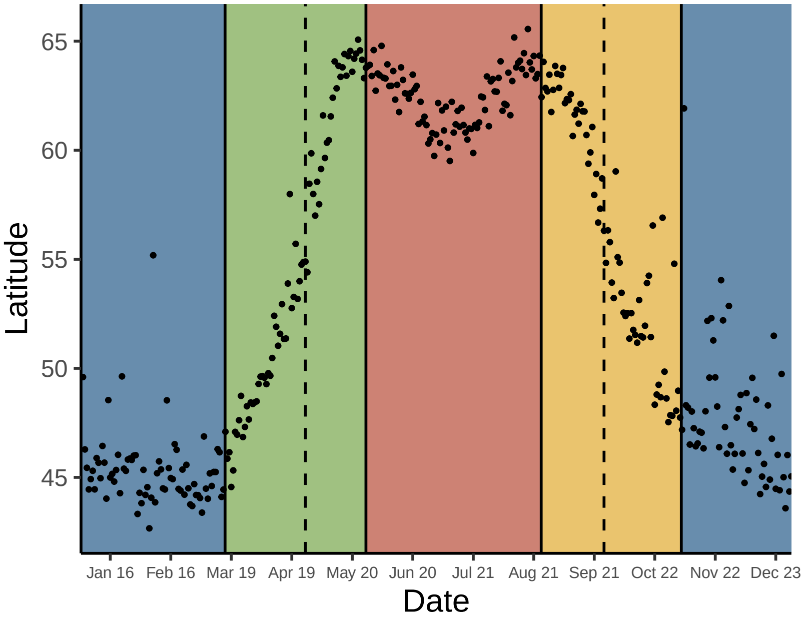

*73. Spizelloides arborea*

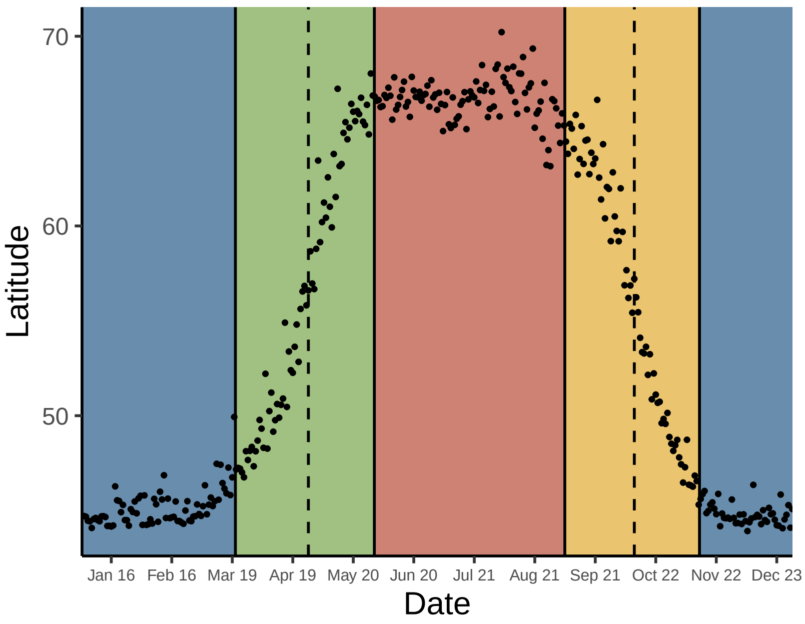


*74. Junco hyemalis*

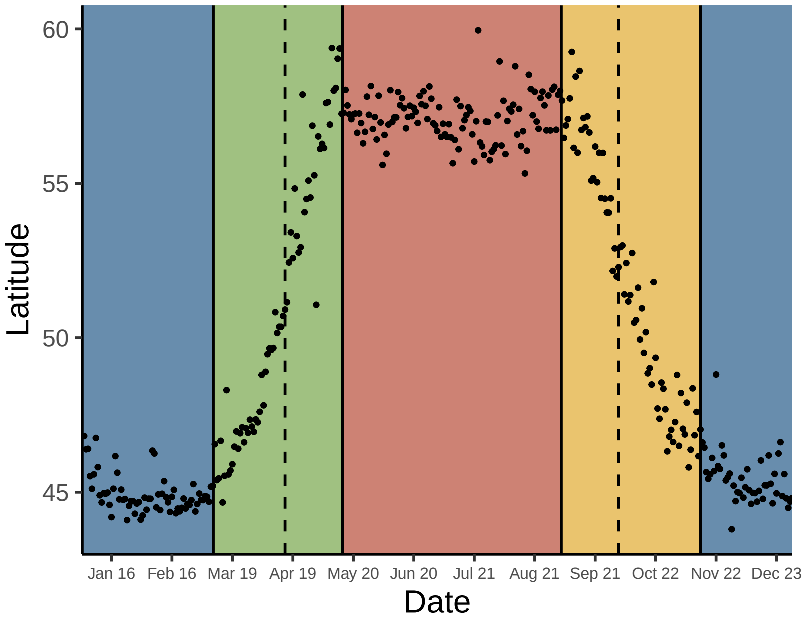


*75. Zonotrichia leucophrys*

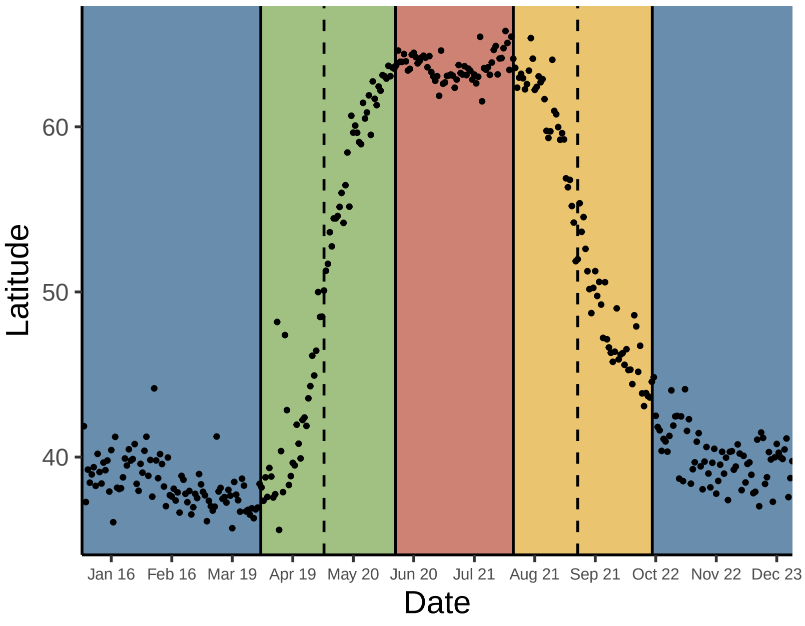


*76. Zonotrichia atricapilla*

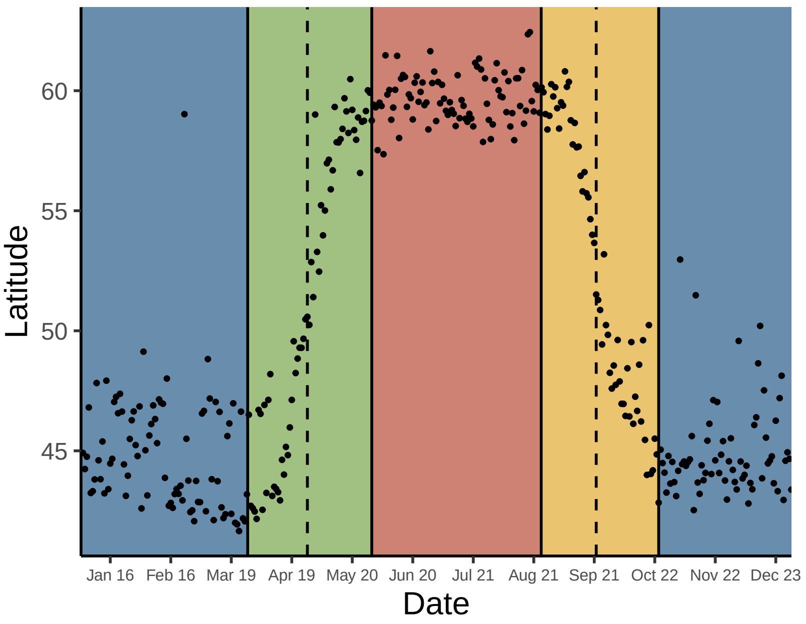


*77. Artemisiospiza nevadensis*

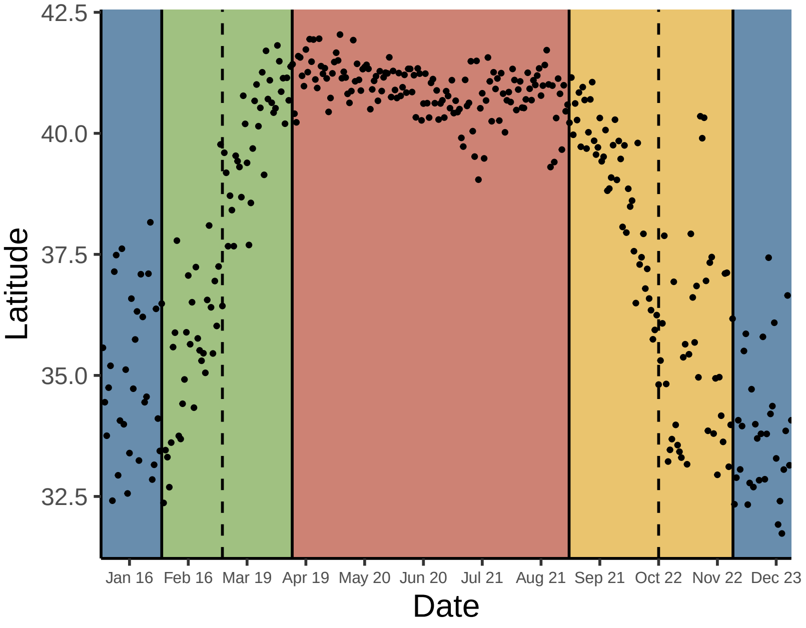


*78. Pooecetes gramineus*

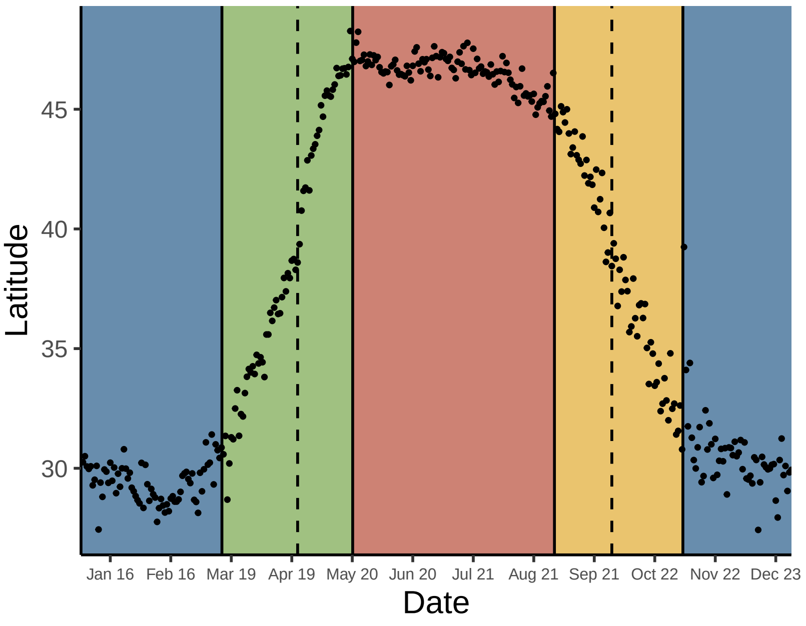

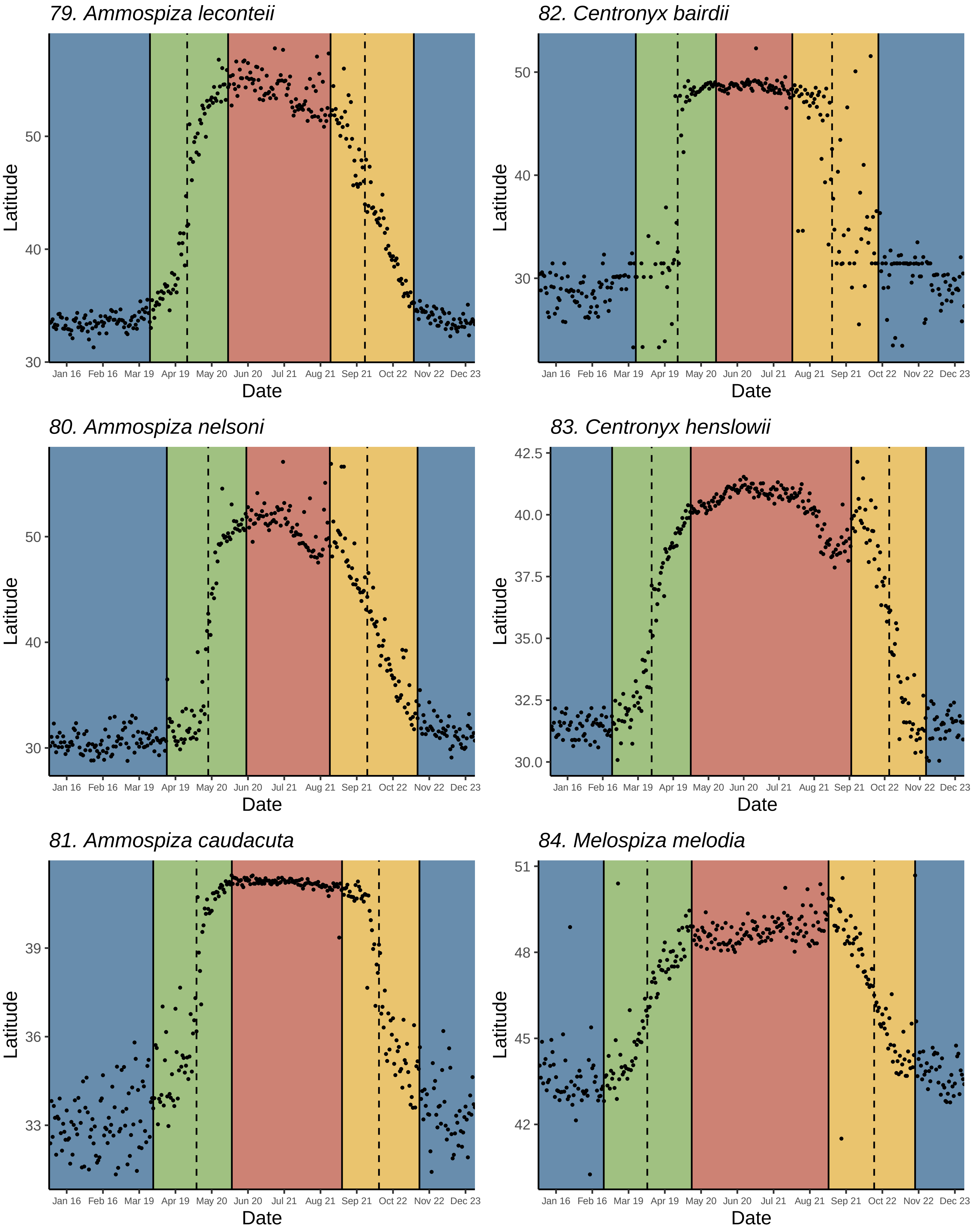
79. Ammospiza leconteii
82. Centronyx bairdii
80. Ammospiza nelsoni
83. Centronyx henslowii
81. Ammospiza caudacuta
84. Melospiza melodia
Latitude
Date
Jan 16 Feb 16 Mar 19 Apr 19 May 20 Jun 20 Jul 21 Aug 21 Sep 21 Oct 22 Nov 22 Dec 23
30
40
50
30.0
32.5
35.0
37.5
40.0
42.5
33
36
39
42
45
48
51

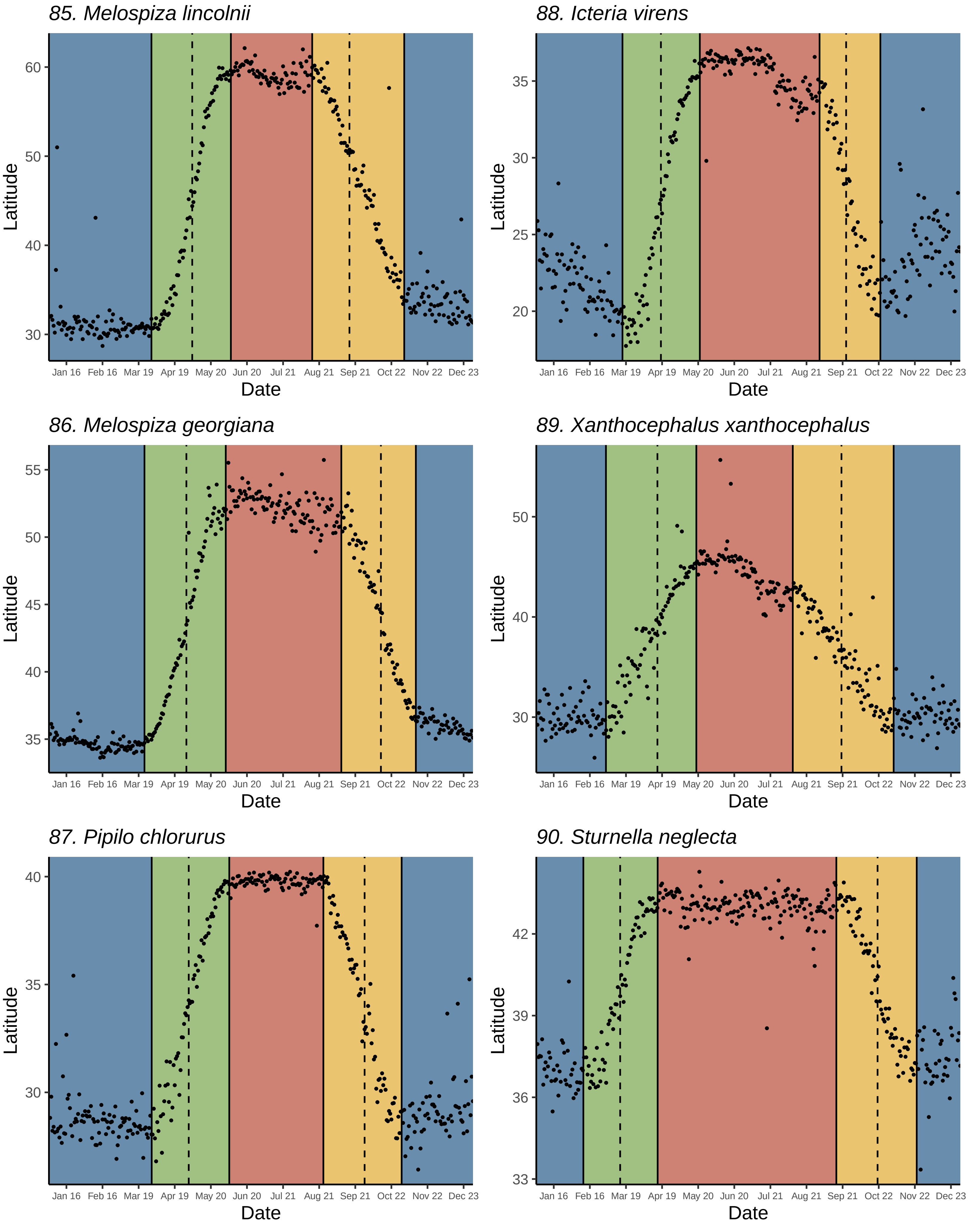
85. Melospiza lincolnii
88. Icteria virens
86. Melospiza georgiana
89. Xanthocephalus xanthocephalus
87. Pipilo chlorurus
90. Sturnella neglecta
Latitude
Date
Jan 16
Feb 16
Mar 19
Apr 19
May 20
Jun 20
Jul 21
Aug 21
Sep 21
Oct 22
Nov 22
Dec 23

*91. Icterus spurius*

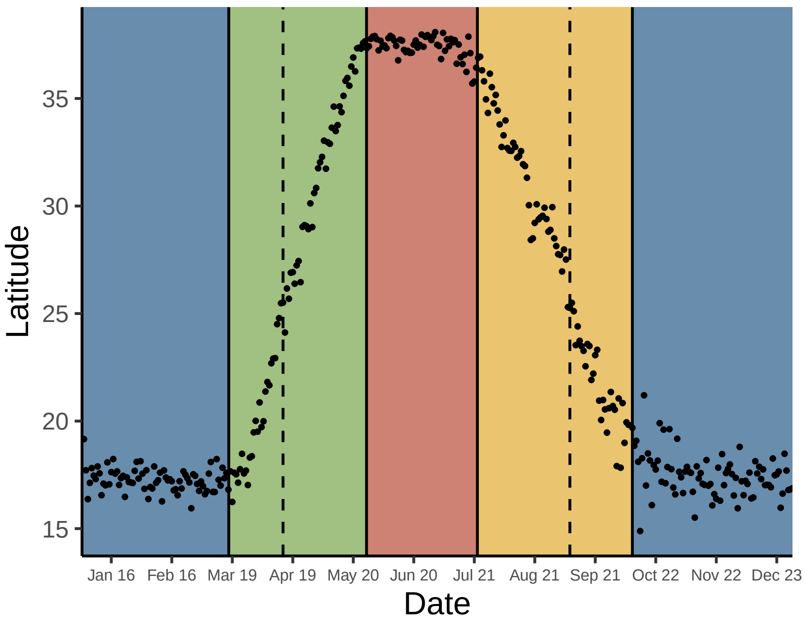


*92. Icterus cucullatus*

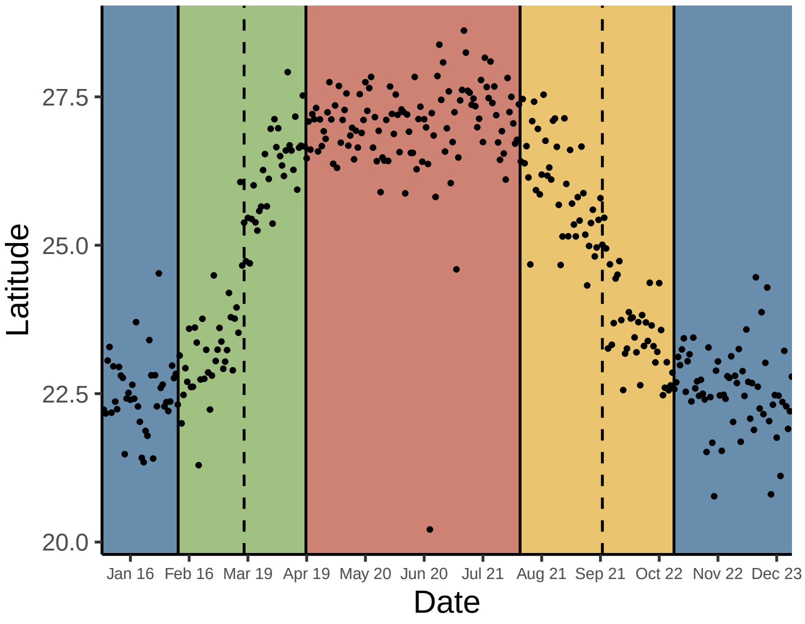


*93. Icterus bullockii*

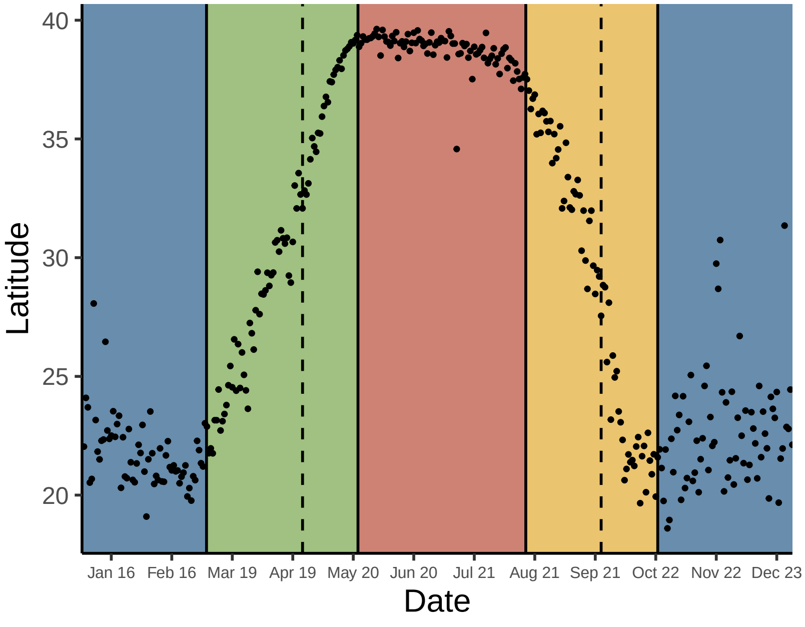


*94. Icterus galbula*

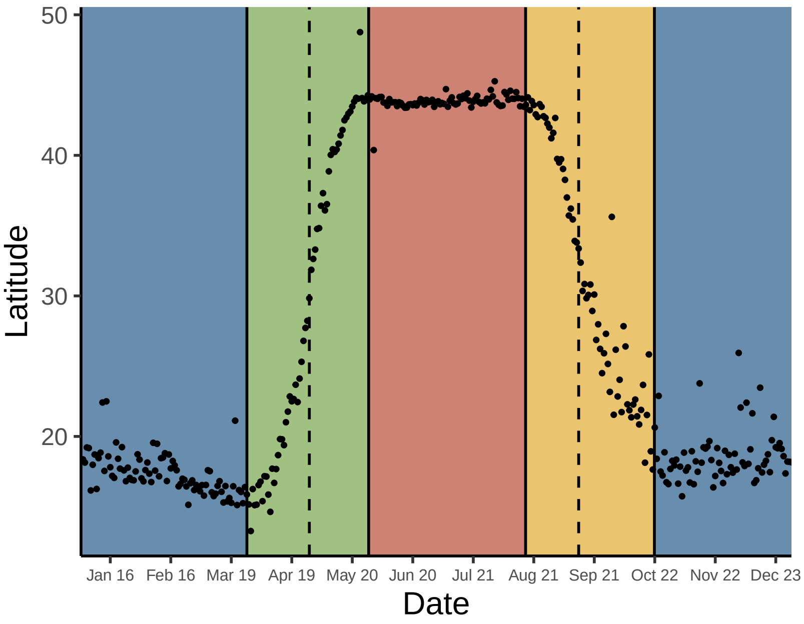


*95. Icterus parisorum*

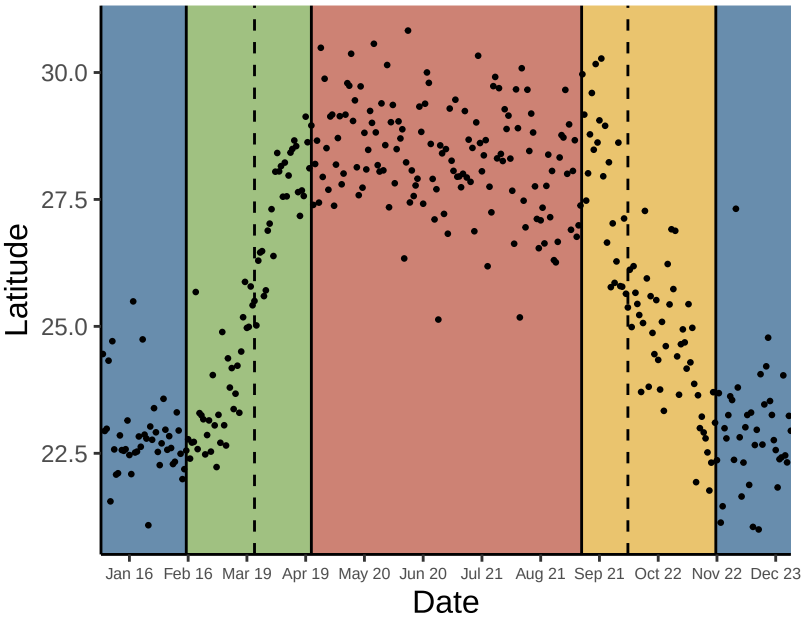


*96. Molothrus ater*

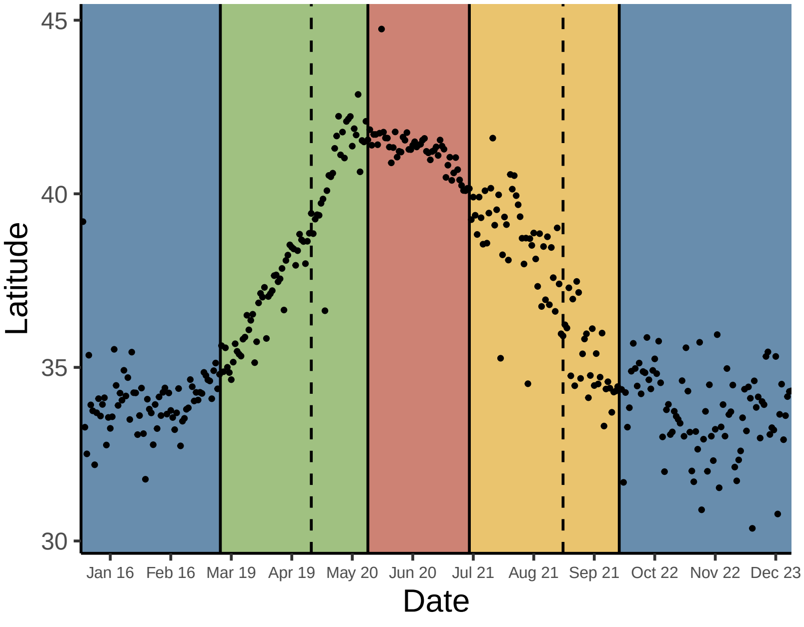

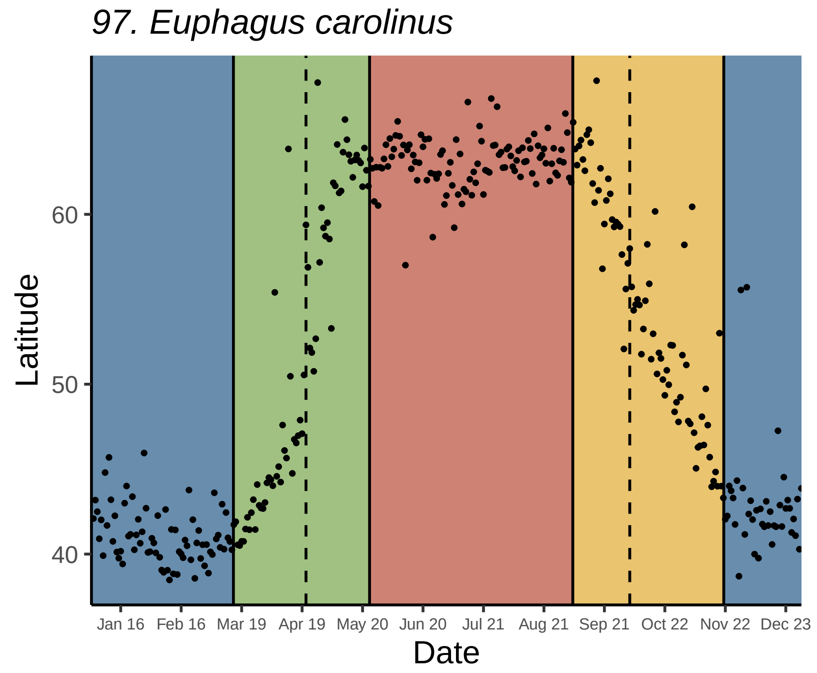
97. Euphagus carolinus
Latitude
Date
Jan 16
Feb 16
Mar 19
Apr 19
May 20
Jun 20
Jul 21
Aug 21
Sep 21
Oct 22
Nov 22
Dec 23

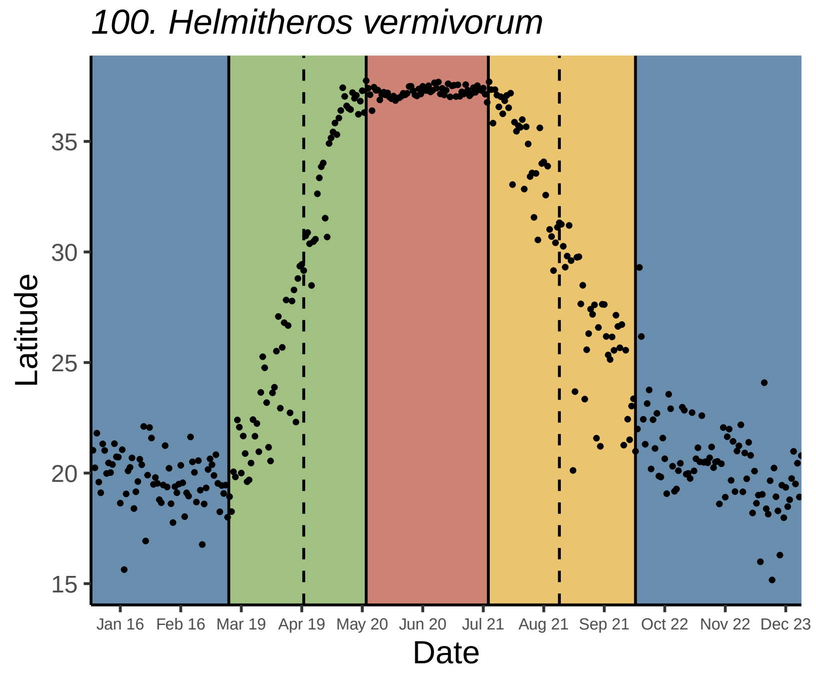
100. Helmitheros vermivorum
Latitude
Date

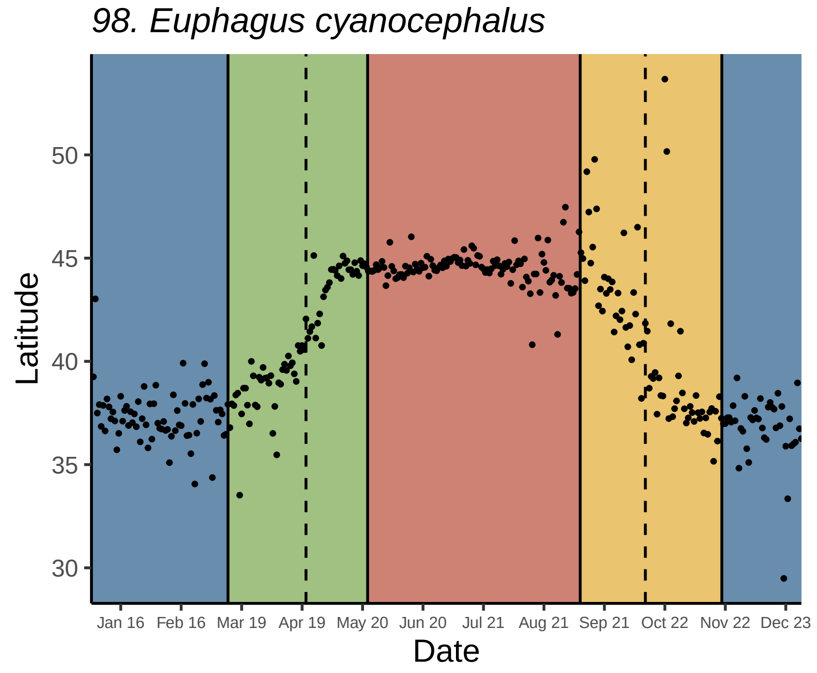
98. Euphagus cyanocephalus
Latitude
Date

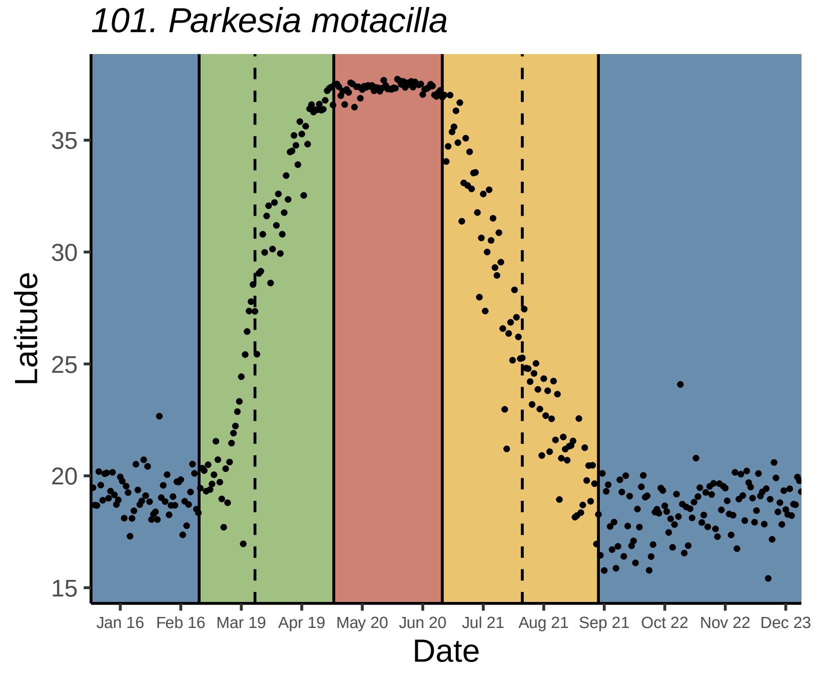
101. Parkesia motacilla
Latitude
Date

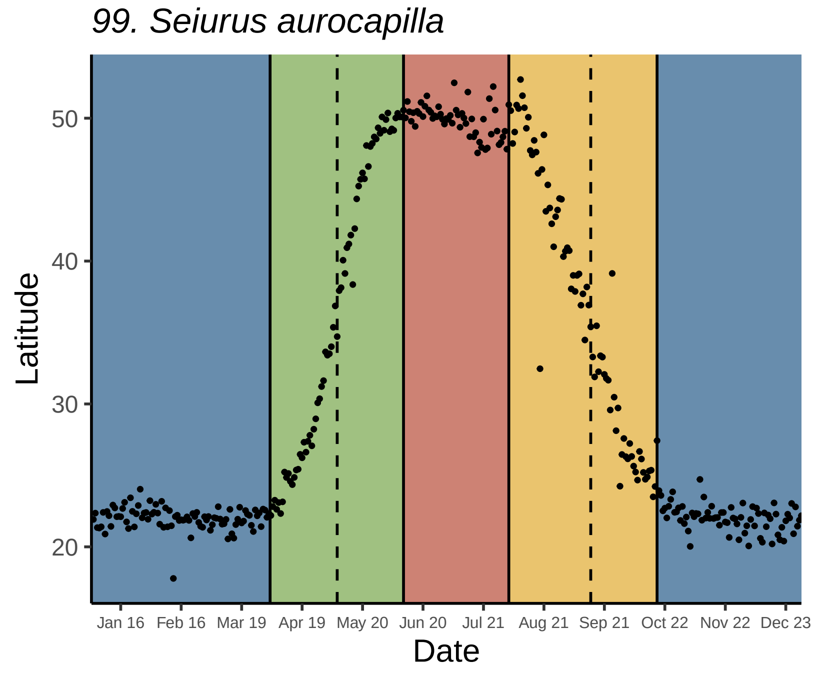
99. Seiurus aurocapilla
Latitude
Date

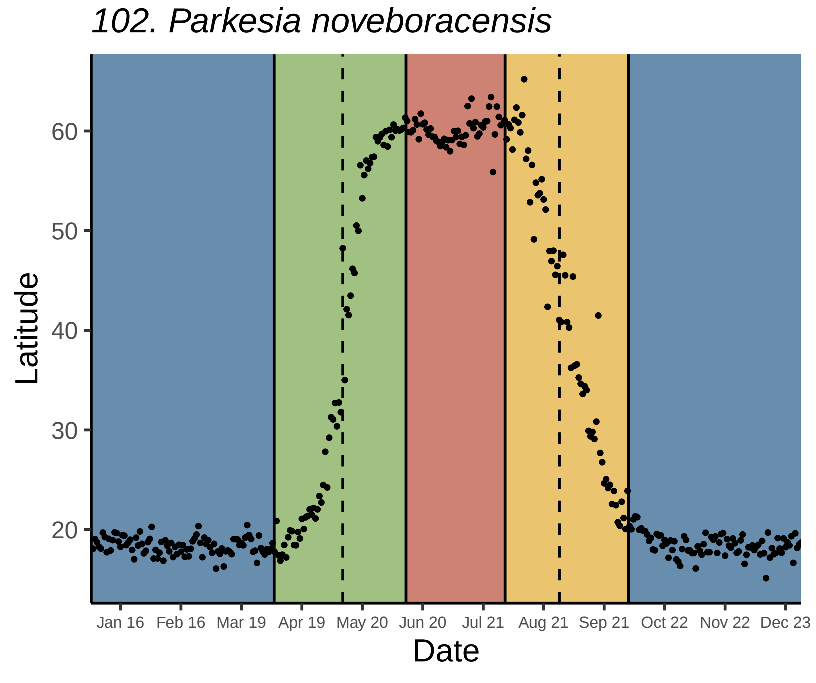
102. Parkesia noveboracensis
Latitude
Date

*103. Vermivora chrysoptera*

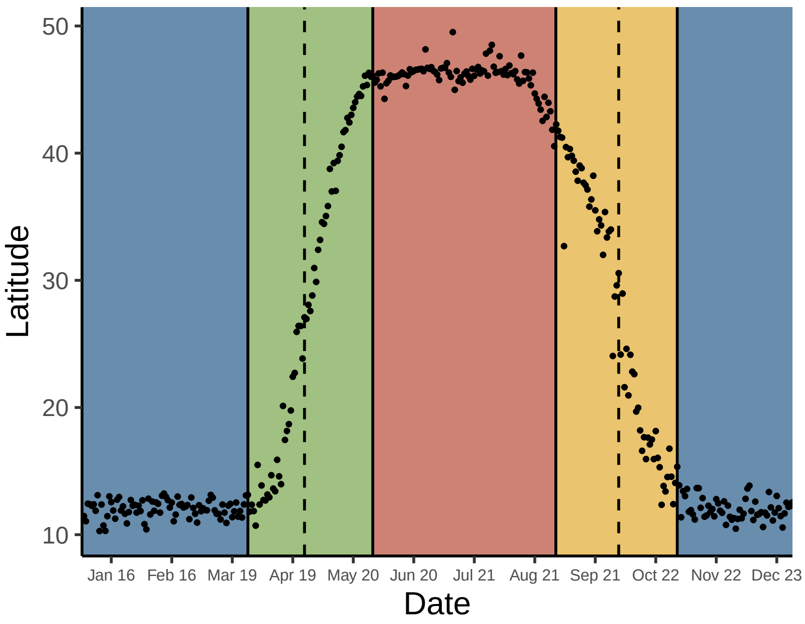


*106. Protonotaria citrea*

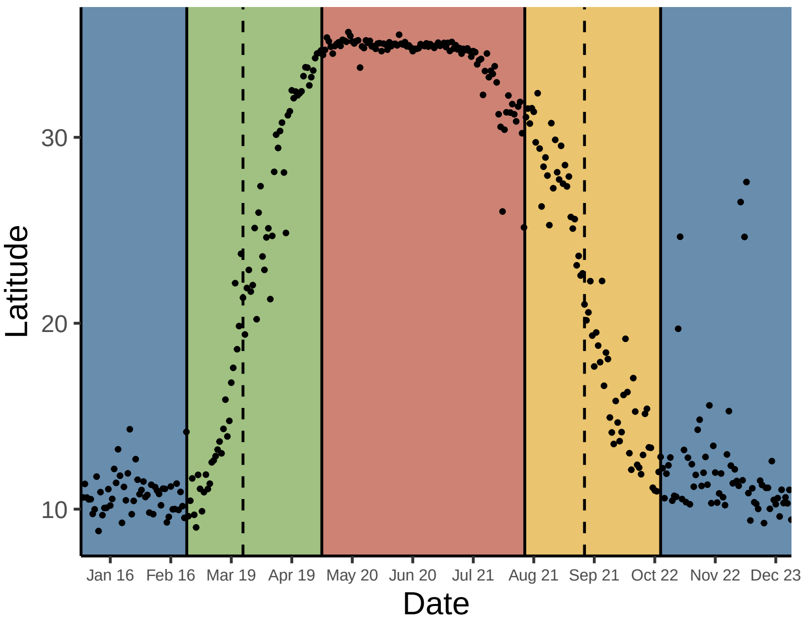


*104. Vermivora cyanoptera*

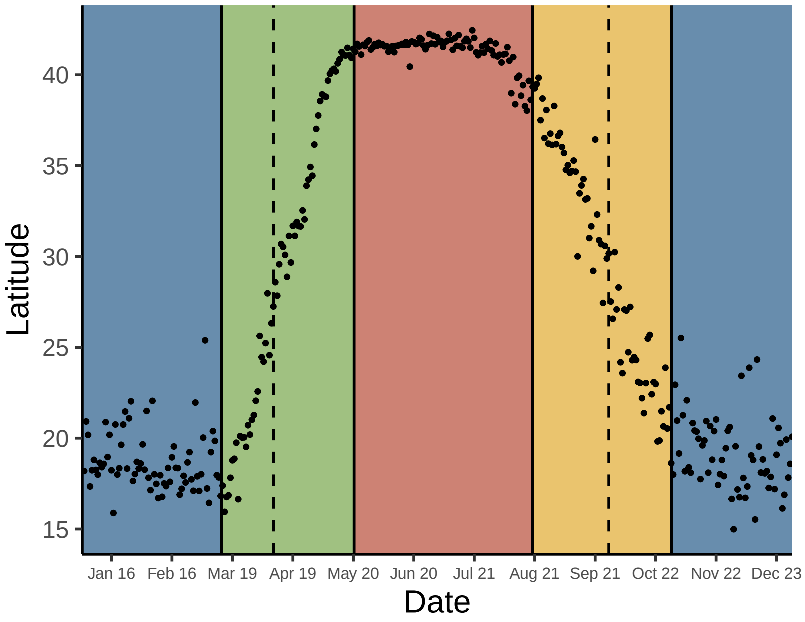


*107. Limnothlypis swainsonii*

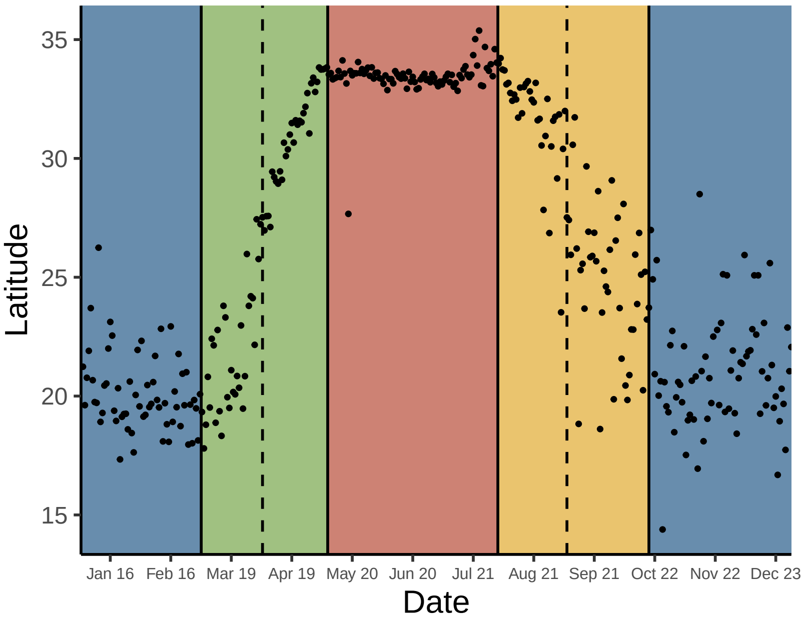


*105. Mniotilta varia*

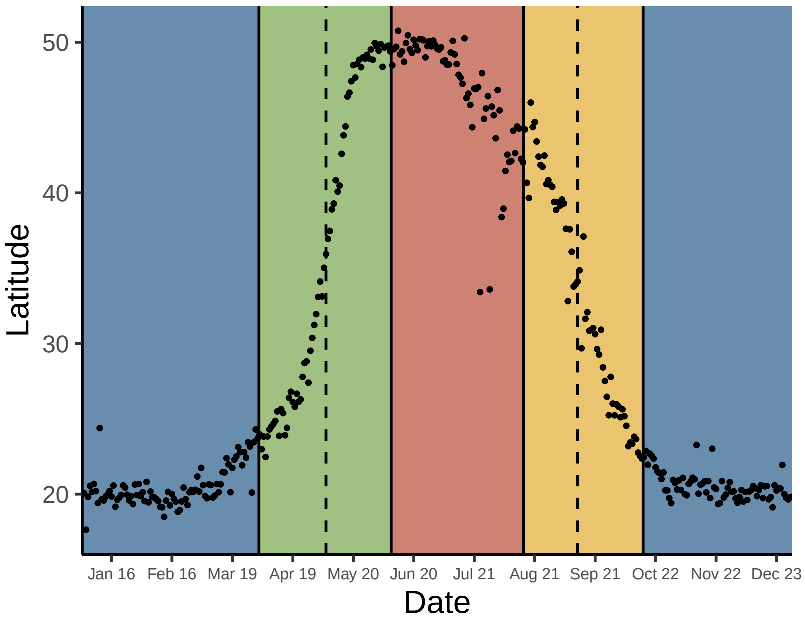


*108. Leiothlypis peregrina*

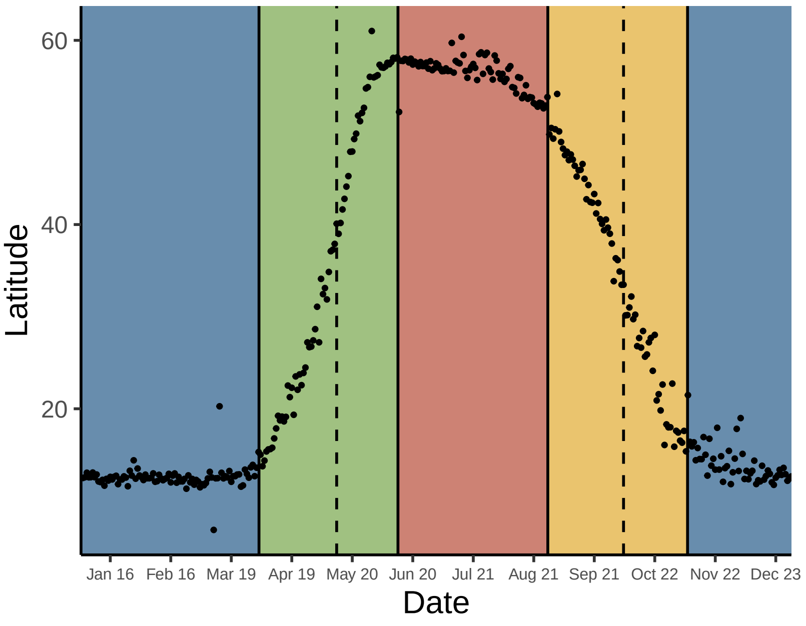

*109. Leiothlypis celata*

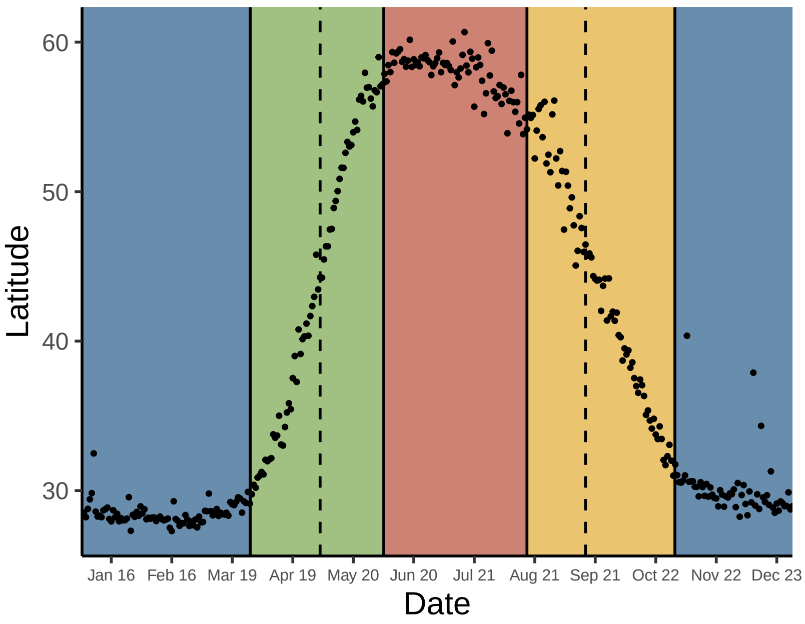


*112. Leiothlypis virginiae*

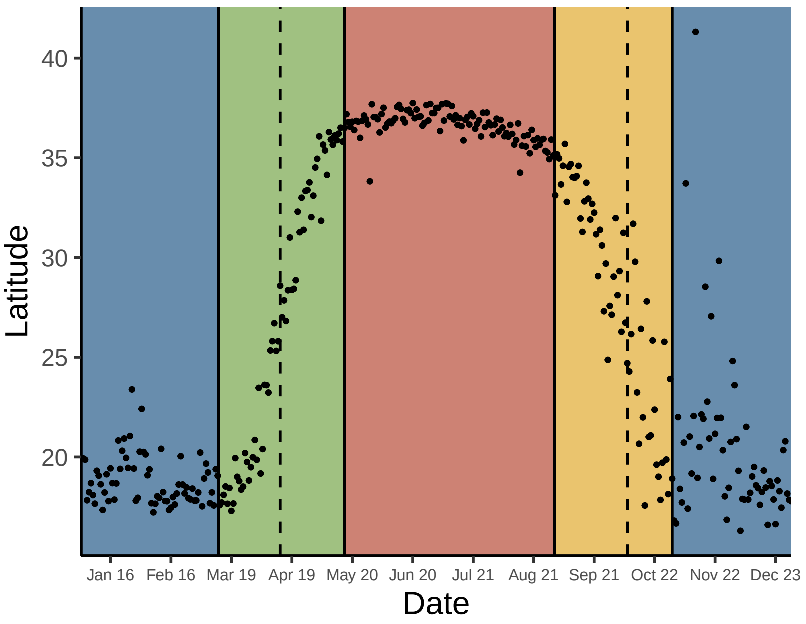


*110. Leiothlypis luciae*

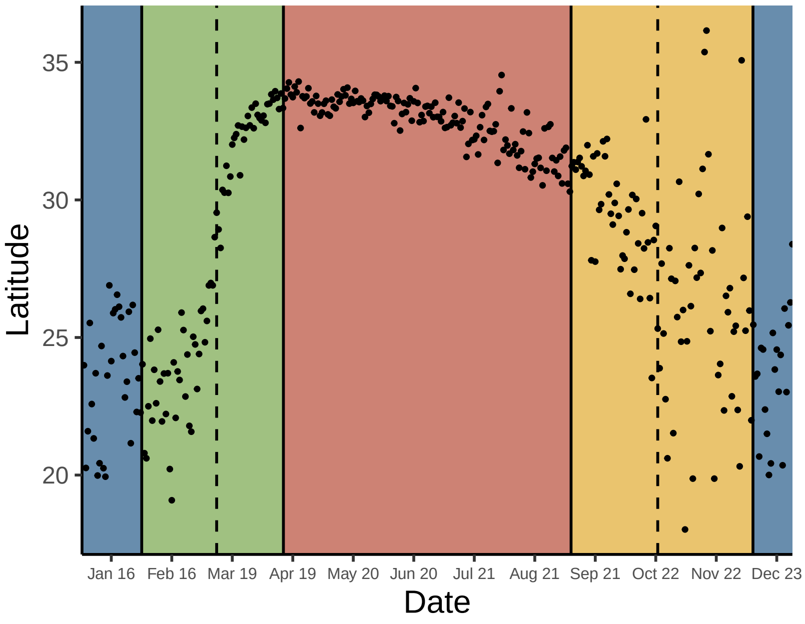


*113. Oporornis agilis*

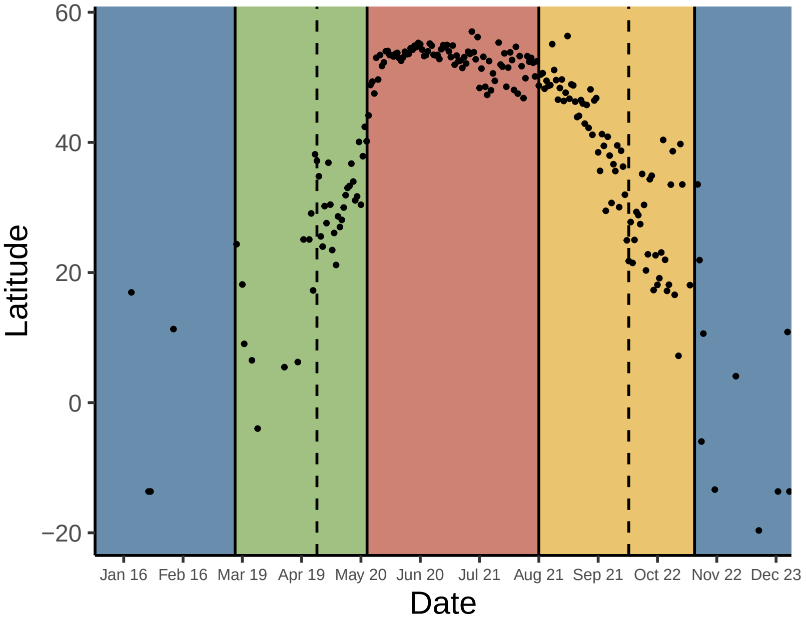


*111. Leiothlypis ruficapilla*

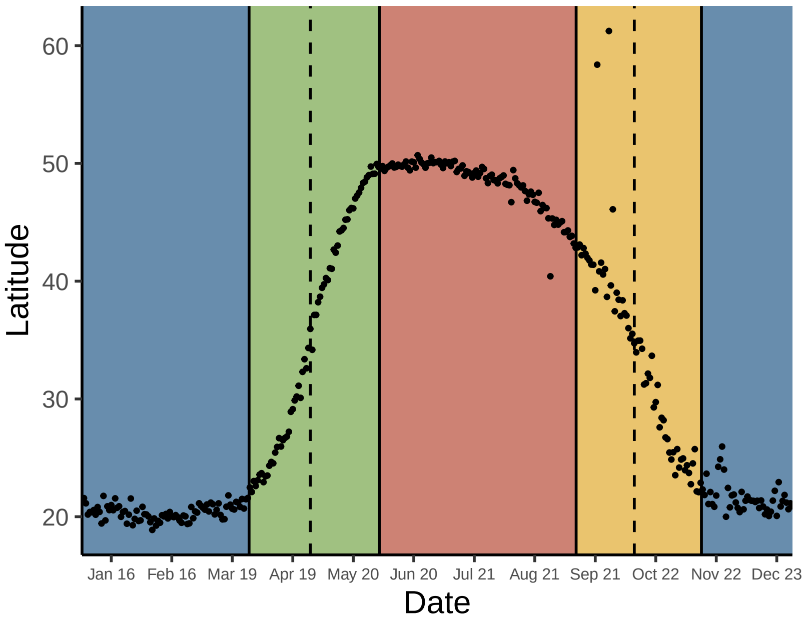


*114. Geothlypis tolmiei*

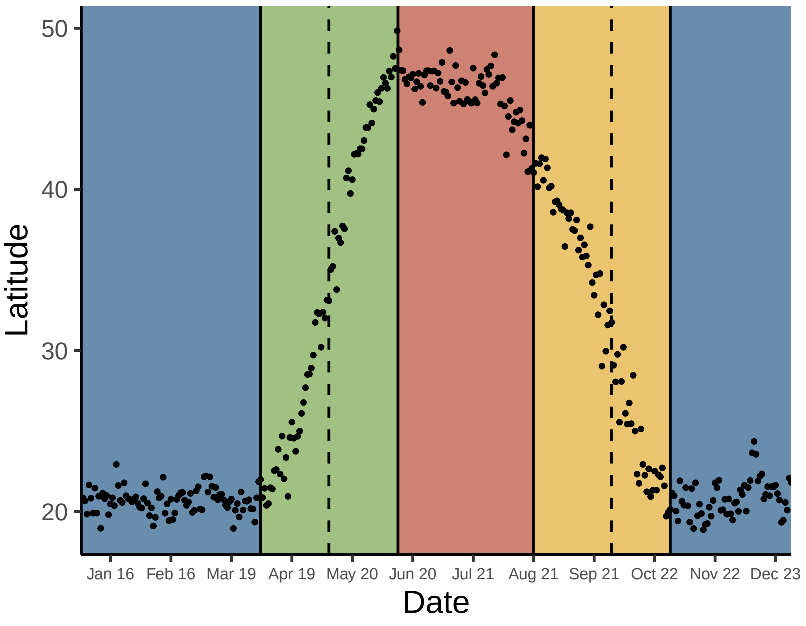

*115. Geothlypis philadelphia*

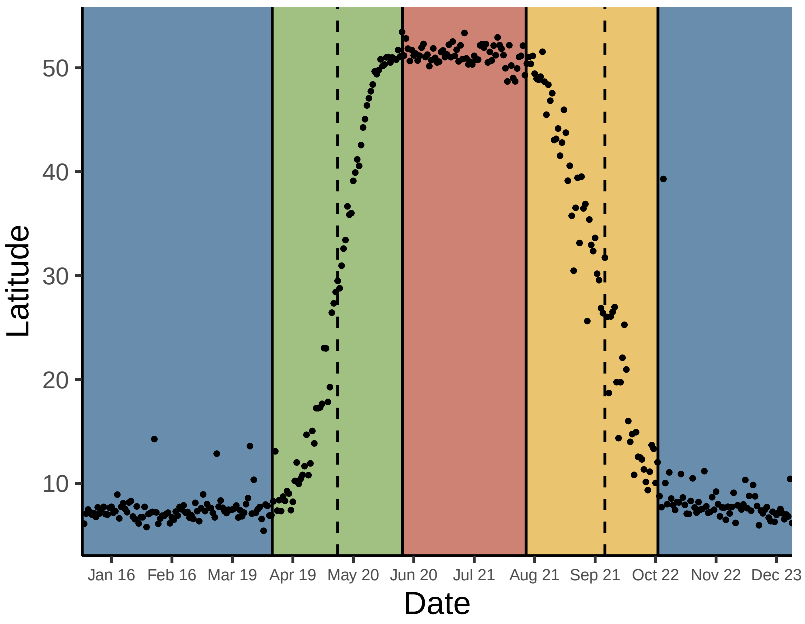


*118. Setophaga ruticilla*

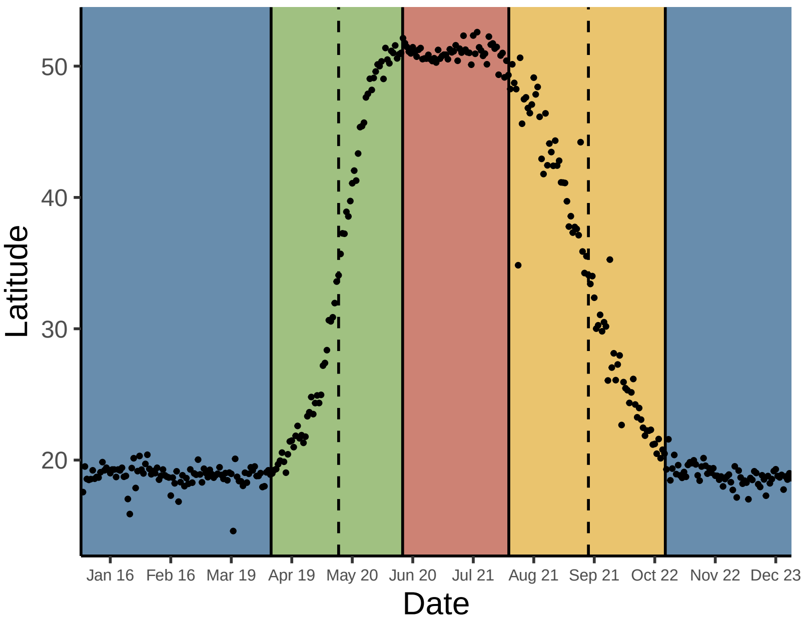


*116. Geothlypis formosa*

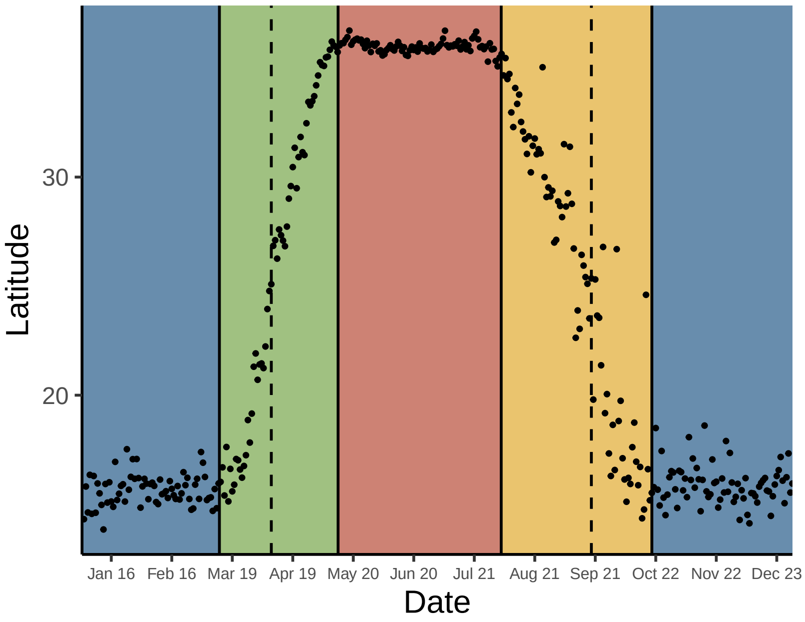


*119. Setophaga kirtlandii*

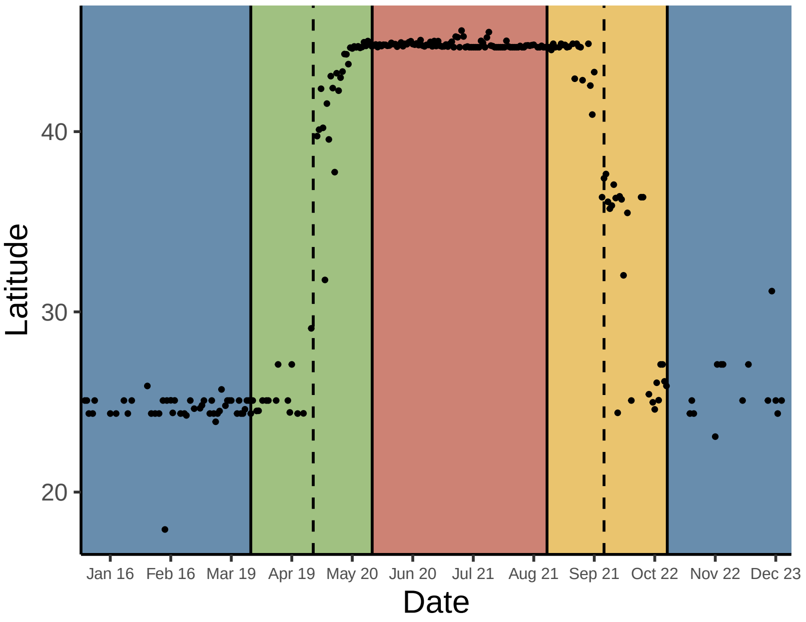


*117. Setophaga citrina*

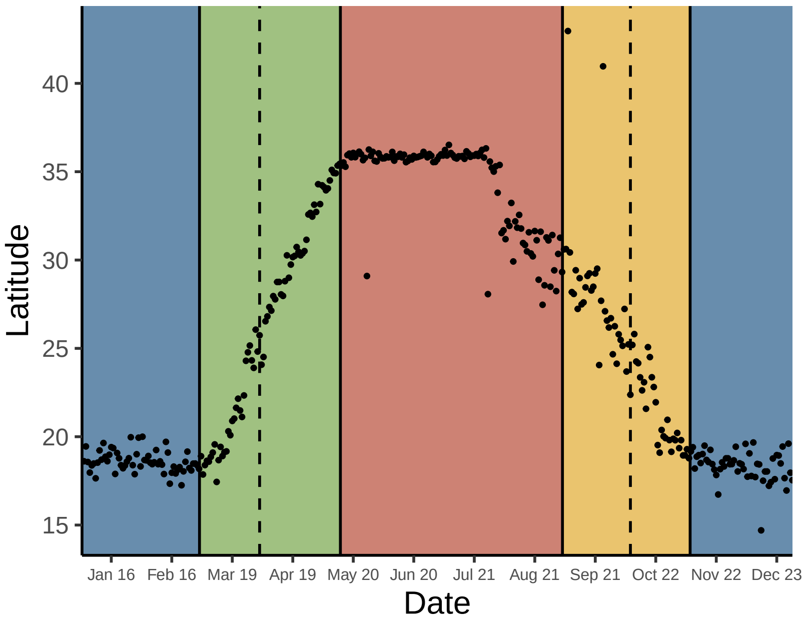


*120. Setophaga tigrina*

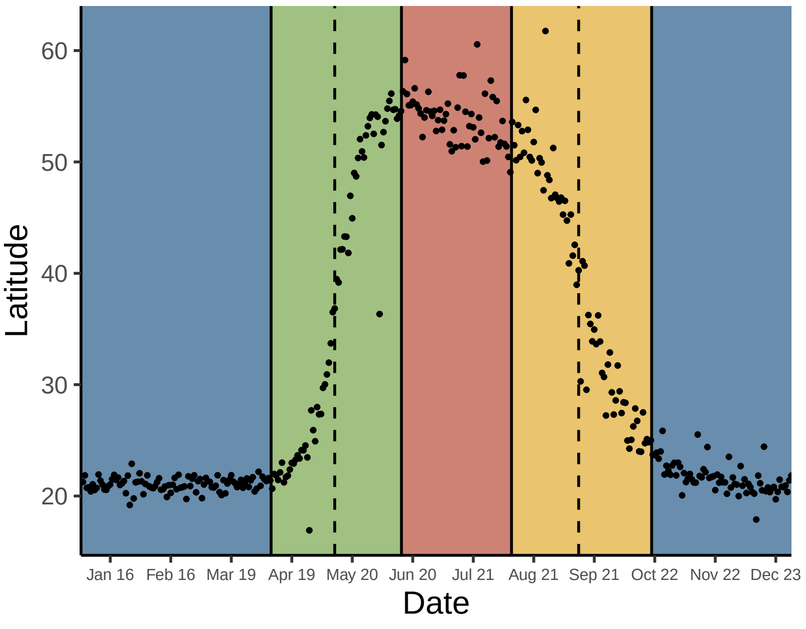

*121. Setophaga cerulea*

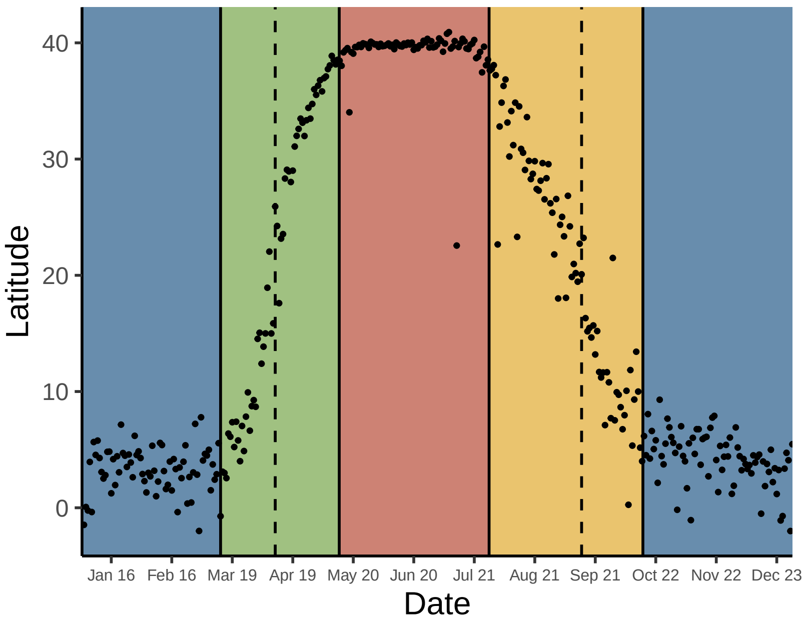


*124. Setophaga castanea*

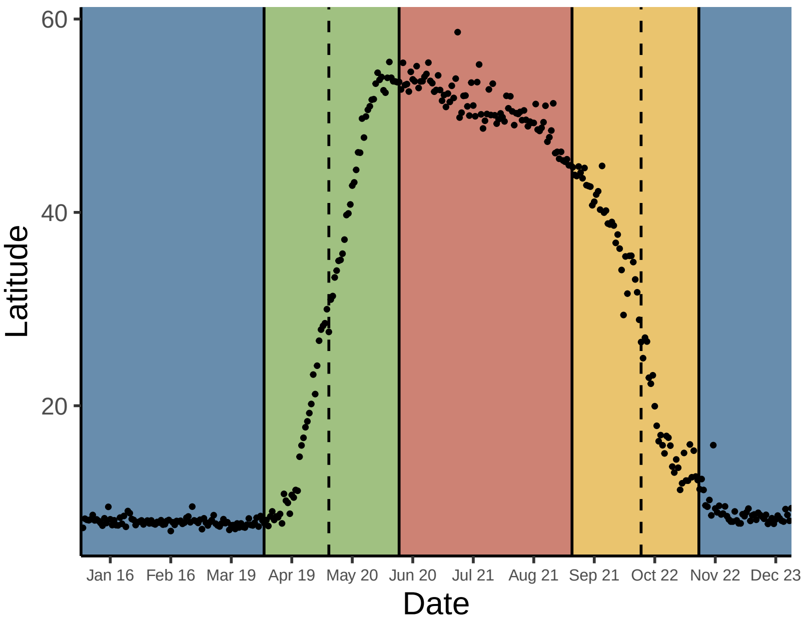


*122. Setophaga americana*

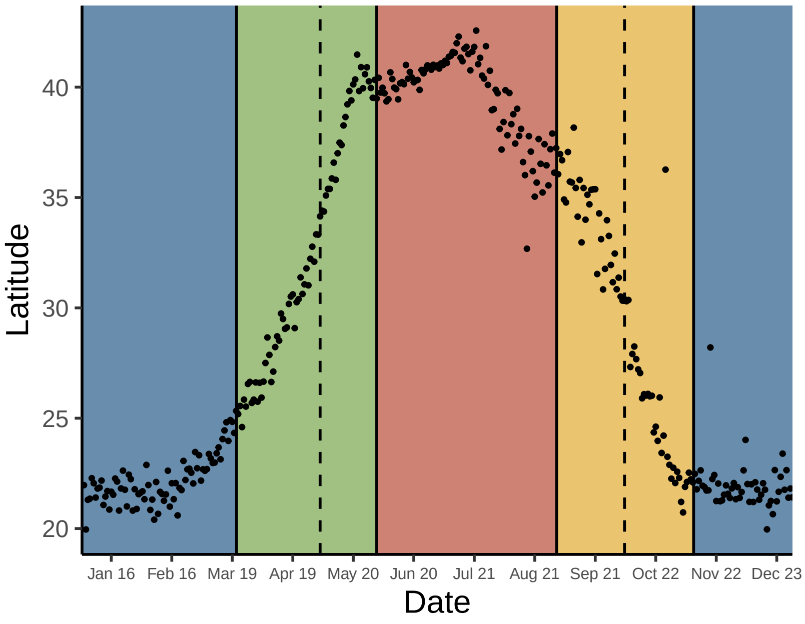


*125. Setophaga fusca*

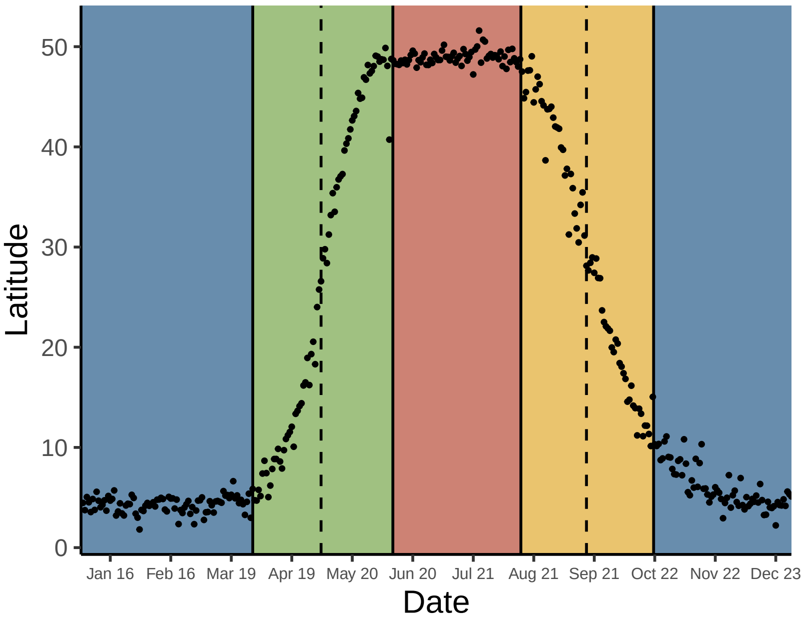


*123. Setophaga magnolia*

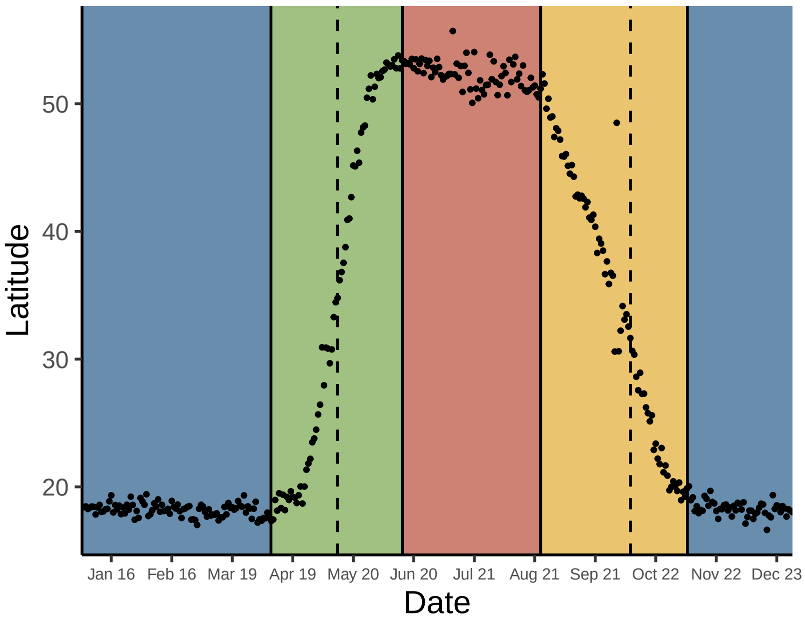


*126. Setophaga pensylvanica*

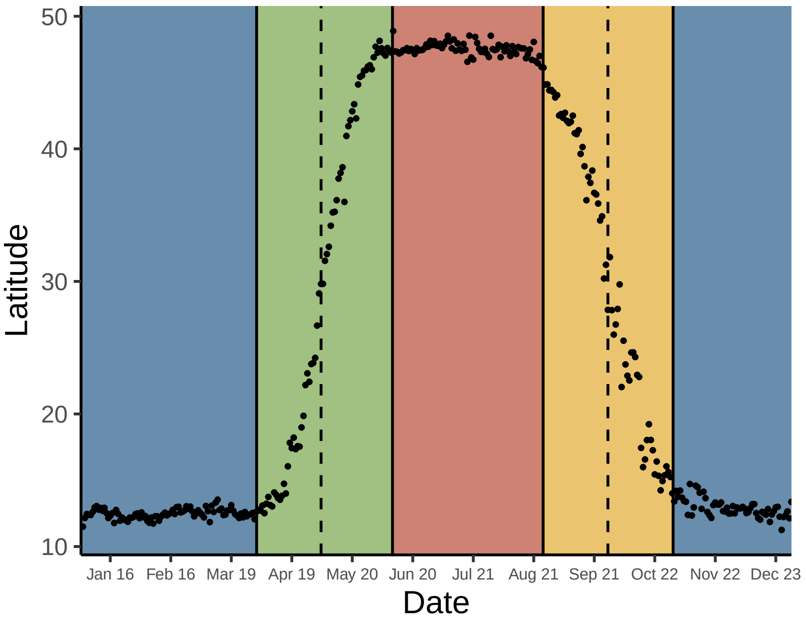

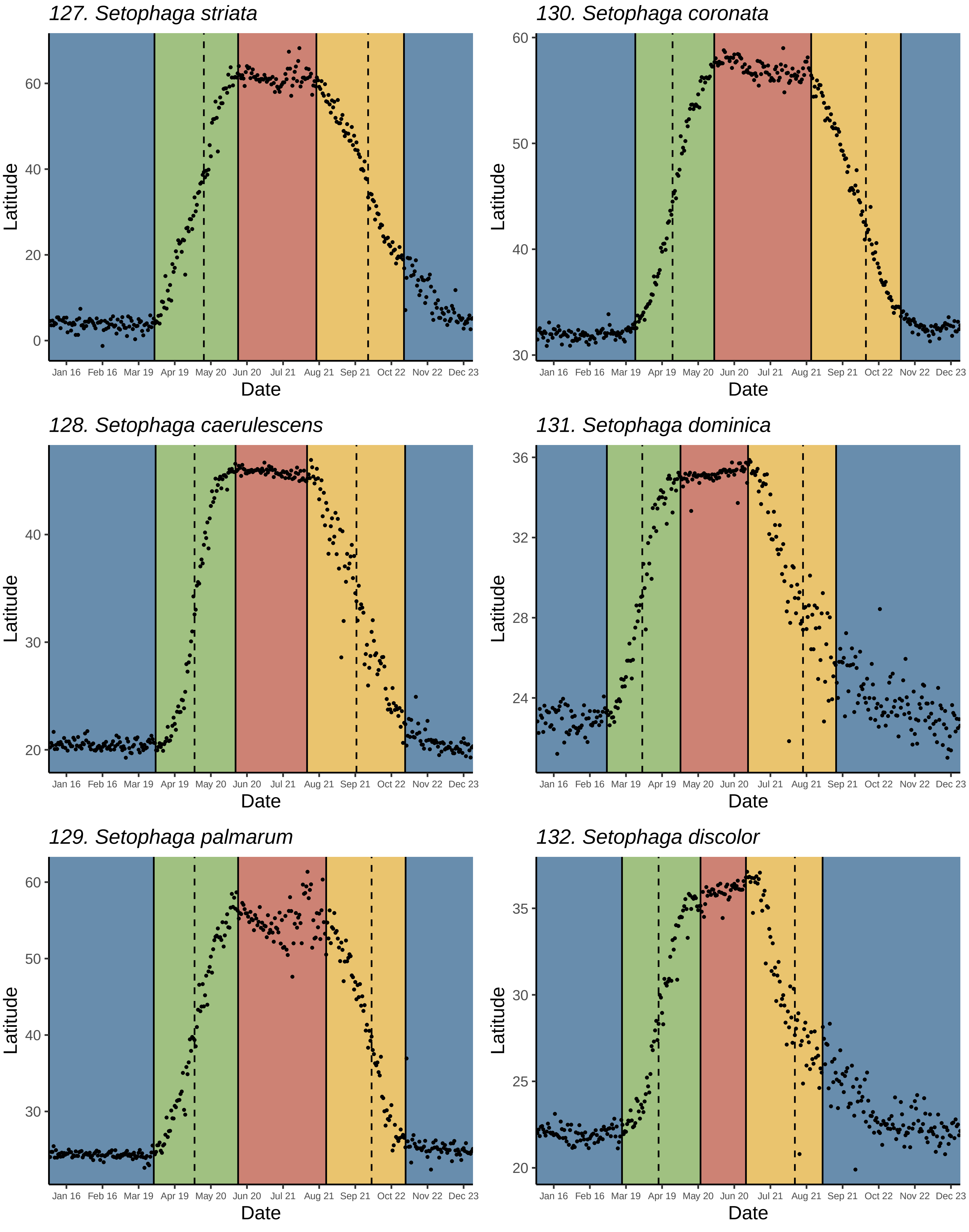
127. Setophaga striata
130. Setophaga coronata
128. Setophaga caerulescens
131. Setophaga dominica
129. Setophaga palmarum
132. Setophaga discolor
Latitude
Date
Jan 16
Feb 16
Mar 19
Apr 19
May 20
Jun 20
Jul 21
Aug 21
Sep 21
Oct 22
Nov 22
Dec 23

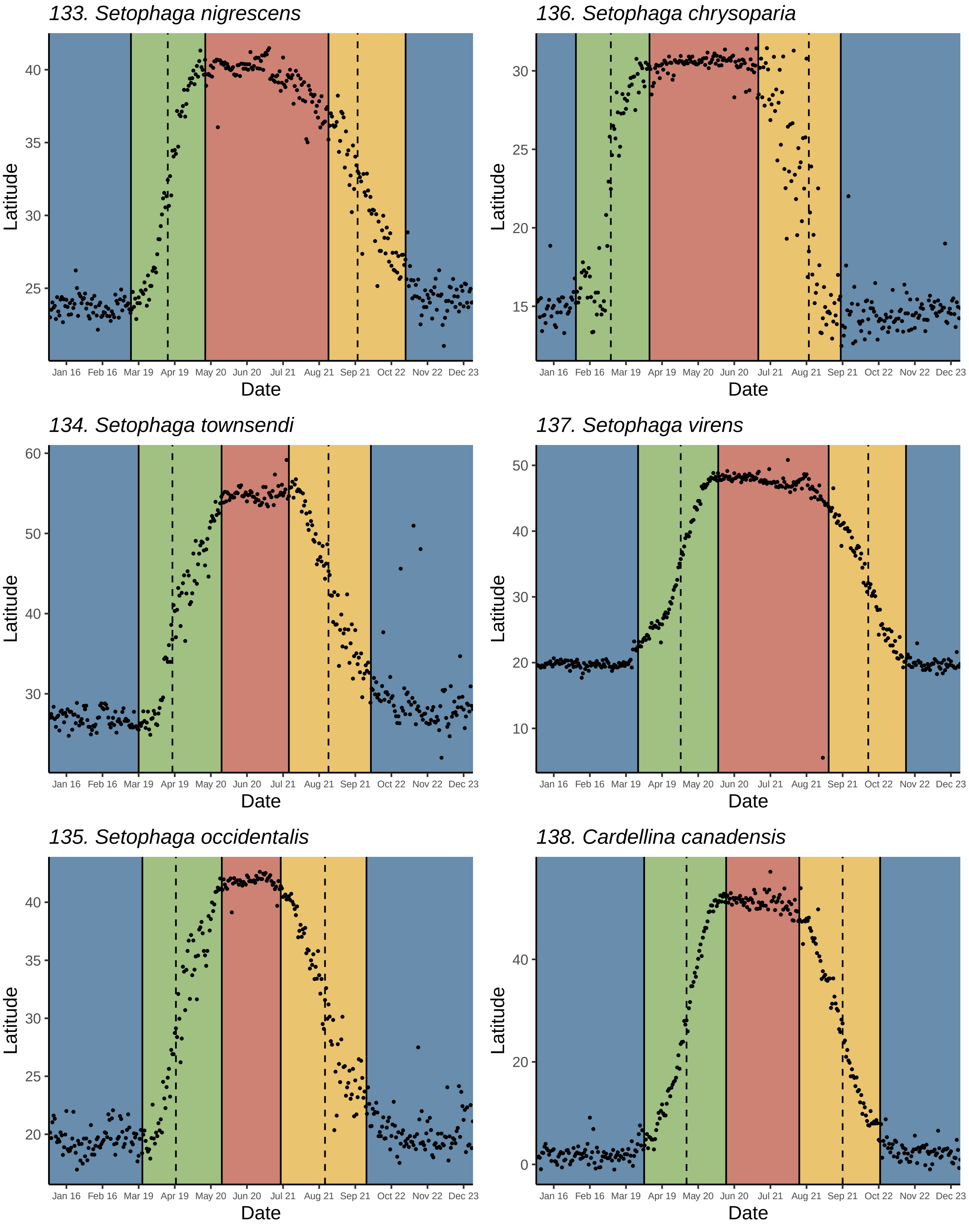
133. Setophaga nigrescens
136. Setophaga chrysoparia
134. Setophaga townsendi
137. Setophaga virens
135. Setophaga occidentalis
138. Cardellina canadensis
Latitude
Date
Jan 16
Feb 16
Mar 19
Apr 19
May 20
Jun 20
Jul 21
Aug 21
Sep 21
Oct 22
Nov 22
Dec 23

*139. Cardellina pusilla*

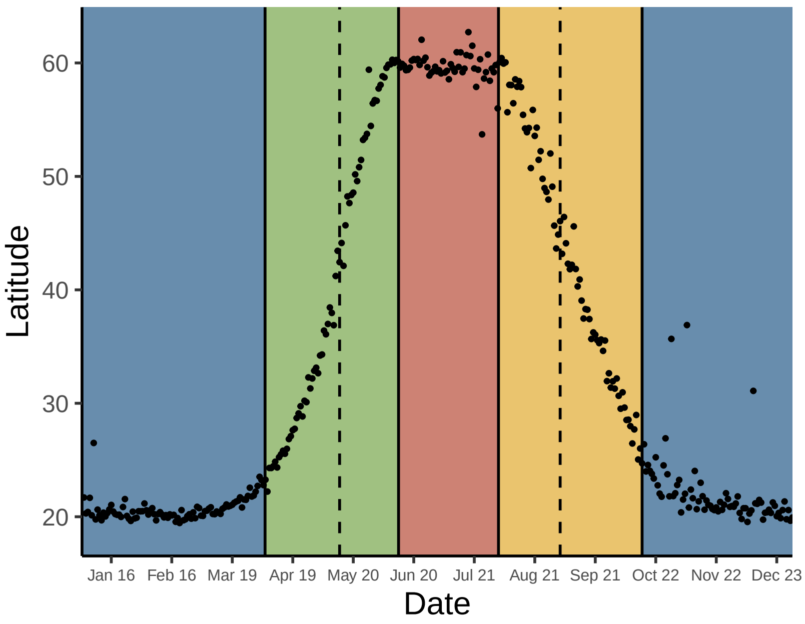


*140. Cardellina rubrifrons*

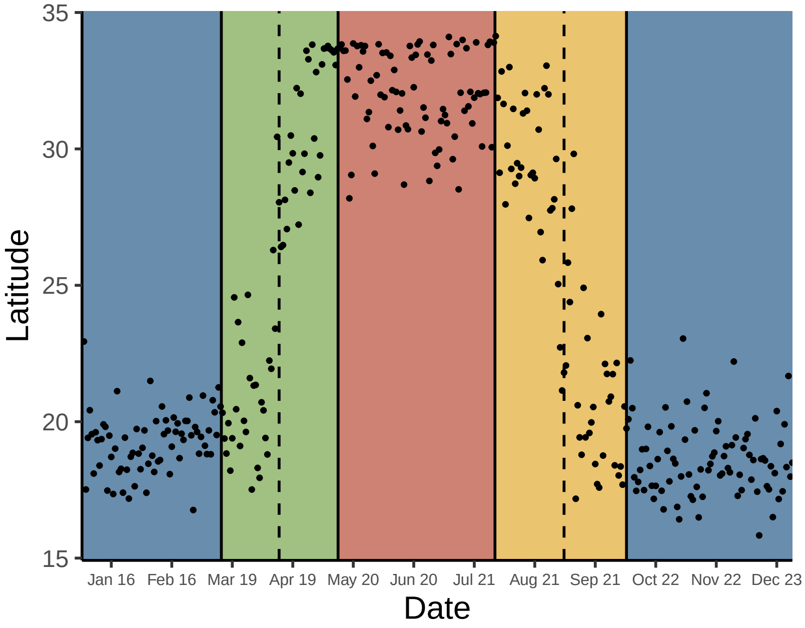


*141. Piranga rubra*

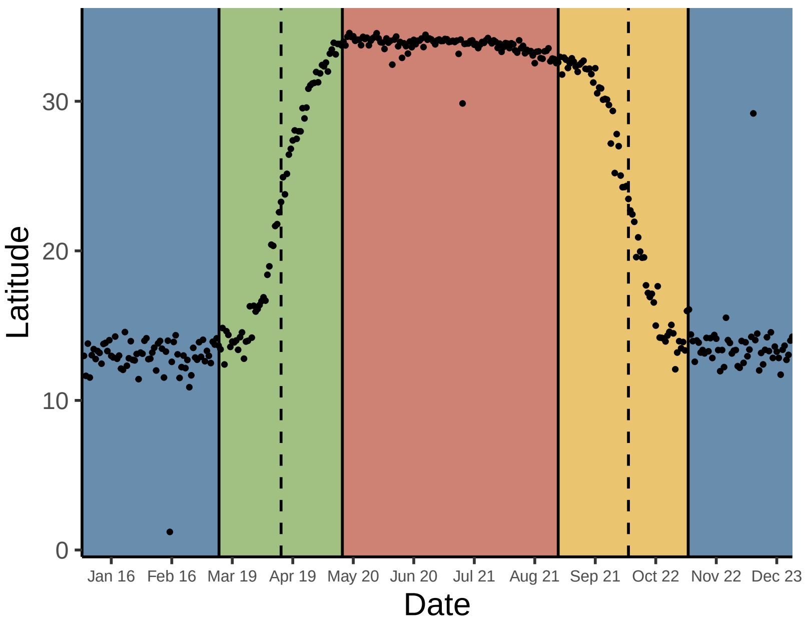


*142. Piranga olivacea*

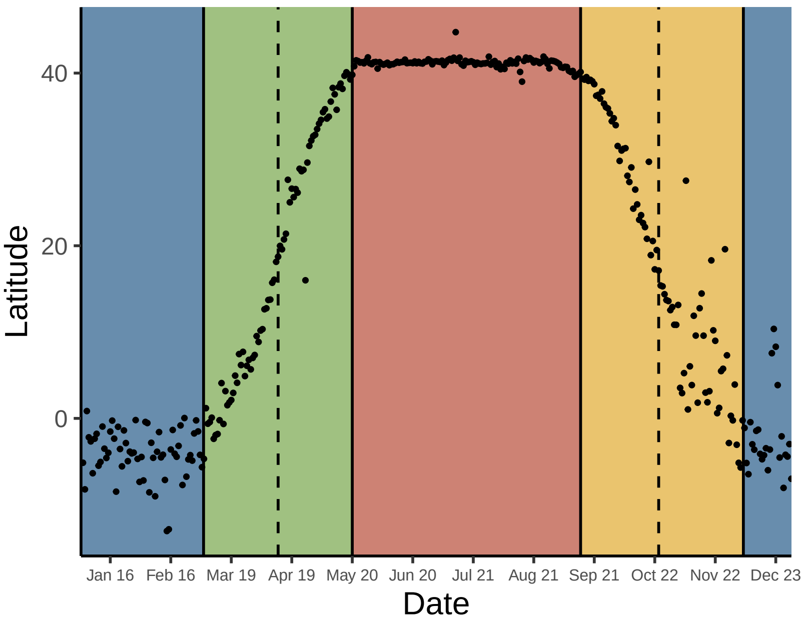


*143. Piranga ludoviciana*

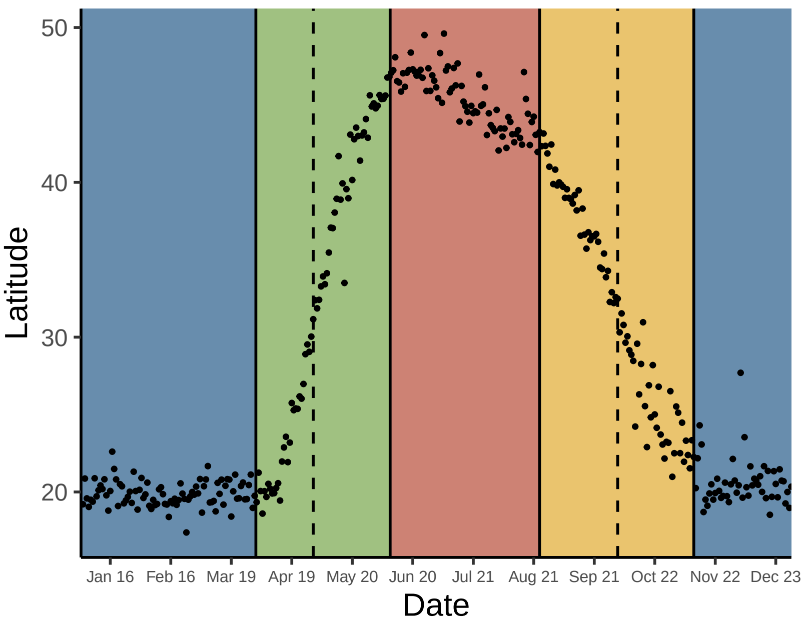


*144. Pheucticus ludovicianus*

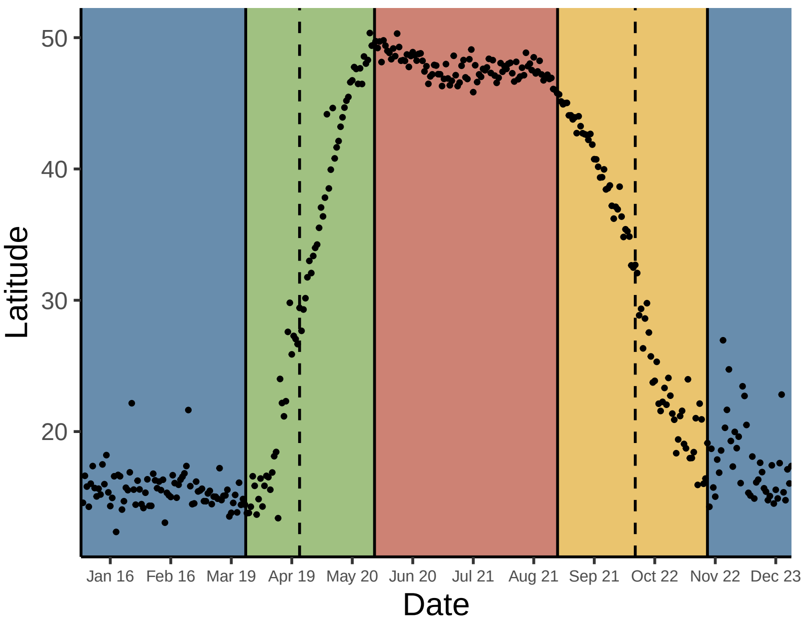

*145. Pheucticus melanocephalus*

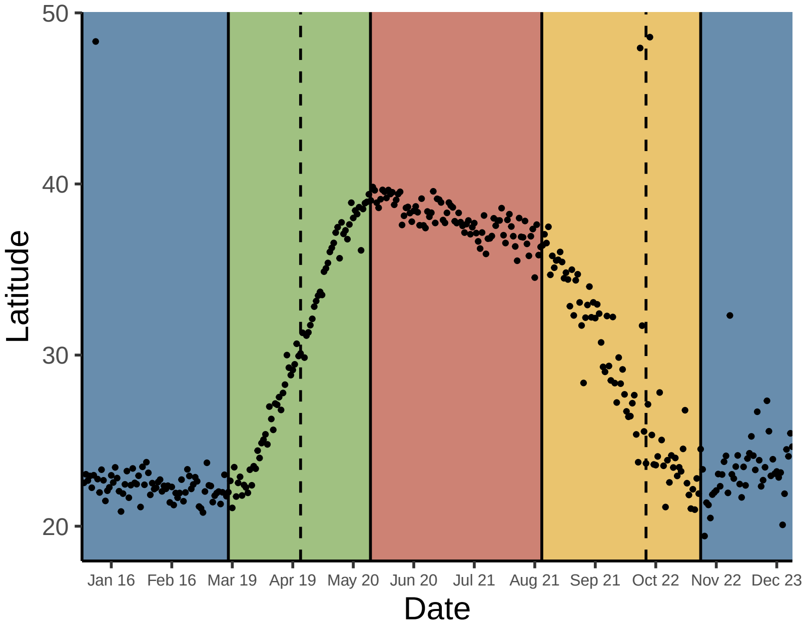


*146. Passerina caerulea*

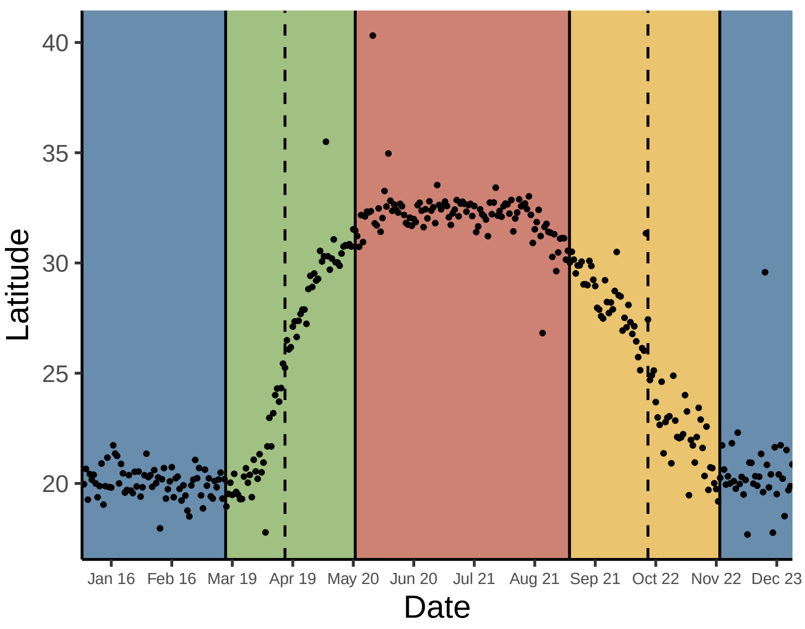


*147. Passerina amoena*

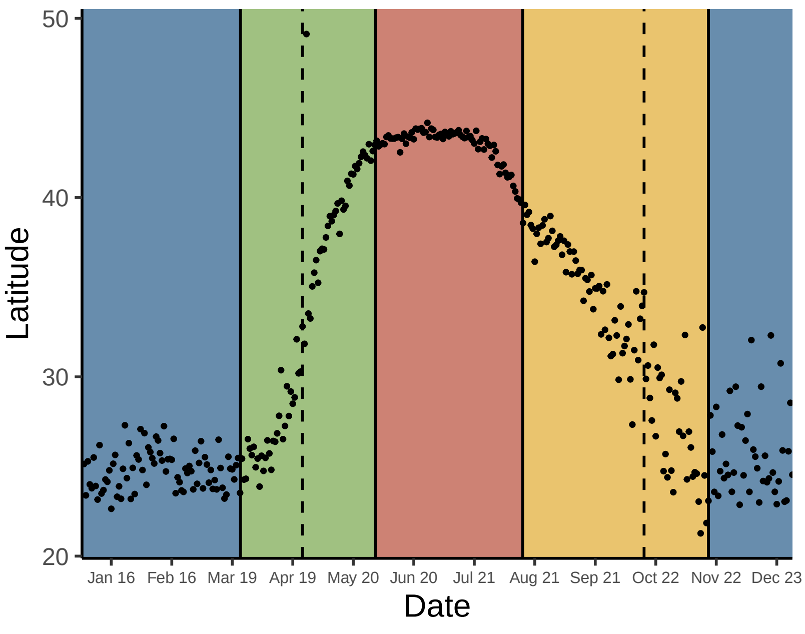


*148. Passerina cyanea*

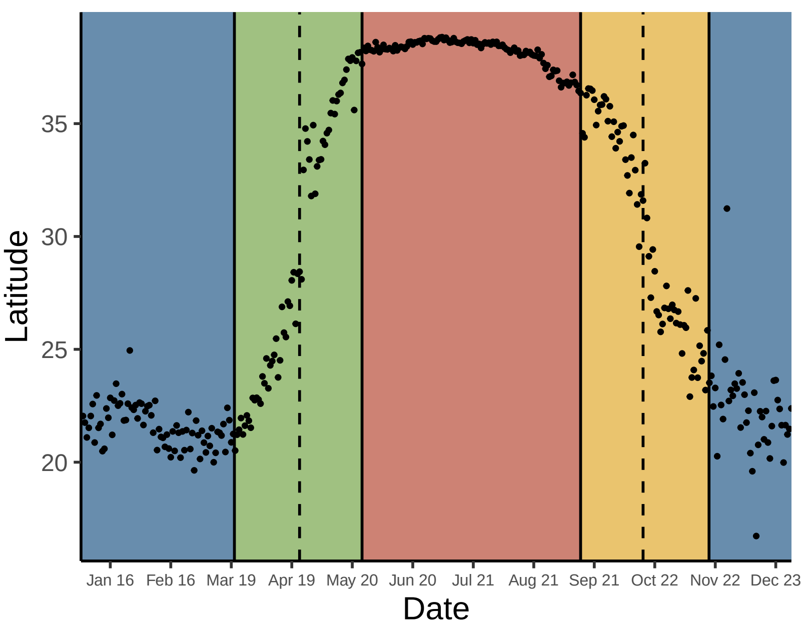


*149. Passerina versicolor*

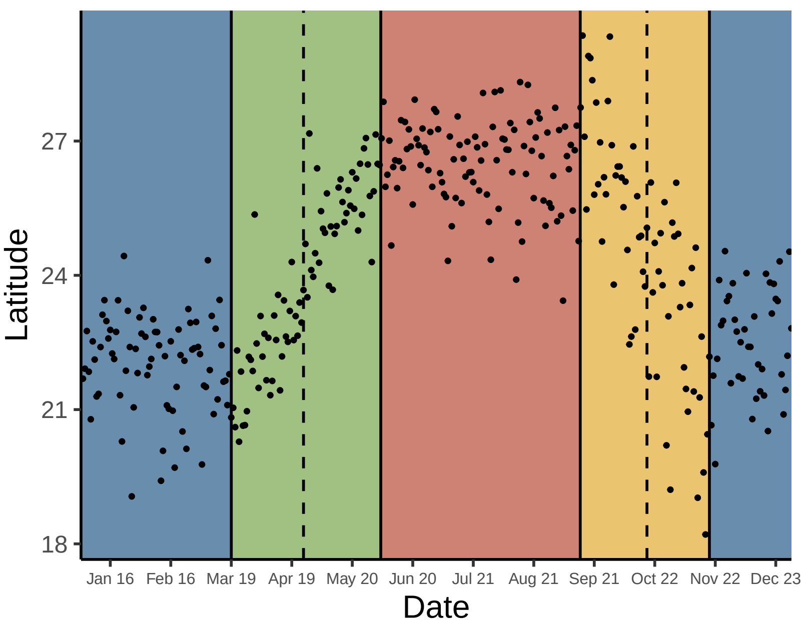


*150. Passerina ciris*

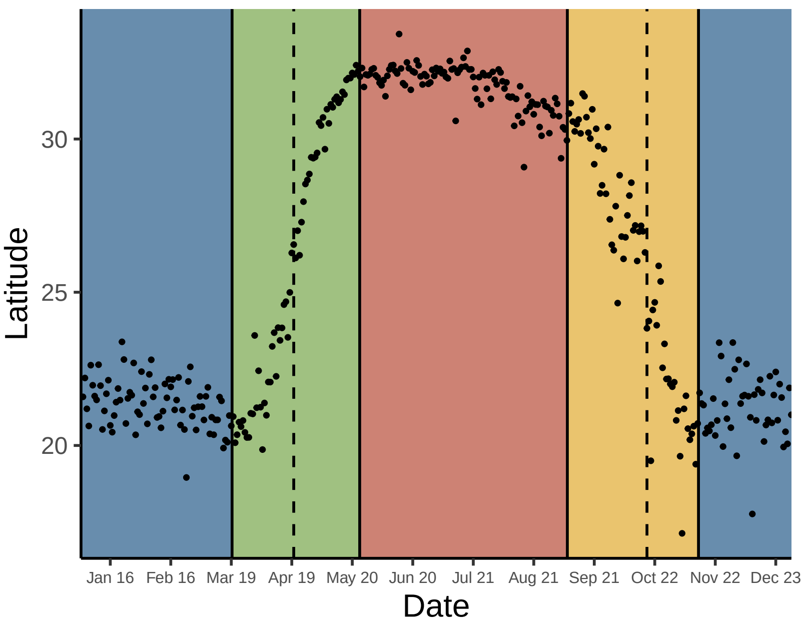
